# Protein Dynamics at Different Timescales Unlock Access to Hidden Post-Translational Modification Sites

**DOI:** 10.1101/2025.06.25.661537

**Authors:** Benjamin Lang, Julian O. Streit, Richard W. Kriwacki, John Christodoulou, M. Madan Babu

## Abstract

Post-translational modifications (PTMs) alter the proteome in response to intra- and extracellular signals, providing fundamental information processing in development, homeostasis and disease^1–5^. Here, we used proteome-wide structure predictions to investigate the structural context of protein modification sites and found nearly a fifth to be buried within proteins^6^. These cryptic sites within folded domains showed evolutionary conservation across species, as well as significantly less variability within the human population. We also found significantly more modification sites to be affected by recurrent cancer mutations and disease variants, with some variants appearing to mimic the modified or unmodified state, supporting their functionality and disease relevance. To understand how these buried sites become modified, we studied 23 phosphorylation sites across seven proteins using all-atom molecular dynamics simulations. The simulations revealed a range of mechanisms spanning broad timescales, ranging from picosecond fluctuations to seconds for proline isomerisation-gated phosphorylation sites. Notably, some sites remained fully buried in the native structure. Some conformational states with exposed cryptic sites resembled structures of co-translational protein folding intermediates formed on the ribosome^7^, indicating an interplay between protein biogenesis, folding, modifications, and signalling. We propose more broadly that protein conformational fluctuations may serve as structure-specific regulatory mechanisms by kinetically burying or exposing the unmodified, and in some cases even modified, sidechains, thereby expanding the functional versatility of proteins and enabling signal transduction across diverse timescales.

## MAIN

Protein modifications, usually referred to as post-translational modifications (PTMs), rapidly and dynamically diversify the proteome in response to both intra- and extracellular signals^5,8,9^. Examples include phosphorylation, acetylation, methylation and ubiquitination. Hundreds of modification types have been described in proteins, and high-throughput mass spectrometry experiments have led to the identification of hundreds of thousands of modification sites in humans, many of which are linked with genetic disorders and cancers^2–4^. Elucidating the mechanisms regulating the modifications at these sites is of clear biomedical importance.

Proteins are comprised of both folded domains and intrinsically disordered regions (IDRs), and it is well appreciated that many PTMs regulate protein function within conserved IDR motifs^1,10,11^. However, folded domains are frequently key to protein function, and how their functions are regulated by PTMs is less well understood. Intriguingly, evidence for modification of buried or “cryptic” sites in folded domains has emerged in recent years^12–14^. Previous studies identified buried phosphorylation sites in several proteins^6^, with these residues exposed through local conformational changes^15,16^. However, reports have generally focussed on individual proteins, rather than examining such sites systematically across the proteome and in evolution.

When modifications are introduced in buried sites within a protein’s life cycle is also often unclear. Rather than occurring strictly post-translationally, a portion has been noted to occur co-translationally, during folding or unfolding, potentially regulating these processes^17–20^. Using experimental structures, it has been suggested that buried modification sites that become solvent-exposed through conformational fluctuations appear more likely to be functional than other buried sites^21^. It has also been suggested, however, that buried sites are less likely to contribute to regulatory function and more likely to be exposed on misfolded proteins^22^. However, some of these are not degraded by the protein homeostasis machinery^23^. Meanwhile, another in-depth study found very little impact of solvent accessibility on the functionality of phosphorylation sites^24^.

To address the potential generality, consequences and mechanisms of direct regulation of folded domains by PTMs, we investigated the structural contexts and evolution of buried protein modification sites at scale using proteome-wide structure predictions from AlphaFold and AlphaSync^25,26^, and the role of protein dynamics using all-atom molecular dynamics simulations. By integrating sequence, variation, mutation, and structural data at the species, population and individual levels, we identified significant evolutionary constraints for buried modification sites that support their functional importance. In line with their functional relevance, our analyses of molecular dynamics simulations revealed a range of mechanisms exposing natively buried phosphorylation sites over a wide range of timescales. Modifications of buried PTMs may therefore be controlled by protein dynamics and folding, and it has been noted that phosphorylation can promote conformational changes without complete unfolding of folded domains^27,28^. The burial of these sidechains therefore likely enables direct modulation of function of folded domains and acts to extend signalling timescales by slowing (de)-modification kinetics through transient or long-lived sidechain burial. Our findings may thus have broad implications for disease-relevant signalling systems.

## RESULTS

### Many human protein modification sites are buried within proteins

We first analysed the solvent-accessible surface areas of known protein modification sites using structural models from the AlphaFold Protein Structure Database^25^ and our updated complementary database, AlphaSync^26^. Using relative accessibility values from AlphaSync, we applied two literature-based thresholds: residues that were ≤ 25% exposed according to their relative solvent-accessible surface area^29^ (RSA) were considered *buried*, while residues with RSA > 25% were considered surface residues^30^. Surface residues with an average RSA across a ± 10 amino acid window (RSA_10_) of < 55% were considered *structured*, while those with RSA_10_ ≥ 55% were considered intrinsically *disordered*^31^.

Strikingly, we observed that a substantial fraction of sites (17.6%) were buried within proteins (**Fig. 1a**). This finding remained consistent across different modification types (**Fig. 1b-c**), data sources and source subsets (**Fig. 1d**), as well as when filtering for structural models of the highest confidence (using AlphaFold pLDDT and PAE scores, **Extended Data Fig. 1a-k**). A subset of 12,509 sites (2%) was fully inaccessible (SASA of 0 Å²), with phosphorylation sites constituting the majority at 11,046 sites, but leaving 1,463 fully buried sites of other types.

**Figure 1.**
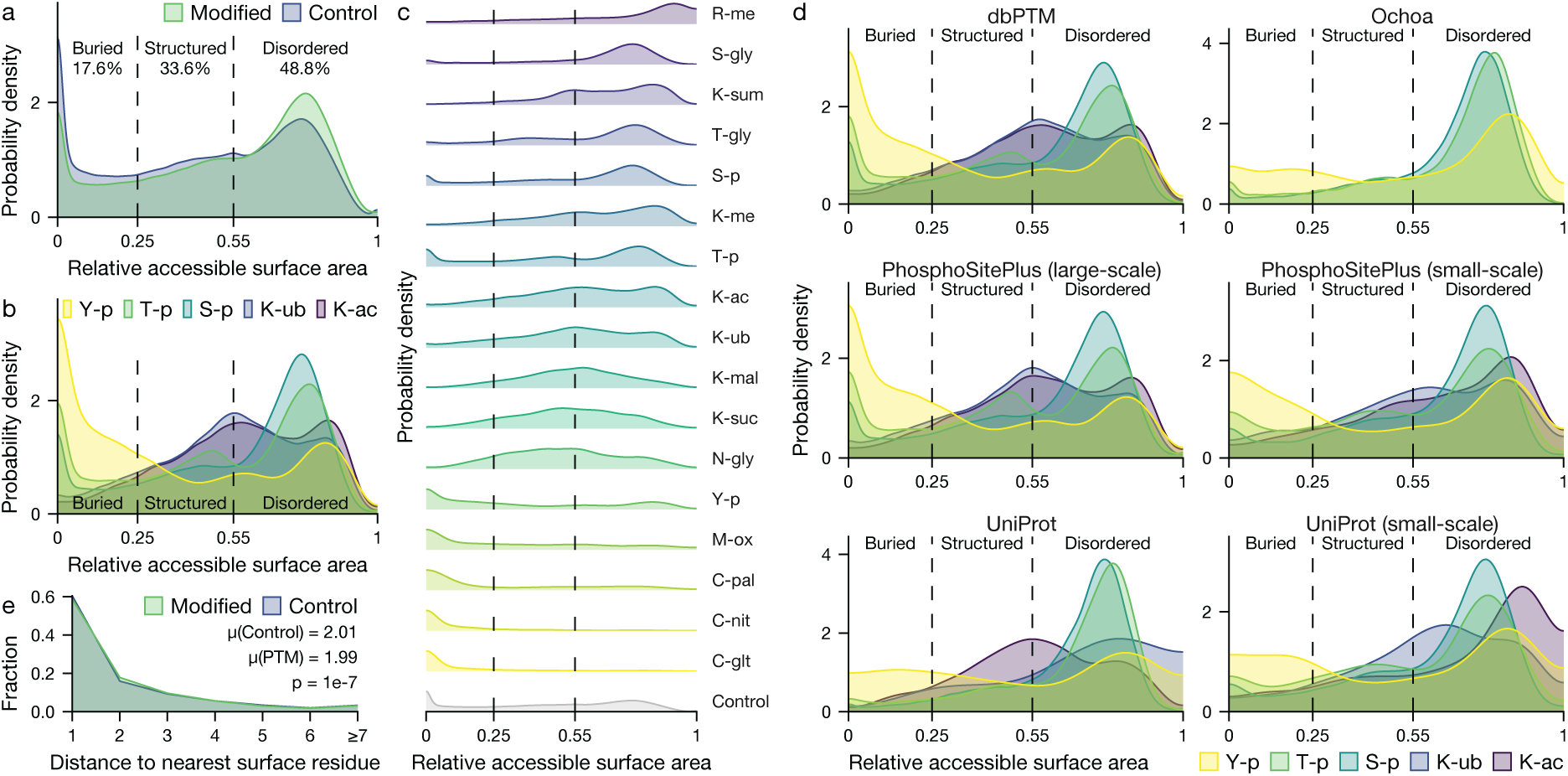
Many human protein modification sites are buried within proteins. We observed this across modification types, data sources, source subsets and structural model confidence thresholds (**Extended Data Fig. 1a-k**). Where shown, control plots were weighted to match modification site amino acid frequencies for comparability. **a**, A substantial fraction of protein modification sites is buried (17.6%), with 12,509 sites (2%) being fully buried (SASA of 0 Å^2^), compared to 26.6% and 3.3% of control residues, respectively. As expected, modified residues are more frequently found in intrinsically disordered regions overall. **b**, All major modification types have a significant fraction of buried sites, although they differ in their structural preferences. Tyrosine phosphorylation has the highest fraction of buried sites (43%), with 3.8% being fully buried, compared to 5.4% of control tyrosines. Y-p, T-p, S-p: tyrosine, threonine and serine phosphorylation; K-ub, K-ac: lysine ubiquitination and acetylation. **c**, Structural preferences for all major modification types (1,000 or more sites), arranged by descending median accessibility. While all types display some fraction of buried sites, the most abundant modifications, serine, threonine and tyrosine phosphorylation, additionally display a notable fraction of fully buried sites (SASA of 0 Å^2^). Please see Methods for a full list of abbreviations. **d**, All modification data sources and subsets showed similar and noteworthy fractions of buried modification sites for major modification types. **e**, Buried modification sites are significantly, but marginally closer to the nearest surface residue along the primary sequence than comparable control residues (in amino acids, Wilcoxon rank-sum test, W = 1.7e11, P = 1e-7).

All major modification types displayed a significant fraction of buried sites, although they differed considerably in their structural preferences (**Fig. 1b-c**). Tyrosine phosphorylation (Y-p) showed the highest proportion of buried sites at 43% (with 3.8% being fully solvent-inaccessible), compared to 54.8% of control tyrosine residues. Serine (S-p) and threonine (T-p) phosphorylation, as well as cysteine and methionine modifications, displayed particularly notable fractions of buried sites (**Fig. 1c**). We used a comprehensive set of modification data sources (**Fig. 1d**), ranging from high-confidence curated resources such as UniProt^32^ (including a subset of exclusively low-throughput studies) and systematic mass spectrometry reprocessing (Ochoa *et al.*^24^), to more inclusive sources such as PhosphoSitePlus^33^ (mass spectrometry) and dbPTM^34^ (a combined database). As trends were maintained across sources and their subsets (**Fig. 1d**), and with stringent structural confidence filtering (**Extended Data Fig. 1k**), we generally used the union of all sites in our analyses. We also noted that buried modification sites were significantly, though marginally, closer to the nearest surface residue along the primary structure than buried control residues. Note that e.g. in a β-sheet, a surface distance of one amino acid may still mean that a residue remains stably buried (**Fig. 1e**).

### Buried modification sites are evolutionarily conserved across species

By tracing the evolution of human modified residues across 199 closely and distantly related non-human species using data from the Ensembl Compara comparative genomics pipeline, 23 of them other primates (**Fig. 2a**), we were able to detect significantly lower evolutionary rates at buried modification sites relative to unmodified buried residues, indicating conservation (**Fig. 2b**). This is remarkable given the already generally high conservation of buried residues due to their importance for protein structure and thermodynamic stability and may indicate additional roles. These findings were robust across different structural filtering, conservation scoring, normalisation, alignment generation, homology handling and statistical testing methods, and they extended to structured surface and disordered modification sites as well (**Extended Data Fig. 2a-c, Supplementary Fig. 1a,c-e**). We ensured that control residues were drawn from the same set of proteins as modified ones to account for different evolutionary histories between proteins (see Methods). The choice of method to calculate the evolutionary rate presented here was made by evaluating the top-performing methods from a benchmark study^35^ on well-conserved catalytic residues from human enzymes^36^, and we saw highly similar results across methods (**Supplementary Fig. 1a-b**).

**Figure 2.**
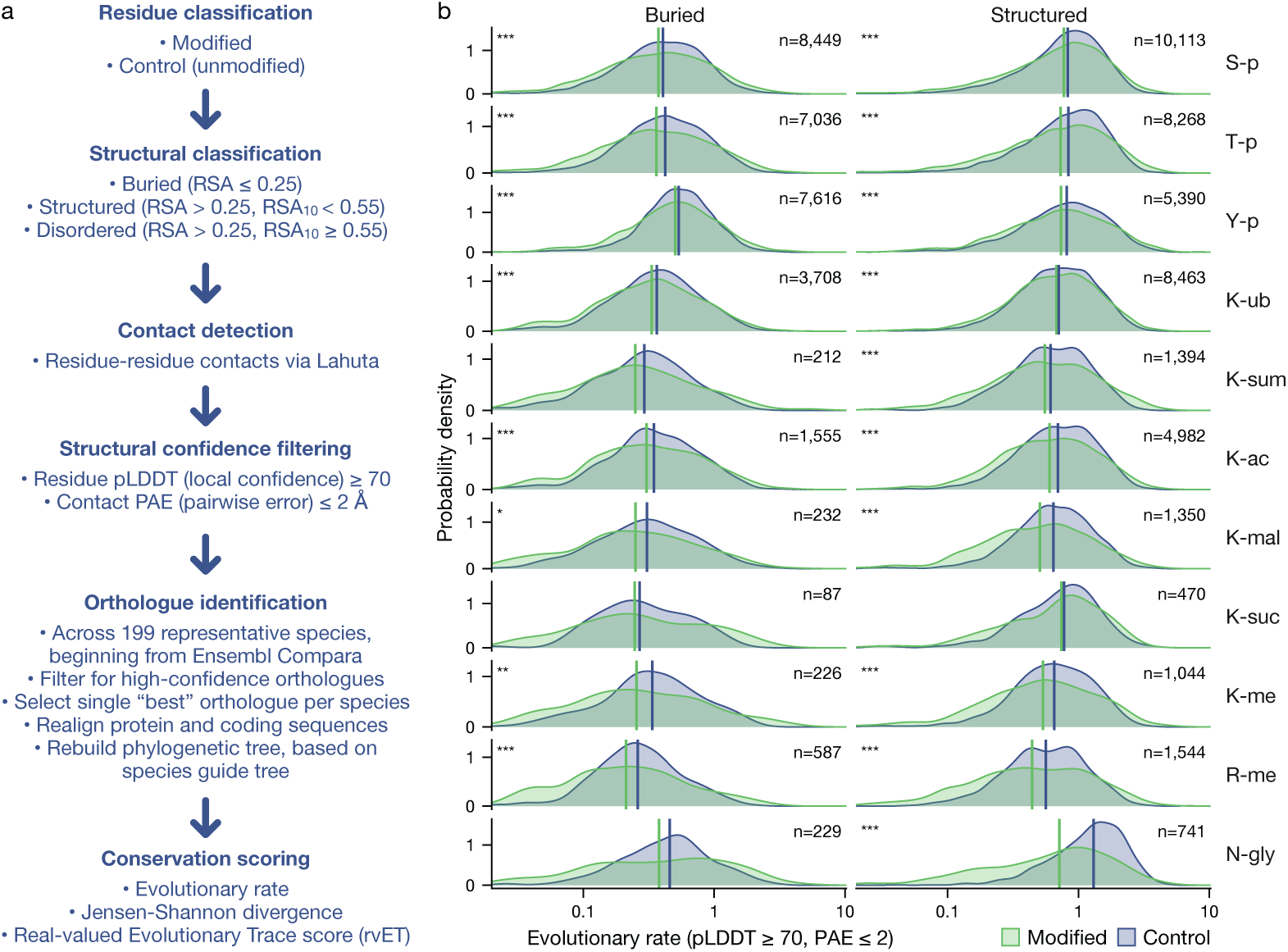
Buried residues that are modified in humans evolve significantly more slowly across species. Structured surface residues were likewise more conserved than controls. **a**, Illustration of the workflow of the evolutionary analysis presented here. **b**, Density plots show evolutionary rates for modified residues (green) compared to control residues of the same amino acid type and structural context (purple). Vertical lines show the median, and sample numbers shown refer to human proteins. For plotting, site-level evolutionary rates were averaged at the protein level, and very low evolutionary rates below 0.02 were set to this as the minimum for visualisation. Only residues with confident structural models were included (AlphaFold pLDDT score ≥ 70, and at least one non-neighbour residue contact with Predicted Aligned Error ≤ 2 Å). Proteins were included only if both modified and control residues were available for a given modification type and structural category, and all control residues were drawn from the modified set of proteins for this type and category. Significance of conservation was assessed using one-tailed asymptotic Wilcoxon-Mann-Whitney tests of sites stratified at the protein level with a minimum block size of 2 and FDR-corrected for multiple testing, and is marked using “***” for P < 1e-3 and “*” for P < 0.05. These findings were robust with different structure filtering, conservation scoring, normalisation, alignment and phylogenetic methods (**Extended Data Fig. 2a-c, Supplementary Fig. 1a,c-e**), and the choice of conservation scoring method was calibrated using well-conserved enzyme catalytic residues^36^ (**Supplementary Fig. 1a-b**).

In addition to a highly significant overall shift towards stronger conservation, modified residues in all cases displayed wider distributions of evolutionary rates than controls (**Fig. 2b**). This indicates that while a portion of buried modification sites is highly conserved (left-hand green “shoulder” in each density plot), others evolve more rapidly than comparable buried residues (right) and may be displaying relaxed constraint, lineage-specific adaptations, or recent positive selection. The existence of more rapidly evolving modification sites is expected given their roles in neo- and sub-functionalisation of proteins as shown after gene duplication^37^, and thus in speciation and phenotypic diversity more widely.

### Buried modification sites vary less in the human population, and vary more in genetic disease and cancer datasets

We next investigated the apparent evolutionary constraints on buried modification sites within the human population using non-synonymous single-nucleotide variants and mutations (**Fig. 3**). Using natural variation data from gnomAD v4^38^, we categorised variants by allele frequency and observed a clear depletion of common variants (more than 1 in 10,000 individuals) at structured surface modification sites, but not at buried sites, likely due to their strongly conserved background (**Fig. 3a**). In contrast, rare variants (1 in 10,000 individuals or fewer), which are more likely to be deleterious, showed depletion at both buried and structured modification sites (**Fig. 3b**). Overall, we noted that modified residues were significantly less likely to vary in the human population than unmodified residues (**Fig. 3a-b**, **Supplementary Fig. 2a-b,f-g, Supplementary Fig. 3a, Supplementary Fig. 5a**).

**Figure 3.**
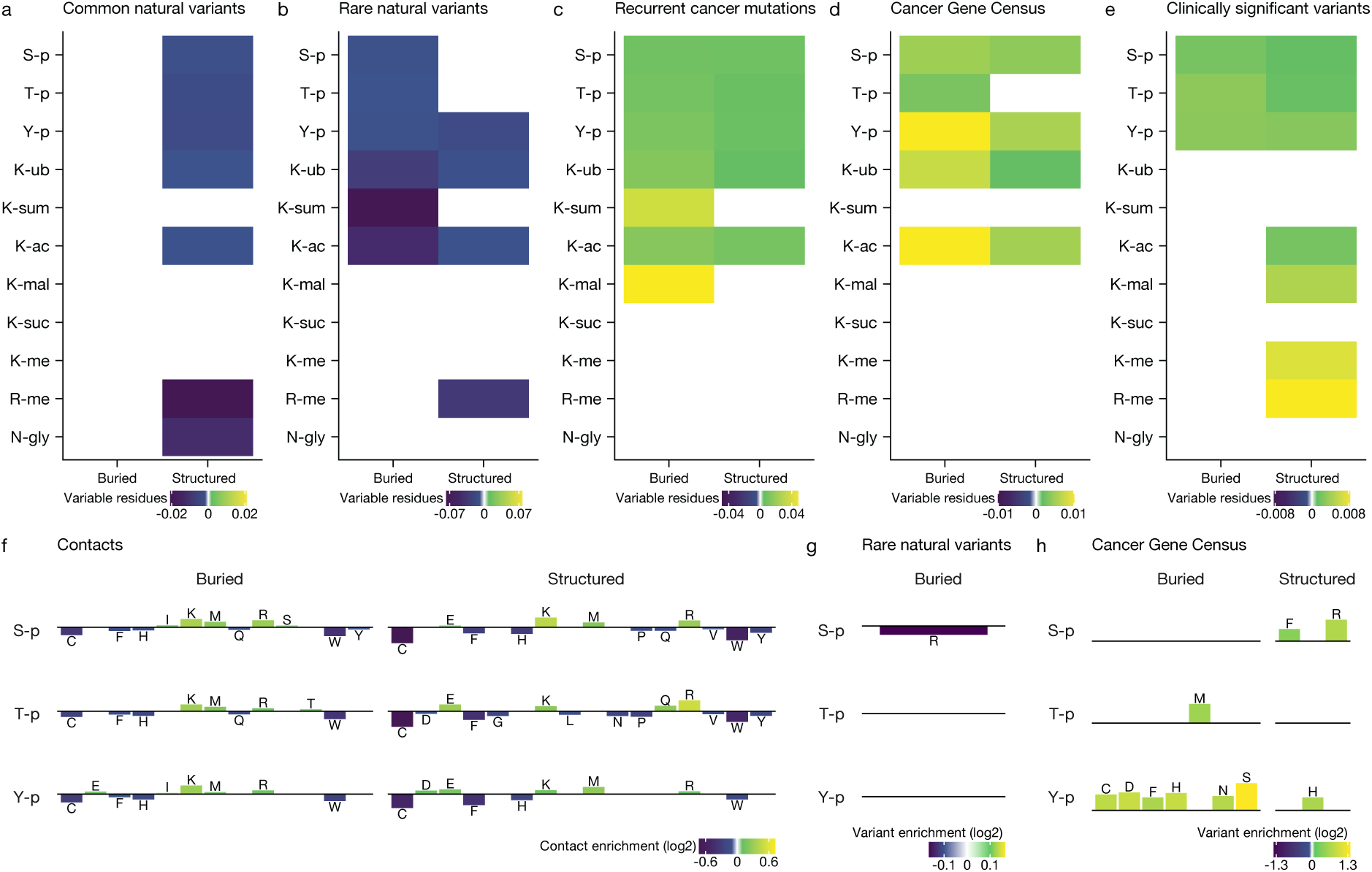
Buried residues that are modified are less often variable in the human population, and are more likely to have disease variants and cancer mutations, as well as contacts and substitutions supporting their functionality. Heat maps and bar plots showing statistically significant enrichments and depletions for modification sites compared to unmodified controls, indicated by a colour scale for each panel, with purple signifying a lower fraction of variable sites and green signifying a higher fraction. Modified and control residues were stringently filtered for structural model confidence using pLDDT ≥ 70 and PAE ≤ 2, as in **Fig. 2**. Blank panels indicate statistically insignificant differences. Residues were considered variable if they had at least one alternative allele in a given dataset. Statistical significance was assessed using Fisher’s exact test with FDR correction (P < 0.05). These findings were robust with different structural filtering methods (**Extended Data Fig. 3, Supplementary Fig. 3, Supplementary Fig. 5, Supplementary Fig. 6**). **a**, Common natural variants (more than 1 in 10,000 individuals) in the human population from gnomAD v4, showing a depletion in variability only for structured surface modification sites. **b**, Rare natural variants (1 in 10,000 individuals or fewer), showing a depletion in variability for buried as well as structured sites. **c**, Somatic mutations recurrent in cancer (≥ 10 samples) from COSMIC, showing an enrichment in variability for buried sites. **d**, Somatic mutations affecting Cancer Gene Census genes from COSMIC, a set of highly validated tumour drivers and suppressorş showing an enrichment in variability for buried sites. **e**, Genetic disorder-associated variants marked “clinically significant” in ClinVar, showing an enrichment in variability for buried phosphorylation sites. **f**, Intra-protein contacts made by modified residues compared to unmodified control residues, showing an enrichment in contacts with positively charged residues (K, R) for both buried and structured phosphorylation sites. **g**, Bar plot showing enrichment or depletion in rare natural variants (≤ 1 in 10,000 individuals). **h**, Cancer Gene Census mutations.

By contrast, when examining somatic mutations in cancer from COSMIC^39^, we found an enrichment of recurrent mutations (≥ 10 samples) at buried modification sites (**Fig. 3c**), particularly in highly validated tumour drivers and suppressors from tier 1 of the Cancer Gene Census^40^ (**Fig. 3d**). Similarly, buried phosphorylation sites were enriched in genetic disorder-associated variants marked “clinically significant” from ClinVar^41^ (**Fig. 3e**). These findings indicate that some buried modification sites may play critical roles in genetic diseases and cancers, supporting their functional significance in humans (**Fig. 3c-e**, **Supplementary Fig. 2c-e,h-j, Supplementary Fig. 3b-f, Supplementary Fig. 5b-d).**

### Contacts and mimetic variants support the functionality of buried modification sites

Led by the idea that amino acid contacts and substitutions found at buried modification sites might lend additional support to their functionality, we first investigated the contacts made by buried modification sites in their unmodified state (**Fig. 3f**, **Supplementary Table 1**) as determined using distance maxima (see Methods). This showed significant enrichment in contacts between phosphosites (doubly negatively charged when phosphorylated) and positively charged residues such as lysine (K) and arginine (R). This indicates that upon phosphorylation, these nearby positive charges might coordinate the modified residues (explored further below).

Likewise, the specific types of variants and mutations found at buried modification sites may lend additional support of functionality. We tested for enrichment and depletion of specific amino acids (**Fig. 3g-h**, **Supplementary Table 1**). In parallel with the depletion of natural variants at modification sites observed in **Fig. 3a-b**, we saw a depletion of common variants at buried sites that could be considered highly disruptive compared to the modified state: for example, substituting nonpolar alanine (A) for doubly negative phosphothreonine (T-p), or to a lesser extent polar asparagine (N) for phosphoserine (S-p) (**Supplementary Fig. 4a**). For rare variants, positive arginine (R) was likewise depleted at buried phosphoserines (S-p) (**Fig. 3g**), and negative glutamate at lysine acetylation sites (**Supplementary Fig. 4b**). While depletion in specific amino acids is difficult to interpret, it is nonetheless an important finding that may be indicative of the structural context and effects of buried modification sites.

We also observed statistically significant variant enrichments (**Fig. 3h, Supplementary Fig. 4b,d**) that make sense in light of the modified state, for example in rare variants at arginine methylation (R-me) sites, which were enriched in nonpolar methionine (M) (**Supplementary Fig. 4b**), potentially mimicking methylarginine^42^. Most interestingly, we observed COSMIC Cancer Gene Census somatic mutations to negative aspartic acid (D) at buried tyrosine phosphorylation (Y-p) sites (**Fig. 3h**). Though sterically much smaller, aspartate’s charge can make it a successful phosphomimic, mirroring the modified state (see Discussion). Notably, other cancer enrichments for these buried phosphotyrosines included aromatic phenylalanine (F), polar serine (S), cysteine (C), asparagine (N) and potentially uncharged histidine (H) (**Fig. 3h**), all of which might act as potential mimics of the unmodified state. Cysteine might alternatively introduce disulfide bonding, with potentially deleterious effects^43^.

Other enrichments for genetic disease variants and cancer mutations appeared highly disruptive, such as substituting nonpolar isoleucine (I) for buried phosphothreonine (T-p) in ClinVar’s clinically significant variants (**Supplementary Fig. 4a**), and substituting nonpolar methionine (M) for the same in Cancer Gene Census genes (**Supplementary Fig. 4d**). For structured (surface) residues, enrichments were likewise disruptive, e.g. to the opposite charge (E) for lysine acetylation (K-ac) and ubiquitination (K-ub) sites (**Supplementary Fig. 4d**). However, an enrichment for polar glutamine (Q) at lysine acetylation (K-ac) sites in the Cancer Gene Census could potentially be a functional acetyllysine mimic^44^ (**Supplementary Fig. 4d**). As the Census is composed of both oncogenes and tumour suppressors, both overactivation and disruption of protein function may lead to tumour formation.

Though the overall variability enrichments and depletions we observed were of modest magnitudes (**Fig. 3a-e**), the contact (**Fig. 3f**) and variant (**Fig. 3g-h**) enrichments and depletions further added support for the functional importance of buried modification sites (**Supplementary Fig. 5e-f, Supplementary Fig. 6a-d**).

### A wide range of protein dynamics regulate accessibility to buried sites

The above analyses allowed us to identify promising buried modification sites which might become accessible for modification. We focussed on candidates with the following features: i) buried, ii), close to solvent-exposed residues (**Fig. 1e**), iii) confident AlphaFold prediction of the region, iv), availability of an experimental structure, v) a structured domain of reasonable size for simulation, vi) only rare genetic variants and vii) reasonable conservation across species (**Supplementary Table 2**, Methods).

To explore possible mechanisms, we used all-atom molecular dynamics (MD) simulations (see Methods). We focused on phosphorylation sites since phosphorylation is expected to be associated with solvent exposure due to the highly polar and charged nature of the modification, likely less compatible with sidechain burial, though some phosphorylation sites are thought to reconstitute ancestral salt bridges^45^. Simulations have previously been used to investigate local folding and unfolding mechanisms coupled to exposure of buried phosphotyrosine residues in p27^15^ and Vav1^16^. These studies showed that buried phosphorylation sites can be transiently exposed in the unphosphorylated state through local protein dynamics, in line with prior experimental studies^46–48^, and providing a mechanistic basis for their modification by kinases. Building on these observations and our list of buried candidate sites, we investigated 7 other proteins containing buried phosphorylation sites in their unphosphorylated or phosphorylated states (**Fig. 4, Extended Data Figs. 3-10**). We will first provide a general overview of the types of mechanisms revealed by the simulations before describing the proteins individually.

**Figure 4.**
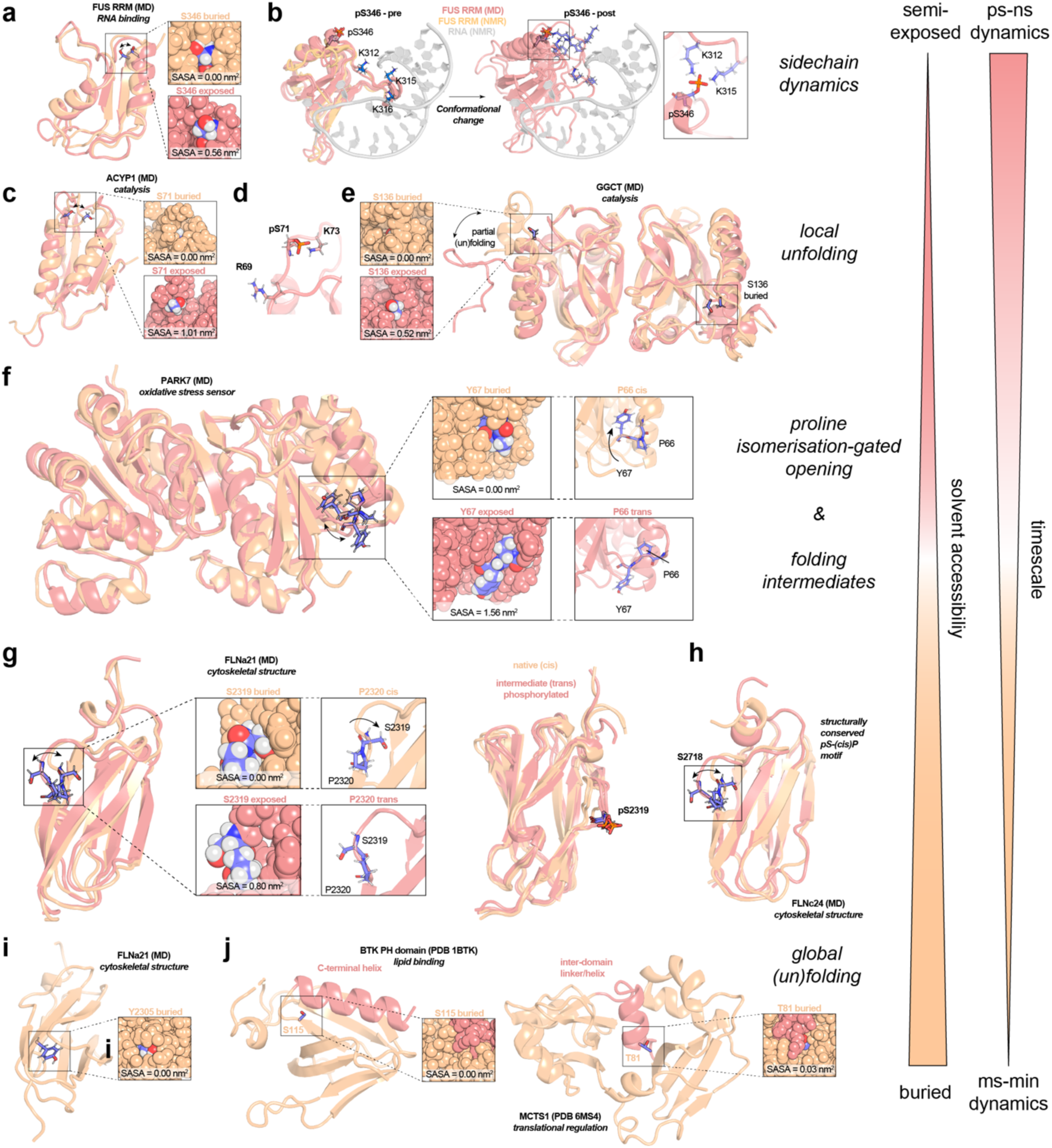
Protein dynamics and conformational changes regulate exposure of cryptic phosphorylation sites as observed in all-atom molecular dynamics (MD) simulations. **a**, S346 in the FUS RNA recognition motif (RRM) domain exhibits sidechain dynamics on the pico- to nanosecond timescale to alternate between buried and exposed conformations. **b**, Phosphorylation of FUS RRM (pS346) results in a conformational change due to contacts between pS346 and K312, K315, and K316. These positively charged residues are responsible for RNA binding via electrostatic interactions with the phosphate backbone (PDB 6GBM^54^). **c**, A disordered loop including S71 in ACYP1 transitions between an exposed and buried S71 conformation on the nanosecond timescale. **d**, Phosphorylated pS71 makes contacts with positively charged sidechains in the structure. **e**, The C-terminus of GGCT (homodimer) is dynamic and partial unfolding in this region leaves S136 exposed to the solvent. **f**, Y67 in the PARK7 homodimer transitions between a buried and exposed conformation, gated by the isomerisation state of the upstream proline P66. A *cis* P66 conformation favours Y67 to be buried, while *trans* P66 promotes opening of the structure around Y67. **g**, Proline isomerisation in human filamin immunoglobulin (Ig)-like domains regulate exposure of upstream serine residues S2319 (FLNa21). A *cis* proline favours a buried serine residue, while isomerisation to the *trans* conformation results in serine exposure to the solvent. The *trans* proline conformation has been shown to correspond to a kinetic folding intermediate^50,58^. (right) Superimposed MD structures of the native (*cis*) and trapped intermediated (*trans*, phosphorylated). **h,** An identical proline-regulated switch is observed for S2718/P2719 in FLNc24 at the conserved pS-(cis)P motif. **i**, Buried Y2305 remains solvent-inaccessible in the core of FLNa21 in all-atom MD simulations. **j**, Example experimental structures of proteins with PTM sites buried in the core of the protein, including tyrosine-protein kinase BTK (PDB 1BTK^82^) and malignant T-cell-amplified sequence 1 (MCTS1, PDB 6MS4^83^). Note that these proteins were not simulated.

In agreement with the experimental evidence of these buried PTM sites becoming modified (**Supplementary Table 2**), we have successfully identified exposed conformational states for most proteins and buried PTM sites considered here (**Fig. 4, Extended Data Figs. 3-10**). The simulations suggest a broad range of exposure mechanisms spanning multiple orders of magnitude in terms of their dynamic timescale. We observed several phosphorylation sites to be dynamically accessible through pico- to microsecond-scale sidechain and backbone fluctuations (**Fig. 4a-c, Extended Data Figs. 3-5 & 9a-f**) and local unfolding of the native protein structure (**Fig. 4c-e, Extended Data Fig. 5-6**). In other cases, we observed the accessibility of buried phosphorylation sites to be gated by proline isomerisation (**Fig. 4f-h, Extended Data Figs. 7-9**), suggesting kinetic control over phosphorylation and signalling dynamics due to the slow process of isomerisation (seconds to minutes^49,50^). Lastly, some highly buried residues in the hydrophobic core of the protein are expected to remain inaccessible to modification through local conformational changes (**Fig. 4i-j, Extended Data Figs. 7q-r, 8o-p**), which indicates their modification may occur prior to protein folding or after unfolding. Overall, these insights highlight the importance of protein dynamics in protein modifications and suggest diverse mechanisms for enabling slow phosphorylation through “kinetic shielding” of phosphosites in their unphosphorylated states.

Moreover, simulations of the phosphorylated states of these proteins showed functionally relevant local and global conformational changes (**Extended Data Figs. 3-10** and see *below*). We also noted that although most phosphorylated sidechains remained solvent-accessible, some examples also exhibited significant burial through shielding of the modified residues by contacts from positively charged sidechains (*see below*, **Supplementary Fig. 7, Fig. 3f**). These structural features may consequently act to also slow down removal of the modification by phosphatases, and enable signalling responses to occur on longer timescales compared to rapidly accessible phosphosites in disordered regions.

### Fast sidechain and backbone dynamics modulate accessibility

We reasoned that phosphorylation sites that are solvent-inaccessible, but located near disordered loop regions and the protein surface, may become exposed through local protein dynamics such as sidechain rearrangements and backbone fluctuations. To investigate this, we simulated the RNA recognition motifs (RRM domains) of the human RNA-binding proteins FUS^51^ and SRSF2^52^, and the Parkinson’s disease-related protein homodimer PARK7^53^. The FUS domain is observed to be stable in our simulations and the average solvent accessibility of cryptic phosphorylation sites T286, S340 and S346 was indeed lower than for other threonine and serine residues (**Extended Data Fig. 3a-h**). However, their solvent-accessible surface areas (SASA) fluctuated greatly on the pico- to nanosecond timescale (**Extended Data Fig. 3d-h**). These fluctuations reflect the fast dynamics of these sidechains, as the backbone fluctuations in these regions are stable (**Fig. 4a**, **Extended Data Fig. 3a-b**). Importantly, extensive simulations of the singly phosphorylated variants revealed insights into the potential functional consequences of these phosphorylation events. Most notably, FUS pS346 underwent a highly reproducible conformational change driven by electrostatic interactions between the phosphate group and positively charged K312, K315 and K316 (**Fig. 4b**). The resulting conformational state appears to be less likely to bind to RNA because the loop containing residues 312–316 forms electrostatic interactions with the phosphate backbone and provides shape complementarity by fitting into the RNA major groove^54^ (**Fig. 4b**). In line with this, K315A and K316A mutations and lysine acetylation of these residues have been shown to reduce binding affinity^51,54^. Phosphorylation of T286 and S340 is also likely to interfere with RNA binding since these residues are located at the RNA binding interface, promote competitive interactions with sidechains of K312, K315, and R371 (all implicated in RNA binding^54^), destabilise the FUS RRM structure locally, result in steric clashes with the RNA, and likely electrostatically repel the domain from the RNA backbone (**Extended Data Fig. 3i-p**). Sequence motif analyses suggested potential modification of T286 by PLK1, PLK4 or GSK3 and of S340 by PLK1^55^, and the apparent modulation of FUS RNA binding affinity inferred from these structural changes suggests that these cryptic sites are indeed functional.

We also investigated three phosphorylation sites in a similar RRM domain of SRSF2 (T14, T22, and T25), which also showed large SASA fluctuations in our simulations ranging from completely buried to fully solvent-accessible (**Extended Data Fig. 4a-g**). Sequence motif analyses suggested possible phosphorylation of T22 and T25 by GSK3 or PLK1, and of T25 by proline-directed (MAPK) kinases^55^. In all cases we found that local sidechain rearrangements on the surface and backbone fluctuations resulted in different conformational states that either bury or expose these residues. We then also simulated the individual phosphorylated variants, and similar to FUS, observed electrostatic interactions between the phosphate groups and positively charged residues, albeit without inducing larger conformational changes (**Extended Data Fig. 4h-q**). These positively charged residues include sidechains that make direct contact with RNA (e.g. R49 and R94)^56^, thus also suggesting that phosphorylation of these threonine residues may modulate RNA binding affinity.

To consider a larger complex, we studied the Parkinson disease protein PARK7 (also called DJ-1). PARK7 is a homodimer and involved in protection from oxidation-related stress^53^. While the dimer structure was observed to be highly stable in our simulations, two phosphosites near the surface and dimer interface, S47 and S57, displayed fluctuations between buried and partially exposed conformations due to local sidechain dynamics, potentially rationalising their modification (**Extended Data Fig. 9a-f**). Extensive simulations of the phosphorylated variants showed a pS47-specific effect on the dimer structure, whereas pS57 remained essentially unchanged with respect to structure and dynamics (**Extended Data Fig. 10a-g**). Specifically, pS47 resulted in a loss of contacts at the dimer interface (increased dimer interface RMSD), suggesting that phosphorylation at this site may destabilise the dimer. Overall, this suggests that cryptic sites near the protein surface can become temporarily accessible through local sidechain and backbone dynamics as observed for FUS, SRSF2, and PARK7. The effects of phosphorylation on the structure of these proteins are indicative of the functional relevance of these modifications.

Interestingly, the sidechain solvent accessibility of the phosphorylated variants of FUS, SRSF2 and PARK7 varied substantially due to sidechain contacts with positively charged residues described above. For example, FUS pS340 and SRSF2 pT14 displayed a low average accessibility and fluctuated between completely buried and exposed conformations, while the others, in particular pT25, remained at least partially solvent-accessible (**Supplementary Fig. 7**).

### Local (un)folding of protein structure to expose buried phosphorylation sites

In two other proteins, we observed accessibility to cryptic phosphorylation sites to be determined by local conformational changes. Human acylphosphatase-1 (ACYP1) exhibited local unfolding of the last helical turn in helix 2 during 1 μs-long unbiased simulations, and the resulting conformational change rotates S71 from a buried to accessible state (**Fig. 4c, Extended Data Fig. 5a-e**). Upon phosphorylation of S71, the increased dynamics surrounding this site are maintained locally but no further large-scale conformational changes were observed on the timescale of our simulations (5 μs), including the active site which remained in a native-like conformation (**Extended Data Fig. 5f-j**). Similar to FUS and SRSF2, however, we also observed electrostatic contacts between the phosphate group of pS71 and nearby positively charged residues (R69 and K73, **Fig. 4c, Extended Data Fig. 5a-e**), in line with the evolutionary hypothesis of phosphorylation establishing switchable salt bridges in protein structures^45^. In this case, pS71 remained completely solvent-accessible after modification (**Supplementary Fig. 7**).

A larger, local unfolding conformational change was observed for human gamma-glutamylcyclotransferase (GGCT), which is a homodimer involved in glutathione homeostasis^57^. In multiple unbiased simulations lasting 1 μs, we observed local unfolding of the C-terminal tail and helix of the homodimer, which otherwise render the phosphosite S136 completely solvent-inaccessible (**Fig. 4e, Extended Data Fig. 6**). This highly dynamic behaviour is qualitatively consistent with the high temperature factors observed for the C-terminus in the crystal structure (**Extended Data Fig. 6b**) and suggests that GGCT exists in two conformational states in solution (**Extended Data Fig. 6c,h**), but we have not attempted to predict the underlying free energy landscape. Instead, we simulated the GGCT dimer with S136 phosphorylated on one chain (pS136), and the C-terminus remained unstructured on the timescale of these simulations (4 μs) while the dimer interface was slightly destabilised (**Extended Data Fig. 6h**). We propose therefore that phosphorylation may shift the equilibrium to the C-terminally unfolded state and may affect protein dimerisation. The phosphate group of pS136 was seen to dynamically interact with positively charged residues in the C-terminal tail (mainly K171) and form hydrogen bonds with H65 and G66 (**Extended Data Fig. 6g-m**). This further suggests that in many cases, these exposed phosphorylated residues are at least partially coordinated by polar and positively charged functional groups that are close in the structure, in this case resulting in transitions between states where pS136 is completely buried and solvent-accessible on the microsecond timescale (**Supplementary Fig. 7**).

### Proline isomerisation can act as a gatekeeper for solvent accessibility

We noticed some phosphosites to be immediately upstream or downstream (-1/+1) of a predicted *cis*-proline and hypothesised that proline isomerisation might influence the accessibility to these cryptic sites (**Supplementary Table 2**). The immunoglobulin (Ig)-like domains of human filamins are a particularly prevalent example, where 60 out of all 72 human filamin Ig-like domains contain a conserved, buried phosphoserine/*cis*-proline site at the C-terminus, and this *cis*-proline is completely structurally conserved in all known experimental structures of these domains^50^. We therefore studied two example domains, the 24^th^ repeat of human filamin C (FLNc24) and the 21^st^ repeat of filamin A (FLNa21). The results obtained for these two domains were essentially identical (**Fig. 4g-h**, **Extended Data Figs. 7-8**), in line with the high structural and functional conservation of this motif, and we thus describe FLNa21 as an example.

The buried serine residue (phosphoserine site) remained completely inaccessible to solvent in unbiased MD simulations (lasting 1 μs) starting from the native structure with the proline in *cis* (S2319 and P2320 for FLNa21, **Fig. 4g**, **Extended Data Figs. 8a-e**). We then used a metadynamics approach to accelerate isomerisation of the proline peptide bond (S2319-P2320, see Methods). In line with all available crystal structures and NMR data on a *D. discoideum* homologue, FLN5^50^ (26% and 23% sequence identity to FLNa21 and FLNc24, respectively), these simulations predicted the proline to be in *cis* 95–98% of the time for FLNa21 and FLNc24, respectively (**Extended Data Figs. 7f-h, 8f-h**). Thus, the *trans*-proline state constituted a minor (2–5%) conformational population at equilibrium, although it has been observed experimentally that this *trans*-proline state can accumulate as a kinetic folding intermediate during refolding of the FLN5 homologue^50^, where the serine-equivalent residue M741 is solvent-exposed in the folding intermediate but buried in the native state (**Extended Data Figs. 7o-p**). Moreover, during *de novo* folding on the ribosome (co-translational folding) we have recently shown two folding intermediates to be highly populated (>40%)^58,59^, and these are conserved across filamin domains with one intermediate being in the *trans*-proline conformation^7^. In this *trans* conformation, our metadynamics simulations revealed that S2319 (FLNa21) becomes fully exposed (**Fig. 4g**, **Extended Data Figs. 8h-i**). Simulations of the phosphorylated protein further showed that the protein becomes trapped with P2320 in *trans* and with pS2319 remaining solvent-exposed (**Supplementary Fig. 7**), which results in local structural perturbations and increased dynamics (**Extended Data Figs. 8j-n**). These destabilising effects agree with experimental studies of the *D. discoideum* homologue FLN5, which is destabilised by 4 kcal mol^-1^ when the equivalent proline P742 is mutated to alanine (P742A)^50^. Such destabilising effects may be relevant for filamin proteins due to their functional importance in actin crosslinking, signalling, and mitosis, which relies on mechanical stability of the Ig-like domains^60–63^. Overall, this suggests a mechanism where filamin Ig-like domains may become phosphorylated at this serine/proline motif to decrease mechanical stability, where a minor *trans*-proline population under native conditions and co-translational folding intermediates^50,58,59^ provide temporary access to these buried phosphoserine sites.

We also further investigated PARK7, which contains another site, Y67, that is buried under helix 2 in the native structure^53^ and immediately downstream of a *cis* proline, P66 (**Fig. 4f**). We hypothesised that the isomerisation state of P66 may influence the conformational preferences of Y67, which remained completely buried in unbiased simulations lasting 1 μs (**Extended Data Fig. 9a-c**). We accelerated sampling of the local conformational landscape using metadynamics (see Methods), which revealed that Y67 exposure is intricately linked to the isomerisation state of P66. In solution, our simulations suggest PARK7 to exist in two conformational states (each ∼50% in population) with P66 either in *cis* or *trans* (**Extended Data Fig. 9g-k**). When P66 is in *cis*, Y67 remains predominantly buried, while a *trans*-P66 conformation promotes exposure of Y67 due to a local conformational change (**Fig. 4f**, **Extended Data Fig. 9g-k**). When Y67 is phosphorylated, the P66 *cis* population decreases from 45 ± 1 % to 31 ± 3 % and pY67 remains solvent-exposed (**Extended Data Fig. 10h-k, Supplementary Fig. 7**). In the exposed pY67/*cis*-P66 state, the dimer is seen to remain stable in unbiased simulations up to 4 μs, apart from partial unfolding of helix 2 and increased dynamics in the vicinity of the phosphorylation site (**Extended Data Fig. 10l-p**) in part due to electrostatic contacts of the phosphate group with K4 and R98 (**Extended Data Fig. 10q-s**). We therefore conclude that similar to filamin domains, an adjacent *cis* proline plays an important role in regulating access to Y67, which upon phosphorylation induces a local conformational change and remains solvent-exposed.

### Buried phosphorylation sites and protein folding

We noticed that the filamin Ig-like domains FLNc24 and FLNa21 harboured additional phosphorylation sites with low solvent accessibility, varying from partially to completely buried (**Extended Data Figs. 7q-r & 8o-p**). Notably, this included multiple tyrosine residues which are observed to remain buried in the hydrophobic core in unbiased simulations initiated from the native structure (**Fig. 4i**). This suggests these sites may become phosphorylated during global conformational dynamics, such as protein folding or unfolding. Despite the low sequence identity of filamin Ig-like domains (20–40%), their structures are remarkably well-conserved^50^, including the C-terminal *cis*-Pro/Ser motif harbouring a buried phosphoserine site (**Supplementary Table 3**). Moreover, we recently found that these domains fold co-translationally via conserved folding pathways^7^, and structural features of folding intermediates may thus potentially serve as recognition sites for kinases. Consistent with the notion that co-translational modifications may occur more broadly, we found that buried modification sites were significantly shifted towards the N-terminus by an average of -7 amino acids or 0.8–0.9% of protein length, suggesting early modification during translation (**Supplementary Fig. 8a-d**). Most eukaryotic proteins are multi-domain structures^64^, which can fold domains sequentially and co-translationally, and thus this analysis supports the idea that some of these modifications may occur co-translationally on folding intermediates or unfolded nascent chains where vectorial emergence of the N-terminus provides a kinetic window for modification prior to burial upon folding. Two other examples that highlight this potential interplay between folding and modification are the human protein tyrosine kinase BTK and malignant T-cell-amplified sequence 1 (MCTS1), which contain buried sites covered by secondary structure elements and at domain interfaces, respectively (**Fig. 4j**). Co-translational folding via stable folding intermediates that form during biosynthesis is a widespread observation^59,65–67^, and we therefore suggest that in addition to post-translational unfolding, some natively buried residues may become modified during protein biosynthesis and co-translational folding.

## DISCUSSION

Structural models from AlphaFold allowed us to study the structural evolution of protein modifications across species in unprecedented detail. We applied these to the study of orthologous residues across 200 species, as well as to human variants seen in genetic disorders and cancer, and investigated 23 sites and 3 flanking prolines across 7 proteins in detail using molecular dynamics simulations. We found that a substantial fraction of protein modification sites (17.6%) were buried within protein structures in their unmodified state, challenging the common assumption that modification sites must be solvent-exposed in the native structure. The increased evolutionary conservation of many of these buried sites, coupled with their decreased variability in the human population and their enrichment in disease-associated variants and mutations, supports their functional importance, as opposed to these sites representing traces of misfolding or experimental error, e.g. from high-throughput mass spectrometry. Our results also reconcile well, however, with the view that some buried sites do represent markers of protein misfolding^22^. In this context, we note that buried ubiquitination sites display an appreciable number of very slowly-evolving sites (**Fig. 2b**). Such conserved sites in the interior of a protein may indeed act as sensors of unfolding, leading to polyubiquitination and subsequent degradation, which notably occurs for up to 15% of human nascent polypeptides during biosynthesis on the ribosome^68^, though not all misfolded conformations are likely to be recognised^23^.

Analysing the contacts and variants found at modification sites lends further evidence to their functionality. When analysing the contacts made by buried PTM sites in their unmodified state, we found a significant enrichment for phosphorylation sites to be in proximity to positively charged residues (**Fig. 3f**). As the distance maxima we used for contact determination were relatively stringent (**Supplementary Table 4**), it is likely that this remains an underestimate of such potential coordination or “shielding” of doubly negatively-charged phosphorylated residues. We also found a depletion in natural variants (**Fig. 3a-b**) and an increase in disease- and cancer-associated variability at modification sites (**Fig. 3c-e**). Analysing the types of variants observed further supported the functionality of buried modification sites (**Fig. 3g-h**). The effect of a single amino acid change on a protein can be profound, including fold transition^69^. It is thus interesting to note that normally dynamic and reversible modification sites (e.g. phosphotyrosine, Y-p) might become functionally fixed through mutations, either in a constitutively active state through a modified-like residue (e.g. negatively charged aspartate, D), or constitutively inactive through a residue resembling the unmodified state (e.g. phenylalanine, F)^70,71^. These mimics are clearly imperfect, especially sterically and given the absence of an aromatic ring in aspartate in addition to its single negative charge^71^. However, phosphomimetic and other mimetic mutations have been used with success in mutational studies^42,44^, including aspartic acid for tyrosine phosphorylation at two nearly-buried sites with RSA 26% in Raf and calmodulin^26,70,72^. When studying the types of variants and mutations present at modification sites in highly validated Cancer Gene Census genes from COSMIC (**Fig. 3h**), we did indeed observe a significant enrichment of somatic mutations to aspartic acid as well as to phenylalanine at buried tyrosine phosphorylation sites. We likewise saw an enrichment for ClinVar clinically significant variants of structured tyrosine phosphorylation sites (Y-p) to aspartic acid (D) when relaxing our structural confidence filter slightly (pLDDT ≥ 70 without a low-PAE contact requirement, **Supplementary Fig. 5e**), lending further support to this hypothesis of partial mimicry. Importantly, tyrosines are the only phosphorylatable residues for which mutation to a negatively charged amino acid (D) is available via a single base change within their codons, rendering these mutations uniquely accessible (as well as to unmodified-like phenylalanine (F), **Fig. 3h**).

Our all-atom MD simulations have uncovered a range of possible mechanisms rationalising the exposure and modification of buried protein modification sites. Additional simulations of the phosphorylated states induced conformational changes and altered protein dynamics (**Fig. 4, Extended Data Figs. 3-10**), suggesting these sites are structurally important. Sites near the protein surface or disordered loops were observed to dynamically fluctuate between exposed and buried conformations on picosecond to microsecond timescales (**Fig. 4a**). Backbone dynamics resulting in local unfolding were also observed to make buried sites accessible for modification (**Fig. 4c-e**). Compared to other, completely exposed phosphorylation sites, these partially buried systems may act to induce slow modification kinetics, and these proteins may therefore have evolved to tune their dynamics to achieve a particular rate of response for their regulation or signalling. Moreover, by modifying these proteins in their folded states, kinases may be able to exploit structural features of their target proteins to achieve specificity^73^. The modification of some residues can also result in sequential exposure and modification of other sites close in the native structure, as illustrated for p27 by experiments^48^ and MD simulations^15^.

Another interesting mechanism exploits proline isomerisation, which can induce local conformational changes resulting in solvent exposure of otherwise buried phosphorylation sites (**Fig. 4f-h**). In this way, prolines may also confer slower kinetics in the phosphorylation process because proline isomerisation is likely rate-limiting or regulated by prolyl isomerases^74^, a diverse family of proteins^75^. Prolyl isomerases have been observed to be regulated by phosphorylation of their target motifs^76,77^, and *vice versa*, phosphorylation can be regulated by the isomerisation state of prolines in a consensus motif^78^. Our simulations predicting kinetic control of solvent exposure for buried phosphorylation sites via proline isomerisation may be, to our knowledge, a new regulatory layer for protein modification sites in structured protein regions.

Phosphorylation sites found buried in the hydrophobic core of proteins (**Fig. 4i-j**) suggest that these sites may become modified during protein folding, perhaps acting to influence folding pathways^19^ as a form of signalling and exerting long-term effects. Most proteins begin to fold during biosynthesis on the ribosome (co-translational folding)^58,59,65–67,79,80^, and the ribosome itself acts as a platform for many other enzymes that modify nascent polypeptides^81^. It is therefore possible that many buried modification sites are introduced during translation, which has been suggested by biochemical experiments^20^. For example, Akt has been shown to be phosphorylated co-translationally by mTORC2^18^, and cAMP-dependent protein kinase A can auto-phosphorylate during biosynthesis^17^. We have recently shown that folding pathways for similar proteins are conserved on the ribosome^7^, which suggests that the structural features of co-translational folding intermediates may aid in recognition and specificity for interacting with kinases. For example, FLNa21 has been shown to populate a co-translational folding intermediate with a disordered C-terminus and with P2320 in *trans*^7^: taken together with our simulations presented in this work, which show S2319 to be accessible when P2320 is in the *trans* conformation as opposed to *cis*, this raises the possibility that FLNa21 may become modified either in the unfolded or intermediate state. Phosphorylation of the equivalent serine S1533 (upstream of *cis*-P1534) in the upstream domain FLNa15 is, in fact, a mechanism in mitosis^63^ and underscores the potential functional importance of these signalling systems in filamins.

Protein modifications are generally considered a rapid and reversible mechanism for signalling and other biological processes^5^. However, when investigating buried protein modification sites, we found a broad range of exposure mechanisms spanning orders of magnitude in timescales, ranging from picosecond fluctuations to seconds for proline-gated phosphorylation sites^50^, with other sites remaining buried persistently in the native state. These persistently buried sites may be accessible and become modified during folding or global conformational changes, potentially affecting folding pathways and essentially capturing the information state of the cell at synthesis. We suggest that buried modification sites may represent a previously unappreciated set of longer-term signalling mechanisms for biological information processing, in agreement with the observation that some phosphorylated residues also remain partially and transiently buried in the modified state (**Supplementary Fig. 7**). Given the evidence of evolutionary, structural, cancer and genetic disease relevance we found for natively buried modification sites, we consider the regulation of their accessibility and kinetics through protein dynamics and folding to be a highly promising mechanism and concept for further biomedical investigation.

## METHODS

### Overview

We investigated the evolutionary conservation and structural dynamics of buried protein modification sites using conservation scoring, natural and disease variants, cancer mutations, contact analyses, and all-atom molecular dynamics methods.

### Obtaining protein modification sites

We included protein modification types with at least 1,000 known human canonical-isoform sites across four data sources (UniProt version 2022_04 from October 2022^32^, PhosphoSitePlus version 2022-10-22^33^, dbSNP’s 2022 version obtained 2022-10-26^84^ and Ochoa *et al.*^24^), leading to 17 different modification types, ranked here by descending number of known sites and abbreviated as follows: serine phosphorylation (S-p), threonine phosphorylation (T-p), lysine ubiquitination (K-ub), tyrosine phosphorylation (Y-p), lysine acetylation (K-ac), lysine sumoylation (K-sum), arginine methylation (R-me), threonine glycosylation (T-gly), serine glycosylation (S-gly), lysine malonylation (K-mal), methionine sulfoxidation (M-ox), lysine methylation (K-me), asparagine N-linked glycosylation (N-gly), cysteine S-glutathionylation (C-glt), lysine succinylation (K-suc), cysteine S-palmitoylation (C-pal) and cysteine nitrosylation (C-nit). All of these are enzyme-catalysed and reversible^5,9,85–90^. We focussed exclusively on modification data for human proteins since other species had far fewer known sites.

### Proteins and residues included

We included all human reviewed canonical proteins from UniProt release 2022_04^32^. From these, we excluded small peptides below 16 aa as AlphaFold is unlikely to produce accurate structures for these^25^. We also excluded membrane residues as modification and control sites, i.e. residues in segments that are annotated as “intramembrane region” or “transmembrane region” in UniProt, since these would incorrectly be classified as solvent-exposed. We also generally excluded proteins that did not have modification sites or control residues for a given modification type, ensuring that proteins were modifiable and that controls were available.

### Structure models

We obtained structural models from the AlphaFold Protein Structure Database (AFDB)^25^, complemented by additional and updated structure predictions from AlphaSync^26^ for UniProt release 2022_04. This complementary set of structures enabled us to include nearly the entire proteome, including canonical proteins containing ambiguous FASTA^91^ characters such as B (aspartic acid or asparagine), Z (glutamic acid or glutamine), U (selenocysteine) or X. These very rare characters were replaced with reasonable substitutes unlikely to alter structure predictions^26^, enabling study of hundreds more proteins. We excluded three proteins for which structures were unavailable from AlphaSync 2025_01 (accessions P53794, Q99550 and Q6ZMR5), as their 2022_04 sequences had since become obsolete in UniProt. As mentioned, we also excluded small peptides below 16 amino acids.

### Structural contexts (buried, structured, disordered)

Residue-level absolute solvent-accessible surface area (SASA, Å^2^) and relative solvent-accessible surface area (RSA) were obtained from AlphaSync^26^, where they had been calculated as follows: DSSP version 4.2.2.1^92,93^ was used to calculate SASA. These absolute accessibility values were then converted into relative surface accessibility (RSA) values using reference values for amino acid surface areas within a polypeptide chain from Tien *et al.*^29^.

Any RSA values above 1, as might occur at the termini, were considered 100% on the AlphaSync website. Residues that were exposed ≤ 0.25 according to their relative solvent-accessible surface area (RSA) were considered *buried*, while residues with RSA > 0.25 were considered *surface* residues^30^. Residues with RSA_10_ < 0.55 were considered *structured*, while residues with an average RSA across a ±10 residue window (RSA_10_) of ≥ 0.55 were then considered to be part of an intrinsically *disordered* region^31^. We thus arrived at three mutually exclusive structural categories: *buried* residues, structured surface residues (referred to as *structured*) and intrinsically *disordered* surface residues.

### Structural confidence scores (pLDDT, PAE)

We obtained AlphaFold predicted Local Distance Difference Test (pLDDT) confidence scores at the residue level, and Predicted Aligned Error (PAE, Å) for intraprotein noncovalent contacts, from AlphaSync^26^. These residue-residue contacts had been determined in AlphaSync using a preview version of the Lahuta software (https://bisejdiu.github.io/lahuta/). Lahuta reports intra-protein atom-level contacts between residues based on atomic distances (**Supplementary Table 4**), including the type of contact (e.g. “IonicContacts”).

Both of these scores combined allowed us to select for buried residues whose AlphaFold structural model both displayed high local confidence (pLDDT e.g. ≥ 70) and at least one confident non-neighbour contact (PAE ≤ 2 Å), thus lending further confidence to the buried state. This use of PAE scores to determine contact confidence represents, to our knowledge, a significant novelty.

### Proline isomerisation

AlphaSync also provided omega dihedral angles (ω) calculated using the BioPython Bio.PDB package^94^. Proline isomerisation was defined as *trans* (common) or *cis* (rare) using ω, with absolute values ≤ 50° classed as *cis* and absolute values ≥ 130° classed as *trans*. Values in between were considered indeterminate.

### Homology clusters across species

For analysing modification site conservation across 199 representative species in addition to human, we used data from the Ensembl Comparative Genomics pipeline (Ensembl Compara release 108, October 2022)^95^. Note that Ensembl no longer provides homology clusters for all species it covers, but for this subset of representative species instead (https://www.ensembl.info/2021/01/20/important-changes-of-data-availability-in-ensembl-gene-trees-and-biomart/). The representative species are mostly vertebrates and of low-to-medium evolutionary distance to humans, including 23 other primates (**Supplementary Fig. 2e**), though outgroups including *S. cerevisiae* are included. We reasoned this dataset would optimally allow us to investigate conservation at shorter as well as longer evolutionary distances. We selected all human proteins whose UniProt 2022_04 sequence had an exact match in Ensembl release 108.

### Orthologue clusters from homology clusters

For each of these human proteins and each target species, we selected either its one-to-one orthologue or, in one-to-many orthology situations (https://useast.ensembl.org/info/genome/compara/homology_types.html), its best match.

These “best” one-to-many orthologues for each species were chosen based on whole-genome alignment (https://www.ensembl.org/info/genome/compara/Ortholog_qc_manual.html), synteny and sequence identity (https://www.ensembl.org/info/genome/compara/homology_method.html). The exact sorting criteria were, in descending order: 1. Ensembl Compara’s high-confidence orthology classification (yes/no), 2. the average of the orthologue’s gene order conservation and whole genome alignment scores, 3. its gene order score, 4. its whole genome alignment score, 5. percent sequence identity for the query sequence, and 6. percent sequence identity for the target sequence. Using these criteria, the highest-ranked orthologue was then selected as the single “best” (most likely to be functionally equivalent) one-to-many orthologue. This ensured that each of the 200 representative eukaryotic species included in the Ensembl Comparative Genomics pipeline is represented by at most a single protein in a given homology cluster. We alternatively tested discarding species with one-to-many orthology situations for a given human protein, but we observed that this led to a strong decrease of the number of species covered in each homology cluster (**Supplementary Fig. 2d**), as well as an overestimation of the number of human proteins with orthologues only among primates orthologues (**Supplementary Fig. 2d**).

### Building multiple sequence alignments and phylogenetic trees for orthologue clusters

The above step constituted filtering of Ensembl Compara’s homology clusters, which normally include paralogues. Our motivation was to avoid paralogues as these have been reported to diverge rapidly at post-translational modification sites following gene duplication^96^. After filtering, we needed to rebuild the multiple sequence alignment for each homology cluster for optimal alignment quality, as functionally divergent paralogues had been removed.

Modelling our approach on Ensembl Compara’s pipeline, we tested MAFFT version 7.505 (2022-04-10)^97^ in six different modes to generate multiple sequence alignments. MAFFT offers three main alignment modes: BLAST-like E-INS-i, global alignment-like G-INS-i and local alignment-like L-INS-i. Additionally, MAFFT can be provided with a user-defined phylogenetic guide tree (--treein) or generate its own from the sequences. We tested both options: either letting MAFFT generate its own starting tree (the default), or providing a version of the reference Ensembl Compara species tree (https://raw.githubusercontent.com/Ensembl/ensembl-compara/release/108/conf/vertebrates/species_tree.branch_len.nw) pruned for each homology cluster, removing species not present in a given orthologue cluster^98^. We subsequently benchmarked these six different modes using enzyme catalytic sites to select one primary method (described below).

We also backtranslated alignments to coding sequence (CDS) alignments^99^. We then reconstructed a phylogenetic tree for each of our orthologue-only clusters using TreeBeST^100,101^ for use with conservation scoring methods below, reconciling gene-specific trees with a predefined species guide tree as well as taking into account coding sequences in determining branch lengths.

### Conservation scores

We selected the three top-performing conservation scoring methods from an independent benchmark study^35^. Each of these used a different methodology: an information entropy-based method purely operating on alignment columns (the Jensen-Shannon Divergence^102^, which we inverted by subtracting it from 1 for simpler comparison with the other two methods), a Bayesian phylogenetics-based method based on an alignment as well as a phylogenetic tree and its branch lengths (Rate4Site evolutionary rates^103^), and a hybrid method that combines entropy and phylogenetic tree topology, but not branch lengths (the real-valued Evolutionary Trace score^104^).

When choosing the best conservation score and alignment method using M-CSA catalytic residues, we used all identical amino acids from the same set of proteins as controls. As M-CSA included disordered as well as membrane residues, we did not exclude these from the controls, though we verified that the results were equivalent when filtering out either or both types (data not shown).

Ver rarely, sequence-based mapping between UniProt and Ensembl produced multiple hits.

This occurred when multiple genes encode precisely the same protein sequence, e.g. for calmodulin. In Ensembl Compara, these different genes were then present in different homology clusters. In these cases, we used the minimum reported conservation score with the reasoning that additional copies of a gene can compensate for mutations, with the best-conserved copy being the most accurate indicator of evolutionary constraint.

### Benchmarking conservation scores and alignment methods to select a main approach

A systematic benchmarking of the three conservation scores mentioned above, as well as six different method variants for the orthologue cluster realignment for each, was implemented and performed using catalytic residues from the Catalytic Site Atlas (M-CSA)^36^. The expectation for these residues is for them to be highly conserved, as an enzyme would be non-functional without them, which provides clear and objective targets that a conservation score should be able to differentiate well from other residues.

We used all identical amino acids (potentially catalytic residues) from the same set of proteins (enzymes) as controls. As M-CSA included a small fraction of disordered as well as UniProt-annotated intramembrane residues, we did not explicitly exclude these from the controls, though we verified that the results were highly similar when filtering out either or both types (data not shown).

The results indicated that the best-performing approach was the Rate4Site evolutionary rate method, calculated using Ensembl species tree-guided phylogenetic tree reconstructions and BLAST-like E-INS-i alignments produced by MAFFT, as well as using the “single best paralog” method described above, rather than using one-to-one orthologs only. We thus selected this combination as our main approach (presented in e.g. **Fig. 2b**).

### Analysing conservation across species

To test whether evolutionary rates were significantly lower for modified compared to unmodified control residues for a given modification and structural category, indicating conservation (**Fig. 2b, Extended Data Fig. 2a-c, Supplementary Fig. 1a-e**), we used one-tailed asymptotic Wilcoxon-Mann-Whitney tests of sites stratified at the protein level with a minimum block size of 2 and FDR-corrected for multiple testing.

### Variants

For all variant analyses, we started from genomic coordinates and used the Ensembl Variant Effect Predictor (Ensembl version 108)^105^ to determine protein-level effects consistently. We also limited our analyses to single-nucleotide variants (SNVs) resulting in amino acid changes (missense), reasoning that compared to other variants, SNVs are most likely to emerge independently through mutation rather than being inherited, and that our phenotypic interest is at the protein level only. Additionally, modification sites are unique among signalling features in that they contain a single residue whose mutation results in abolishing the feature, rendering these highly amenable to study by SNVs. As we primarily analysed complete depletion in SNVs (the fraction of invariant residues), we reasoned that focussing on missense SNVs would most directly allow us to address evolutionary pressure on modification sites in the human population.

For consistency, we further limited our analyses to only included proteins that could be mapped by exact sequence match between UniProt 2022_04 and Ensembl 108. Out of 18,822 canonical UniProt proteins, 18,647 (99.1%) had at least one variant in gnomAD v4, indicating effectively complete coverage and ruling out artifactual observation of depletion.

We analysed four main datasets: gnomAD version 4^38^ (whole-genome and whole-exome sequencing, using “joint” allele counts and frequencies) for natural variants, ClinVar release 2023-07-22^41^ for variants associated with genetic disease, COSMIC version 99^39^ for somatic mutations in cancer and Genomenon MasterMind Cited Variants Reference (CVR-2) release 2024-01-03^106^ as by far the largest dataset, based on automated literature mining.

For residue enrichment analyses using Fisher’s Exact Test comparing occurrence of modification with occurrence of the variant, we also required at least 10 modified and 10 control residues per modification type and structural category (buried/structured/disordered). In **Fig. 3**, we increased the stringency to at least 40 modified and 40 control residues.

### Prioritising candidate buried sites for analysis

We used the results and data from the above analyses to guide our selection of candidates for molecular dynamics simulations (resulting in **Supplementary Table 2**). The criteria were, in detail: 1. The residue should be buried (RSA ≤ 0.25). It should be close to solvent-exposed residues (**Fig. 1e**), relatively inclusively defined as a distance to the nearest surface residue (RSA > 0.25) of up to 5 residues along the primary sequence. 3. The residue should have a confident AlphaFold structure prediction for its other characteristics (such as solvent accessibility) to be reliable, defined as an AlphaFold pLDDT score of ≥ 70 (“confident”) for the residue itself as well as on average within a ± 10 amino acid region around it. 4. At least one experimental structure should be available in the PDB^107^ to help guide and parametrise molecular dynamics simulations. When manually selecting our candidate sites for simulation (**Fig. 4a-i**), we specifically looked for NMR structures. 5. The residues should be in a structured domain of reasonable size for efficient simulation, both according to AlphaFold-based prediction of a structured domain (average RSA < 0.55 within a ± 10 amino acid region) and according to a PDB structure. 6. The residue should only have rare genetic variants (1 in 10,000 individuals or fewer) to help ensure functional importance. 7. The residue should display reasonable conservation across species (evolutionary rate ≤ 0.3). For completeness, we also manually added all modification sites for filamins FLNa-c due to their prominence in our study (**Fig. 4g-i**). In combination, these criteria resulted in the identification of 4,507 promising buried sites across 1,346 proteins.

### Unbiased molecular dynamics simulations

Molecular dynamics (MD) simulations were performed with the GROMACS version 2023^108^. For each unmodified protein listed in **Supplementary Table 5**, three independent simulations were run starting from the experimental structure with different initial velocities. The simulations were parameterised with the CHARMM36m (C36m) protein parameters and CHARMM-modified TIP3P water model^109^. All systems were simulated with default protonation states at pH 7.4, except for PARK7, where it has been shown that Glu18 and Cys106 are protonated and deprotonated, respectively^53^. All proteins were placed in the centre of a box (dodecahedron) using gmx editconf -d 1.2 nm, solvated, neutralised, and ions were added according to the conditions in **Supplementary Table 5**. The simulation conditions were chosen to mimic the conditions under which the experimental structures were determined, or under which the proteins have been studied previously. Simulation systems were then energy minimised using the steepest descent algorithm. All following dynamics simulations used the leap-frog integrator, 2 fs timestep and LINCS constraints^110^ all hydrogen bonds. A cut-off distance of 1.2 nm was used for nonbonded interactions with a function between 1.0 and 1.2 nm for Lennard-Jones interactions. Long-range electrostatics were calculated using the Particle Mesh Ewald (PME)^111^ method with cubic interpolation and a fourierspacing of 0.16 nm. Temperature coupling was achieved with the velocity rescaling algorithm^112^ (coupling constant = 0.1 ps). Equilibration was performed for 500 ps in the presence of position restraints on all heavy atoms (force constant = 1000 kJ mol^-1^ nm^-2^) in the NVT ensemble. Simulations with positions restraints were then further equilibrated in the NPT ensemble at 1 bar for 500 ps, using the Berendsen barostat^113^ (compressibility = 4.5x10^-5^ bar^-1^, coupling constant = 2 ps). For production simulations, the position restraints were removed and pressure coupling switched to the Parrinello-Rahman algorithm^114^. Simulations were run for 1 μs, saving coordinates for analysis every 20 ps (50,000 snapshots for each trajectory).

Starting structures for simulations of phosphorylated FUS, SRSF2, ACYP1, GGCT, PARK7 (pS47 and pS57) were taken from the unbiased MD trajectories of the wild-type proteins, using the snapshots that exhibited the maximum sidechain SASA of the unmodified PTM sites. For FLNc24, FLNa21, and PARK7 pY67, where the PTM sites were completely buried throughout all unbiased simulations, we used structures from the metadynamics simulations described below, which enhanced the sampling of conformational space and allowed sampling of conformations where their PTM sites are accessible. Structures were taken from the local free energy minimum corresponding to the accessible state. The phosphate groups were then added using PyMol (version 2.3) and modelled in the deprotonated state (charge -2 *e*). Triplicate simulations were then performed with different initial velocities in each case. Box sizes were increased (gmx editconf -d 1.5-2 nm) to accommodate potential conformational changes and local unfolding. Simulations were then prepared and equilibrated with an equivalent protocol as described above.

RMSD, RMSF and SASA analyses were performed with their respective GROMACS gmx tools rms, rmsf and sasa, respectively^108^. All-atom dimer interface RMSD values (for GGCT and PARK7) were calculated using heavy atoms within 6.0 Å of any other atom of the opposite chain. Distances were calculated using the Python package MDAnalysis^115^. For all simulation results, we present ensemble-averaged properties as the mean with the standard error of the mean (SEM) obtained from three independent replicates.

### Proline free energy calculations (metadynamics): FLNa21 and FLNc24

Well-tempered metadynamics^116^ (WT-METAD) simulations were performed with GROMACS (version 2020)^108^ and PLUMED (version 2.6)^117,118^ to efficiently sample proline isomerisation of FLNc24 Pro2719 and FLNa21 Pro2320.. Triplicate metadynamics simulations were initiated from different coordinates, i.e., fully equilibrated simulation boxes after 1 μs of unbiased MD. The mean and standard error of the mean were then calculated from the triplicate simulations for all quantitative analyses. WT-METAD simulations were run for 1 μs each with equivalent simulation parameters as described above. Two dihedral angles, ζ (Cα_i-1_ – O_i-1_ – Cδ_I_ – Cα_i_) and ψ (N_i_ – Cα_i_ – C_i_ – N_i+1_), where *i* corresponds to the proline residue of interest, were used as collective variables, capture the isomerisation state (∼ 0 for *cis* and ∼± π for *trans*) and C-terminal amide orientation, respectively^119^. The Gaussian deposition rate was set to 1 ps^-1^, and a Gaussian width of 0.07 rad, height of 0.8 kJ mol^-1^, and bias factor of 10, were used. Simulation frames were saved every 5 ps and reweighted as previously described by Tiwary *et al.*^120^ after removing the first 20% (200 ns) of each trajectory due to equilibration of the bias potential.

### Metadynamics simulations of PARK7

We also studied the isomerisation of PARK7 Pro66 and its effect on the accessibility of Tyr67. WT-METAD simulations with GROMACS (version 2020)^108^ and PLUMED (version 2.6)^117,118^ were used to simultaneously bias the proline isomerisation state of Pro66 and protein-sidechain contacts with Tyr67. For the former, we used the improper dihedral angle ζ as described above. Protein-sidechain contacts with Tyr67 were biased to encourage the sampling of conformational states with Tyr67 more solvent-exposed. In the simulations, this was described with the coordination collective variable, *C*, which effectively quantifies the number of contacts between the Tyr sidechain and the protein.

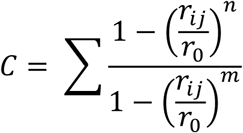

COORD is summed over all atom pairs where one atom belongs to the tyrosine sidechain (excluding the Cβ atom) and another atom is part of chain A of the dimer. Only heavy atoms were included in the calculation of *C* and *r_0_, n,* and *m* were set to 0.6 nm, 6, and 12, respectively. *C* and ζ were then biased with a Gaussian deposition rate of 1 ps^-1^, Gaussian height of 1.8 kJ mol^-1^, and bias factor of 40. The Gaussian widths were set to 2.0 and 0.07 rad, respectively. All metadynamics simulations were initiated from different coordinates, i.e., fully equilibrated simulation boxes after 1 μs of unbiased MD. The mean and standard error of the mean were then calculated from the triplicate simulations for all quantitative analyses. Simulation frames were saved every 5 ps and reweighted with the Tiwary method^120^ after removing the first 20% (200 ns) of each trajectory due to equilibration of the bias potential.

## METHODS REFERENCES

(To be divided out into a separate reference section)

## DATA AVAILABILITY

The AlphaSync data used in this study are available for download and via REST API at alphasync.stjude.org. The MD trajectories will be made available on Zenodo upon publication.

## CODE AVAILABILITY

The source code for the analyses in this study is available at github.com/langbnj/ptms.

AlphaSync’s source code is available at github.com/langbnj/alphasync.

## ACKNOWLEDGEMENTS

We thank Duccio Malinverni and other members of the Babu group for helpful input and discussions and acknowledge ALSAC (https://www.alsac.org) for financial support. This work was also supported by a Wellcome Trust Investigator Award (to J.C., 206409/Z/17/Z). We acknowledge the use of computational resources provided by the Baskerville Tier 2 HPC service (https://www.baskerville.ac.uk/). Baskerville was funded by the EPSRC and UKRI through the World Class Labs scheme (EP/T022221/1) and the Digital Research Infrastructure programme (EP/W032244/1) and is operated by Advanced Research Computing at the University of Birmingham. This project also made use of time on HPC resources on Archer2 (ARCHER2 UK National Supercomputing service, https://www.archer2.ac.uk) granted via the UK High-End Computing Consortium for Biomolecular Simulation, HECBioSim (http://hecbiosim.ac.uk), supported by EPSRC (grant no. EP/R029407/1 and EP/X035603/1). We are also grateful to the UK Materials and Molecular Modelling Hub for computational resources, which is partially funded by the EPSRC (EP/T022213/1, EP/W032260/1 and EP/P020194/1).

## AUTHOR CONTRIBUTIONS

B.L. led the project and M.M.B set the overall direction of research. B.L. and J.O.S. wrote the manuscript. J.O.S. led the molecular dynamics simulations and their interpretation guided by J.C., while R.K. provided the original inspiration and additional insight. All authors read and edited the manuscript.

## ADDITIONAL INFORMATION

Correspondence and requests for materials should be addressed to B.L., J.O.S., J.C. and M.M.B. Supplementary Information is available for this paper.

## EXTENDED DATA FIGURE LEGENDS

**Extended Data Figure 1.**
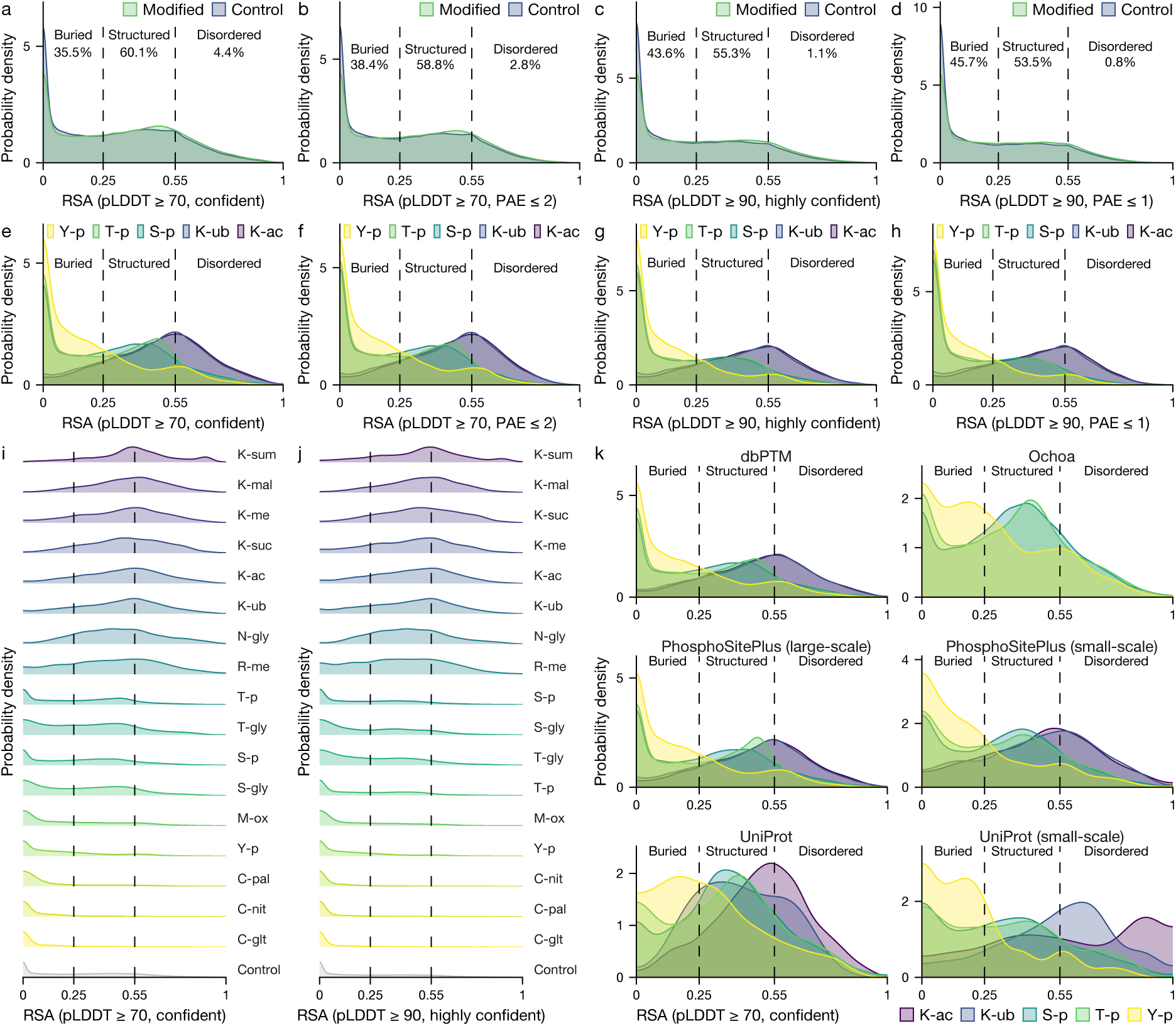
Many human protein modification sites are buried within proteins, across modification types, data sources, and source subsets, even when using stringent structural model confidence filtering. Please refer to **Fig. 1** for comparison. **a-e**, A substantial fraction of protein modification sites remains classified as buried when applying various stringent AlphaFold pLDDT score thresholds, as well as when requiring at least one contact with PAE ≤ 2 or ≤ 1 Å. When applying these thresholds, the proportion of residues classified as disordered using the average RSA across a ± 10 residue window strongly decreased. **e-h**, All major modification types continued to show a significant fraction of buried sites when applying structural confidence filtering. **i-j**, Structural preferences continued to differ between modification types, with some displaying a notable fraction of essentially completely buried sites (SASA near 0 Å^2^). **k**, Different modification data sources and subsets continued to show similar and noteworthy fractions of buried modification sites for major modification types.

**Extended Data Figure 2.**
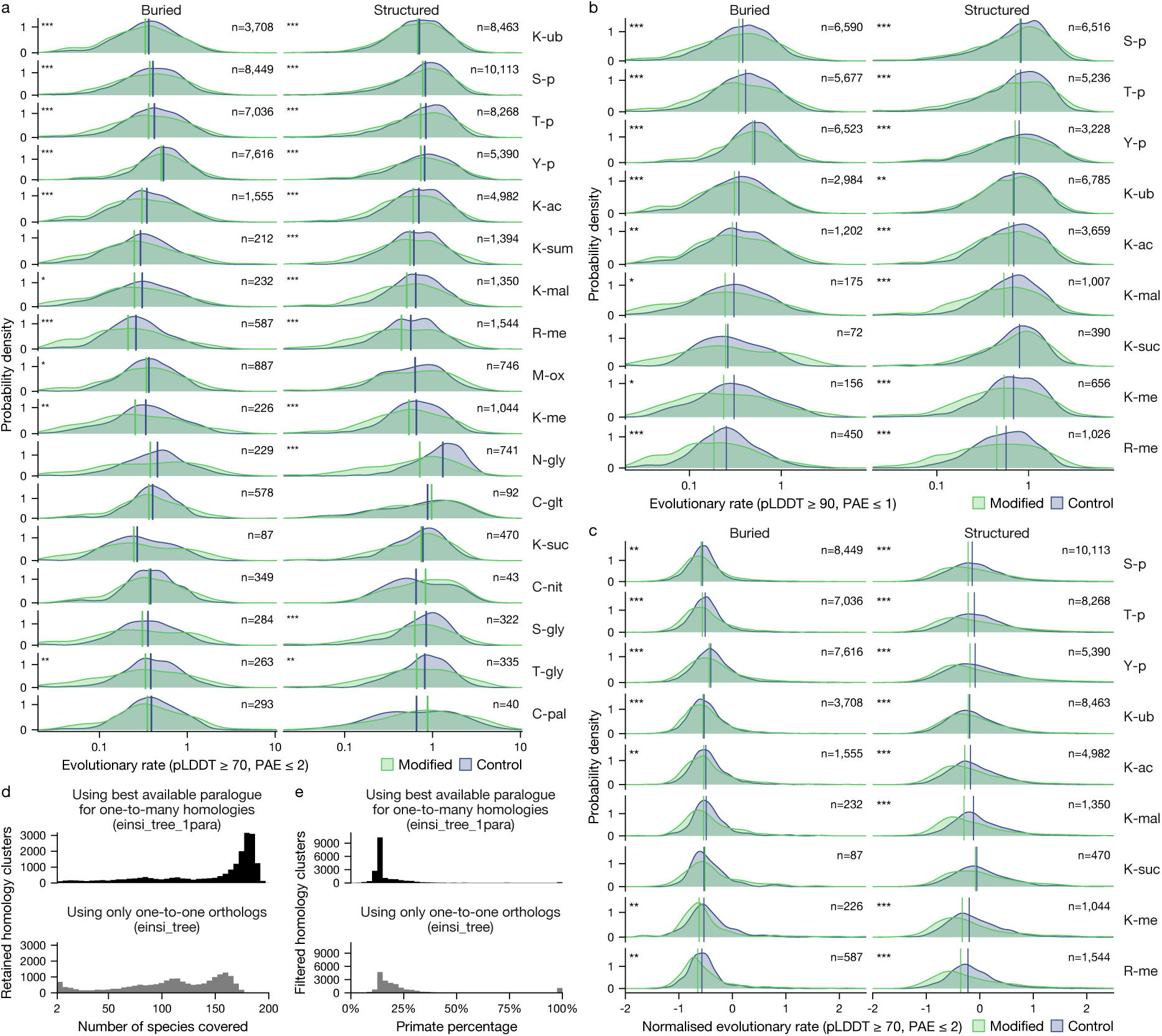
Buried residues remain significantly conserved with stringent structural model confidence filtering and protein-level normalisation. Please refer to **Fig. 2b** for comparison. **a**, Extended version of **Fig. 2b** showing all 17 modification types with over 1,000 known human sites in our dataset, showing significant conservation of buried sites for most types. **b**, More stringent structural confidence filtering continues to show highly significant conservation for buried sites. **c**, Evolutionary rate normalisation at the protein level continues to show highly significant conservation for buried sites. **d**, Using the best-matching homologue in one-to-many orthology situations (black), rather than discarding these (grey) (see Methods), led to a much higher number of species with orthologues for a given human protein, generally close to the maximum of 199 Ensembl Compara species. **e**, Likewise, including best-matching one-to-many orthologues (black) decreased the number of orthology clusters that exclusively contained primates, which were presumably artifactual.

**Extended Data Figure 3.**
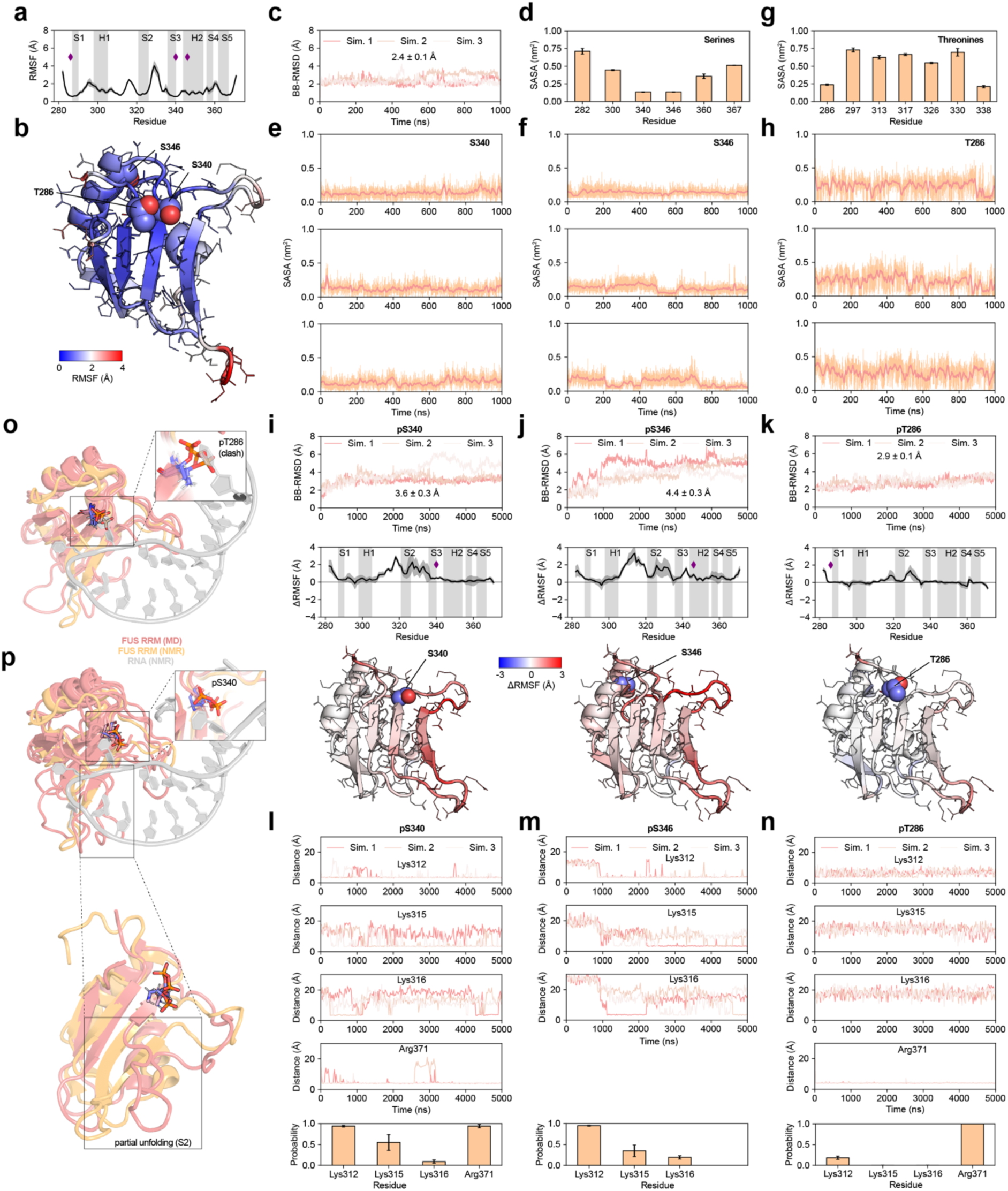
FUS RRM dynamics characterised by molecular dynamics simulations. **a,** Backbone dynamics quantified by the RMSF of the Cα atoms along the sequence of FUS. **b,** Residue-specific RMSF mapped on the NMR structure of FUS RRM (PDB 2LCW). **c,** Backbone RMSD of FUS RRM simulations calculated with respect to the experimental structure (PDB 2LCW). **d,** Average sidechain solvent-accessible surface area (SASA) of all serine residues. **e-f,** Sidechain SASA of buried PTM sites S340 and S346 as a function of simulation time with a moving average over 0.2 ns. **g,** Average sidechain SASA of all threonine residues. **h,** Sidechain solvent-accessible surface area (SASA) of buried PTM site T286 as a function of simulation time with a moving average over 0.2 ns. **i-k,** Backbone RMSD of phosphorylated FUS variants pS340, pS346, and pT286 with respect to their input structures (top). Change in RMSF with respect to wild-type FUS RRM across the protein sequence (middle) and mapped on the experimental structure (bottom). The diamonds indicate the position of the PTM sites. **l-n,** Contact distances between the phosphorylated residues (pS340, pS346, and pT286, phosphorus atom) and positively charged residues (NZ atom for lysine, CZ atom for arginine). The bottom barplot shows the average contact probability (5 Å contact cut-off). **o,** Superimposed structures obtained after 5,000 ns of MD (pT286 variant, three replicates) and the NMR structure of FUS RRM bound to RNA (PDB 6GBM^54^), highlighting steric clashes between pT286 and the RNA. **p,** Superimposed structures obtained after 5,000 ns of MD (pS340 variant, three replicates) and the NMR structure of FUS RRM bound to RNA (PDB 6GBM), highlighting steric clashes between pS340 and the RNA (top) and partial unfolding of the FUS RRM β-sheet (bottom). All averaged quantities represent the mean ± SEM from three independent replicates.

**Extended Data Figure 4.**
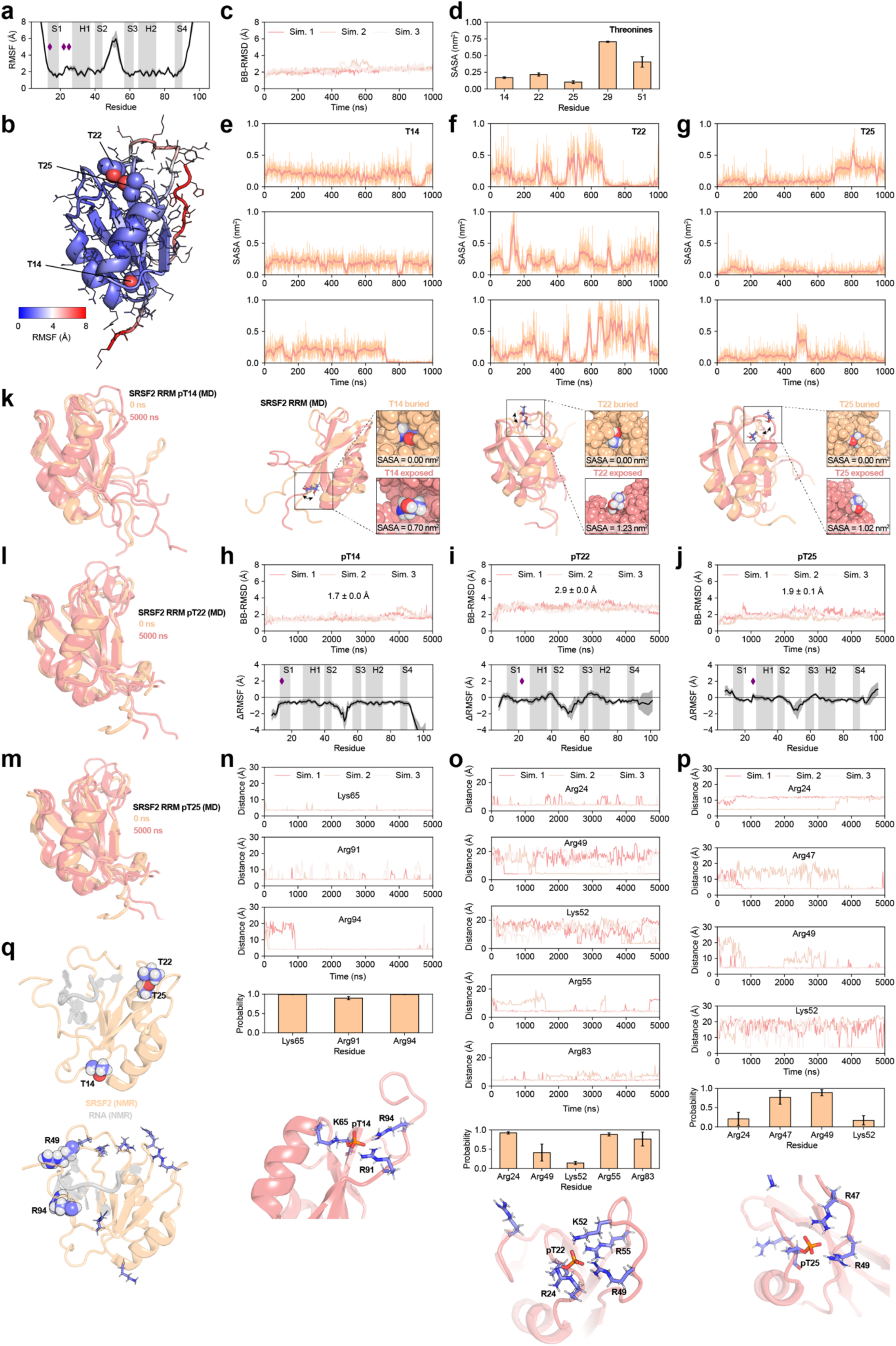
SRSF2 RRM dynamics characterised by molecular dynamics simulations. **a,** Backbone dynamics quantified by the RMSF of the Cα atoms along the sequence of the SRSF2 RRM. **b,** Residue-specific RMSF mapped on the NMR structure of SRSF2 (PDB 2KN4). **c,** Backbone RMSD of FUS RRM simulations calculated with respect to the experimental structure (PDB 2KN4, residues 13-90). **d,** Average sidechain solvent-accessible surface area (SASA) of all threonine residues. **e-g,** (top) Sidechain SASA of buried PTM sites T14, T22, and T25 as a function of simulation time with a moving average over 0.2 ns. (bottom) Structures of buried and exposed states sampled in the simulations. **h-j,** Backbone RMSD of phosphorylated SRSF2 variants pT14, pT22, and pT25 with respect to their input structures (top). Change in RMSF with respect to wild-type SRSF2 across the protein sequence (bottom). The diamonds indicate the position of the PTM sites. **k-m,** Superimposed MD structures at the beginning and end of unbiased all-atom simulations of phosphorylated SRSF2 variants. **n-p,** Contact distances between the phosphorylated residues (pT14, pT22, and pT25, phosphorus atom) and positively charged residues (NZ atom for lysine, CZ atom for arginine). The barplot shows the average contact probability (5 Å contact cut-off). Representative structures showing interactions between phosphorylated and positively charged residues (bottom). **q,** NMR structure of SRSF2 bound to RNA (PDB 2LEB^56^) highlighting the locations of the PTM sites (T14, T22, and T25) and positively charged residues interacting with phosphorylated residues (see panels n-p). R49 and R94 make direct contact with the RNA backbone. All averaged quantities represent the mean ± SEM from three independent replicates.

**Extended Data Figure 5.**
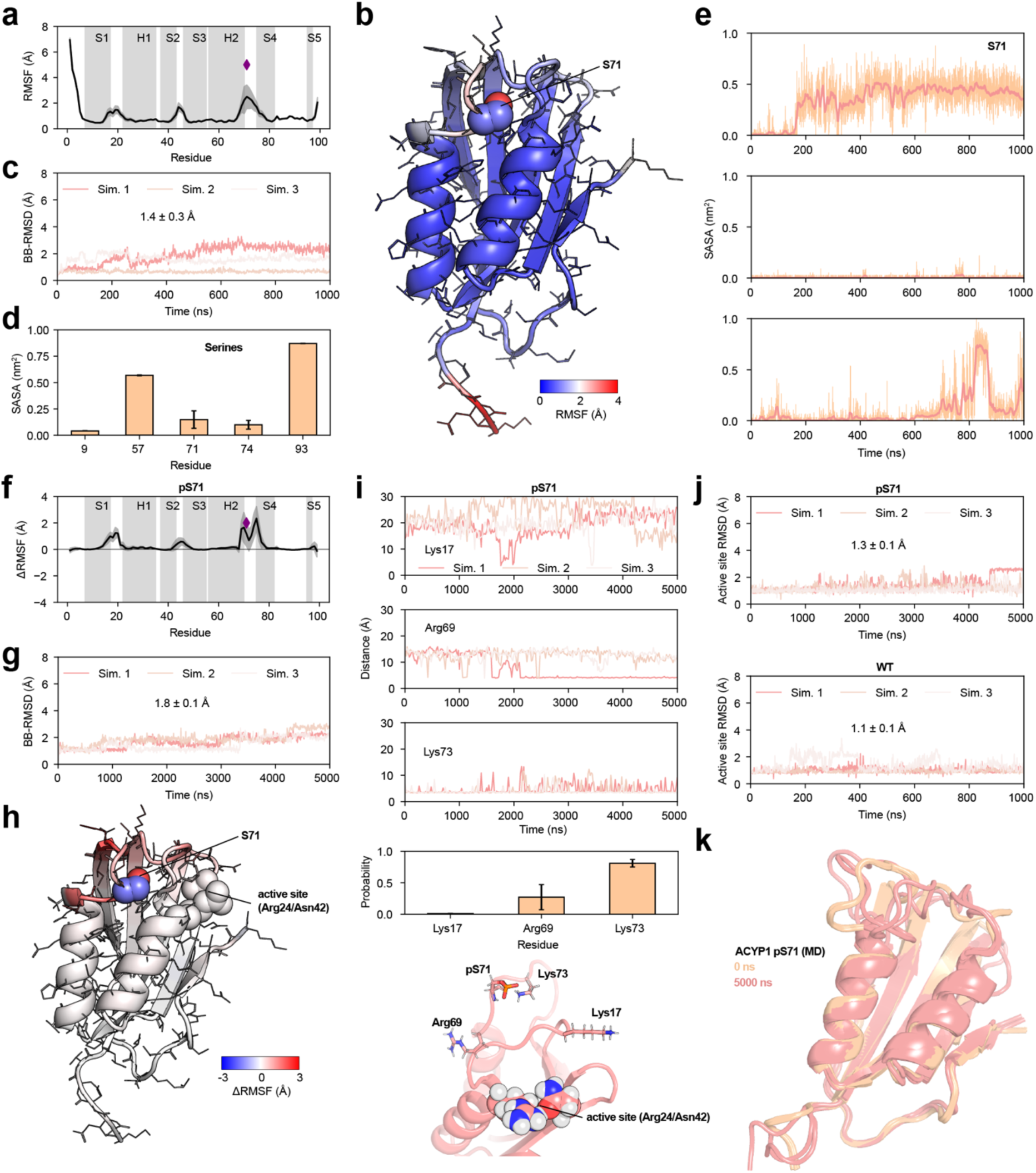
ACYP1 dynamics characterised by molecular dynamics simulations. **a,** Backbone dynamics quantified by the RMSF of the Cα atoms along the sequence of ACYP1. **b,** Residue-specific RMSF mapped on the structure of ACYP1 (PDB 2VH7). **c,** Backbone RMSD of ACYP1 simulations calculated with respect to the experimental structure (PDB 2VH7, residues 6-99). **d,** Average sidechain solvent-accessible surface area (SASA) of all serine residues. **e,** Sidechain SASA of the buried phosphoserine site S71 as a function of simulation time with a moving average over 0.2 ns. **f,** Change in RMSF with respect to wild-type ACYP1 across the protein sequence. The diamond indicates the position of the PTM site. **g,** Backbone RMSD of phosphorylated ACYP1 (pS71) with respect to the input structures (top). **g,** Change in RMSF (pS71-WT) mapped on the structure of ACYP1 (PDB 2VH7). **i,** Contact distances between the phosphorylated pS71 and positively charged residues (NZ atom for lysine, CZ atom for arginine). The barplot shows the average contact probability (5 Å contact cut-off). A representative structure highlights the most probable interactions between pS71 and Lys73 (bottom). **j,** All-atom RMSD of the ACYP1 active site (Arg24 and Asn42) calculated with respect to the experimental structure for WT and pS71 ACYP1. **k,** Superimposed MD structures at the beginning and end of unbiased all-atom simulations of pS71 ACYP1. All averaged quantities represent the mean ± SEM from three independent replicates.

**Extended Data Figure 6.**
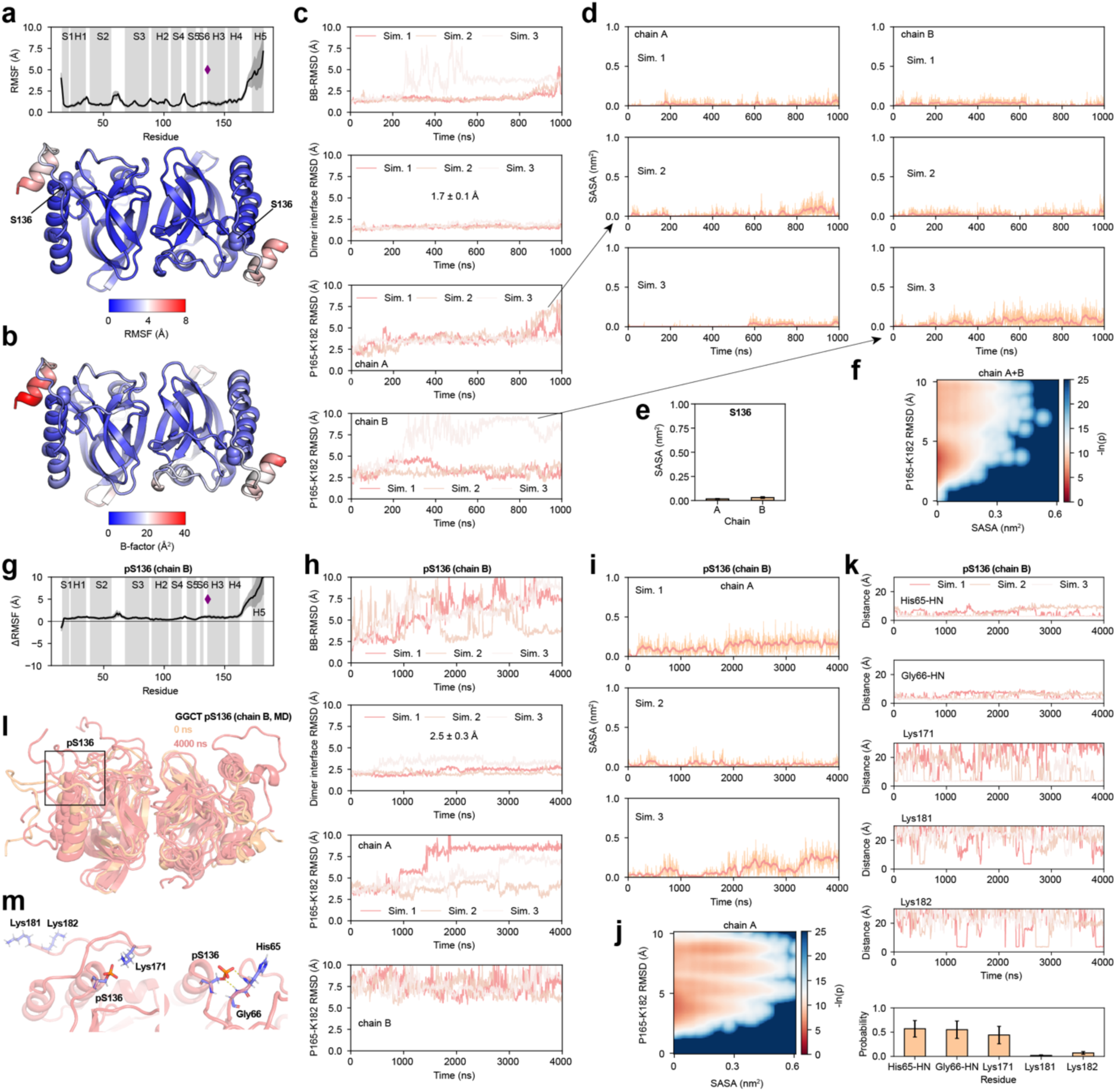
GGCT dynamics characterised by molecular dynamics simulations. **a,** Backbone dynamics quantified by the RMSF of the Cα atoms along the sequence of GGCT (average of both chains) and mapped on the structure of the GGCT dimer (bottom, PDB 3CRY). **b,** Temperature factors of the X-ray crystallography structure (PDB 3CRY). **c,** Backbone, all-atom dimer interface, and all-atom C-terminus (residues 165-182) RMSD of GGCT dimer simulations calculated with respect to the experimental structure (PDB 3CRY). **d,** Sidechain solvent-accessible surface area (SASA) of the buried phosphoserine site S136 in both chains as a function of simulation time with a moving average over 0.2 ns. **e,** Average sidechain SASA of S136. **f,** Free energy landscape as a function of S136 SASA and the all-atom RMSD of the C-terminus averaged over both chains. **g,** Change in RMSF of pS136 (chain B only) GGCT with respect to wild-type GGCT across the protein sequence. The diamond indicates the position of the PTM site. **h,** Backbone, all-atom dimer interface, and all-atom C-terminus (residues 165-182) RMSD of GGCT dimer simulations (with chain B pS136) calculated with respect to the input structure. **i,** Sidechain solvent-accessible surface area (SASA) of S136 in chain A (with chain B S136 phosphorylated) as a function of simulation time with a moving average over 0.2 ns. **j,** Free energy landscape as a function of S136 SASA and the all-atom RMSD of the C-terminus averaged (chain A only). **k,** Contact distances between the phosphorylated pS136 (chain B) and positively charged residues (NZ atom for lysine, CZ atom for arginine, chain B). The barplot shows the average contact probability (5 Å contact cut-off). **l,** Superimposed MD structures at the beginning and end of unbiased all-atom simulations of the pS136 (chain B) GGCT dimer. **m,** A representative structure highlights the most probable interactions between pS136 with Lys171, and hydrogen bonds with amide groups of His65 and Gly66. All averaged quantities represent the mean ± SEM from three independent replicates.

**Extended Data Figure 7.**
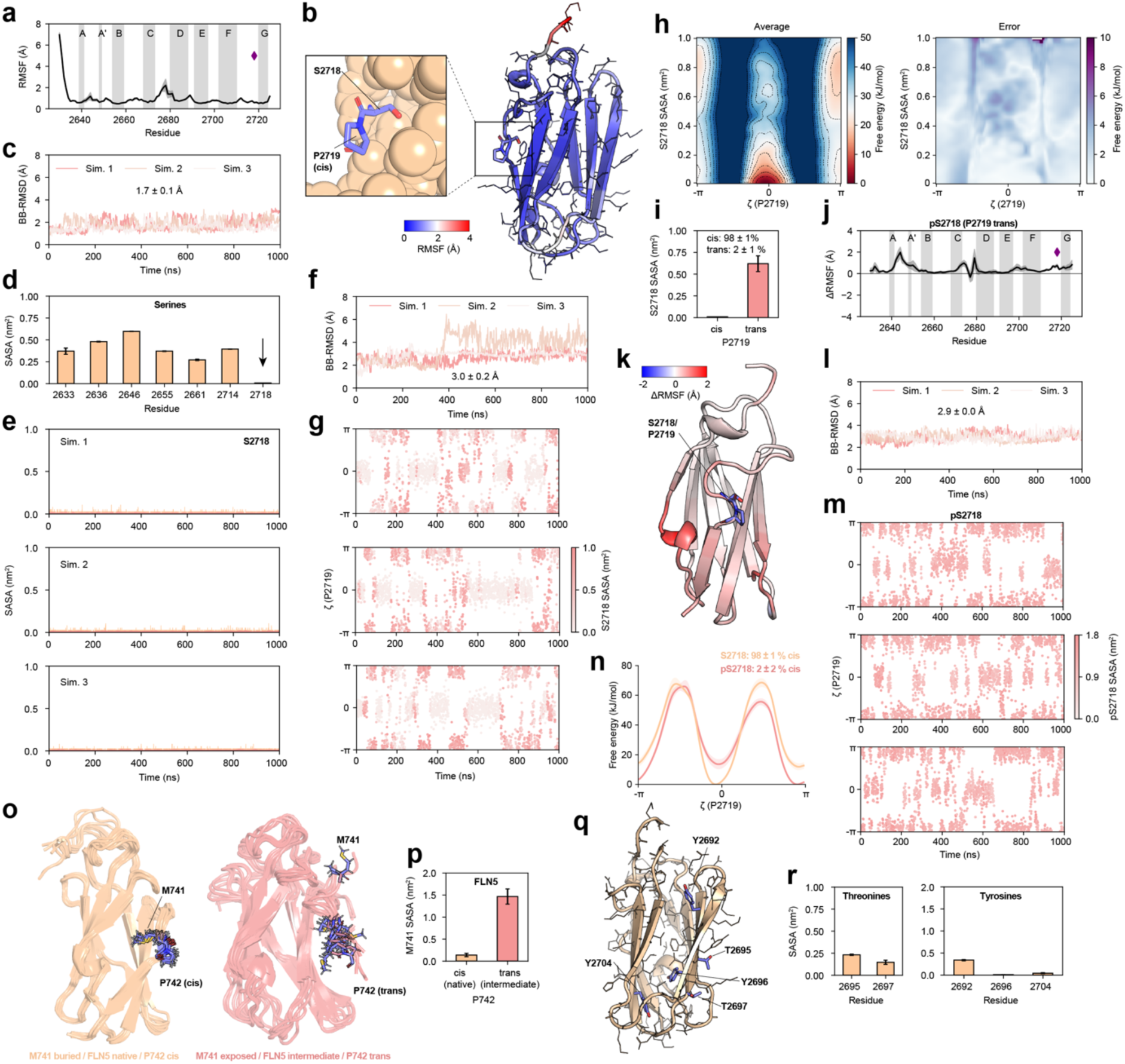
FLNc24 dynamics and proline isomerisation characterised by molecular dynamics simulations. **a,** Backbone dynamics quantified by unbiased MD simulations and the RMSF of the Cα atoms along the sequence of FLNc24 and **b,** mapped on the structure of FLNc24 (PDB 1V05), highlighting the location of the conserved *cis*-proline and upstream serine (residues 2718-2719). **c,** Backbone RMSD of unbiased FLNc24 simulations calculated with respect to the experimental structure (PDB 1V05). **d,** Average sidechain solvent-accessible surface area (SASA) of all serine residues. **e,** Sidechain SASA of the buried phosphoserine site as a function of simulation time with a moving average over 0.2 ns. **f,** Backbone RMSD of FLNc24 during proline (P2719) metadynamics simulations calculated with respect to the experimental structure (PDB 1V05). **g,** Proline metadynamics (P2719) simulations of FLNc24 showing the improper dihedral angle, ζ, where values near 0 and ± π correspond to the *cis* and *trans* conformations, respectively. The scatter points are coloured according to the sidechain SASA of S2718. **h,** Free energy landscape of FLNc24 as a function of P2719 isomerisation (ζ dihedral) and S2718 sidechain SASA calculated from triplicate metadynamics simulations. **i,** Average sidechain SASA of S2718 for P2719 in *cis* and *trans*, including the equilibrium populations of *cis* and *trans* calculated from triplicate metadynamics simulations. **j,** Change in backbone dynamics quantified by the RMSF of the Cα atoms along the sequence of FLNc24 from unbiased MD simulations of FLNc24 pS2718 with P2719 in *trans* (relative to wild-type) and **k,** mapped on the structure of FLNc24 (PDB 1V05) as the difference relative to wild-type (pS2719-WT). **l,** Backbone RMSD of unbiased FLNc24 pS2718 (P2719 *trans*) simulations calculated with respect to the experimental structure (PDB 1V05). **m,** Proline metadynamics (P2719) simulations of FLNc24 pS2718 showing ζ, coloured according to the sidechain SASA of pS2718. **n,** Free energy landscape of FLNc24 WT (S2718) and pS2718 as a function of P2719 isomerisation (ζ) and their equilibrium populations of *cis* P2719. **o,** NMR-restrained ensembles of FLNc24-homologue FLN5 (*D. discoideum*) in its native conformation and a folding intermediate where the P2719-equivalent proline P742 populates the *trans* conformation (ref.^50^). The sidechains of the S2718-equivalent methionine M741 is also shown in the stick conformation. **p,** Sidechain SASA of M741 in the native and intermediate state of FLN5 (errors denote the standard deviation of the ensembles). **q-r,** Additional partially and completely buried PTM sites in FLNc24 mapped on the experimental structure (panel q) and their average sidechain SASA calculated from triplicated unbiased MD simulations (panel r). All averaged quantities represent the mean ± SEM from three independent replicates unless stated otherwise.

**Extended Data Figure 8.**
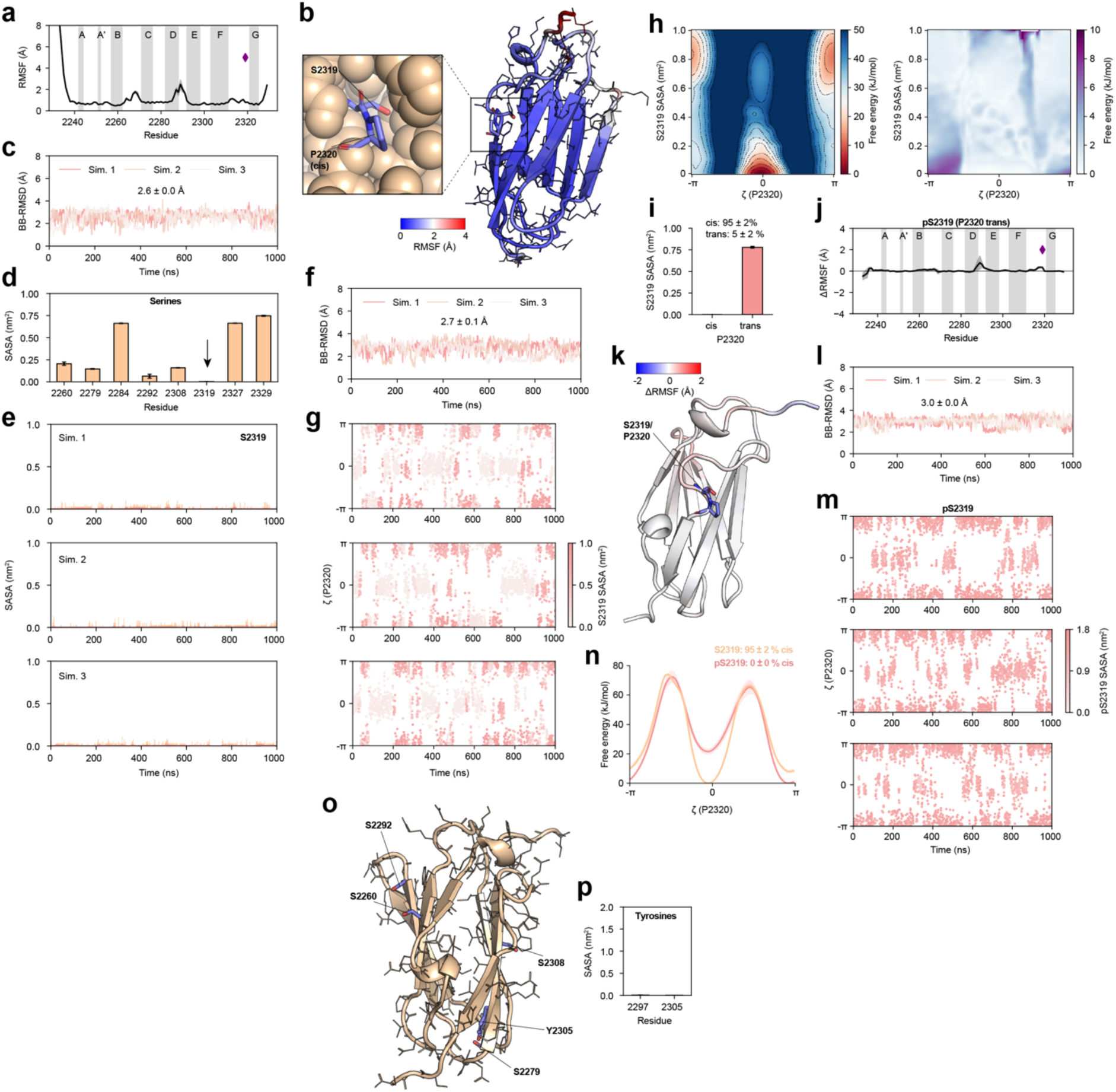
FLNa21 dynamics and proline isomerisation characterised by molecular dynamics simulations. **a,** Backbone dynamics quantified by unbiased MD simulations and the RMSF of the Cα atoms along the sequence of FLNa21 and **b,** mapped on the structure of FLNa21 (PDB 2W0P), highlighting the location of the conserved *cis*-proline and upstream serine (residues 2319-2320). **c,** Backbone RMSD of unbiased FLNa21 simulations calculated with respect to the experimental structure (PDB 2W0P). **d,** Average sidechain solvent-accessible surface area (SASA) of all serine residues. **e,** Sidechain SASA of the buried phosphoserine site as a function of simulation time with a moving average over 0.2 ns. **f,** Backbone RMSD of FLNa21 during proline (P2320) metadynamics simulations calculated with respect to the experimental structure (PDB 2W0P). **g,** Proline metadynamics (P2320) simulations of FLNa21 showing the improper dihedral angle, ζ, where values near 0 and ± π correspond to the *cis* and *trans* conformations, respectively. The scatter points are coloured according to the sidechain SASA of S2319. **h,** Free energy landscape of FLNa21 as a function of P2320 isomerisation (ζ dihedral) and S2319 sidechain SASA calculated from triplicate metadynamics simulations. **i,** Average sidechain SASA of S2319 for P2320 in *cis* and *trans*, including the equilibrium populations of *cis* and *trans* calculated from triplicate metadynamics simulations. **j,** Change in backbone dynamics quantified by the RMSF of the Cα atoms along the sequence of FLNa21 from unbiased MD simulations of FLNa21 pS2319 with P2320 in *trans* (relative to wild-type) and **k,** mapped on the structure of FLNa21 (PDB 2W0P) as the difference relative to wild-type (pS2319-WT). **l,** Backbone RMSD of unbiased FLNa21 pS2319 (P2320 *trans*) simulations calculated with respect to the experimental structure (PDB 2W0P). **m,** Proline metadynamics (P2320) simulations of FLNc24 pS2319 showing ζ, coloured according to the sidechain SASA of pS2319. **n,** Free energy landscape of FLNa21 WT (S2319) and pS2319 as a function of P2320 isomerisation (ζ) and their equilibrium populations of *cis* P2320. **o-p,** Additional partially and completely buried PTM sites in FLNa21 mapped on the experimental structure (panel o) and their average sidechain SASA calculated from triplicated unbiased MD simulations (panels d and p). All averaged quantities represent the mean ± SEM from three independent replicates unless stated otherwise.

**Extended Data Figure 9.**
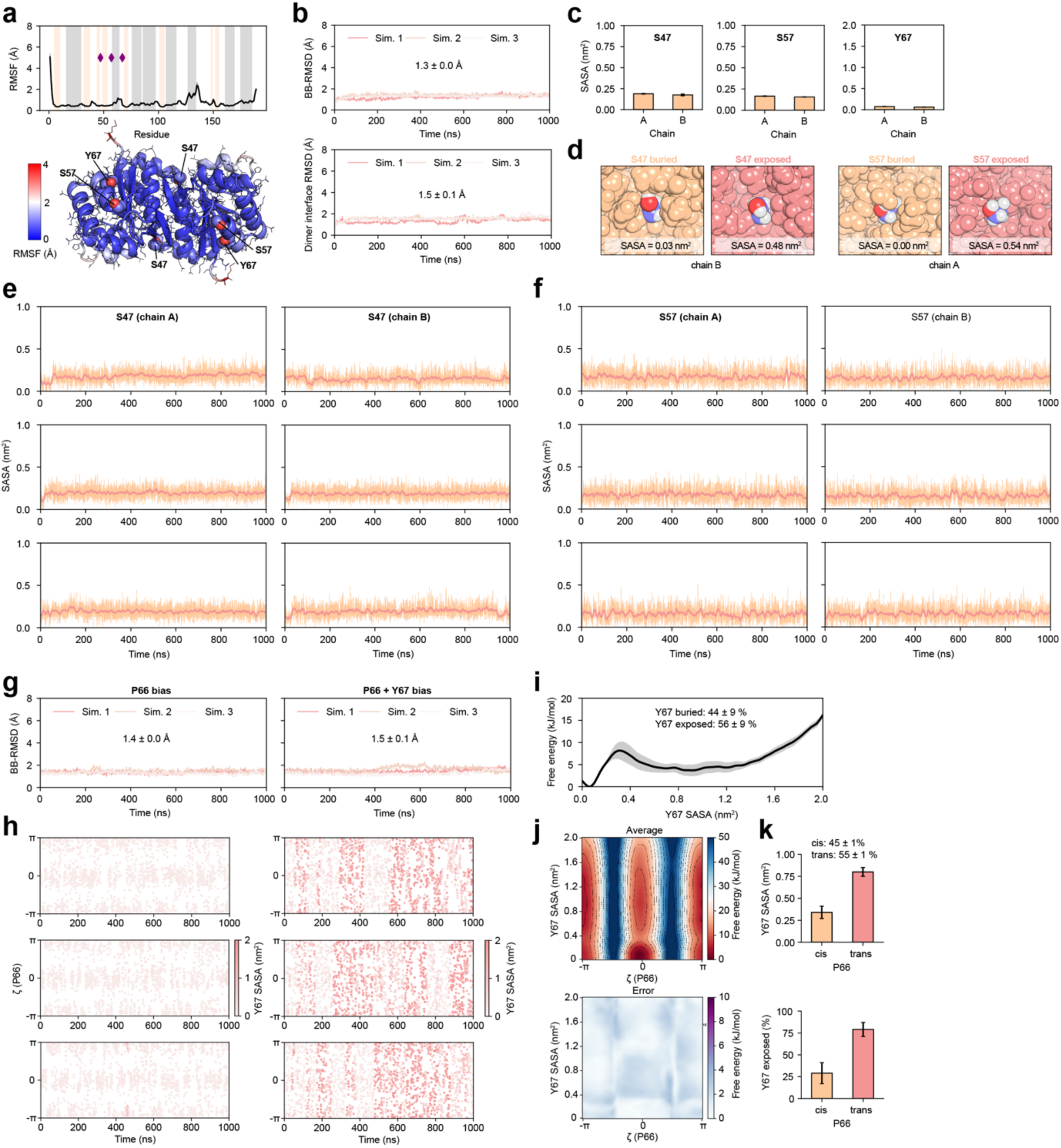
PARK7 sidechain dynamics and local conformational changes induced by proline isomerisation characterised with molecular dynamics simulations. **a,** Backbone dynamics quantified by the RMSF of the Cα atoms along the sequence of PARK7 (average of both chains) and mapped on the structure of the PARK7 dimer (bottom, PDB 2OR3). Grey and orange-shaded regions correspond to α-helices and β-strands, respectively. **b,** Backbone and all-atom dimer interface RMSD of PARK7 dimer simulations calculated with respect to the experimental structure (PDB 2OR3). **c,** Average sidechain SASA of buried PTM sites S47, S57, and Y67 calculated from triplicate unbiased MD simulations (Y67 remains completely buried throughout all unbiased simulations of wild-type PARK7). **d,** Representative structures of buried and partially exposed S47 and S57 sidechain conformations. **e-f,** Sidechain solvent-accessible surface area (SASA) of the buried phosphoserine sites S47 and S47 in both chains as a function of simulation time with a moving average over 0.2 ns. **g,** Backbone RMSD of PARK7 during metadynamics simulations calculated with respect to the experimental structure (PDB 2OR3), biasing proline isomerisation of P66 (left) and proline isomerisation together with the atomic coordination (protein contacts) of the Y67 sidechain (right). **h,** Metadynamics simulations of PARK7 (P66 and P66+Y67 bias as in panel g) showing the improper dihedral angle of P66, ζ, where values near 0 and ± π correspond to the *cis* and *trans* conformations, respectively. The scatter points are coloured according to the sidechain SASA of Y67. **i,** Free energy landscape of PARK7 as a function of Y67 sidechain SASA and the equilibrium populations of the buried and exposed states (separated by a Y67 SASA cut-off of 0.35 nm^2^). **j,** Free energy landscape of PARK7 as a function of P66 isomerisation (ζ) and Y67 sidechain SASA. **k,** Average Y67 sidechain SASA for the P66 *cis* and *trans* conformational ensembles and their populations of the Y67 exposed (SASA > 0.35 nm^2^) state calculated from the metadynamics simulations in panels g-j (P66+Y67 bias). All averaged quantities represent the mean ± SEM from three independent replicates.

**Extended Data Figure 10.**
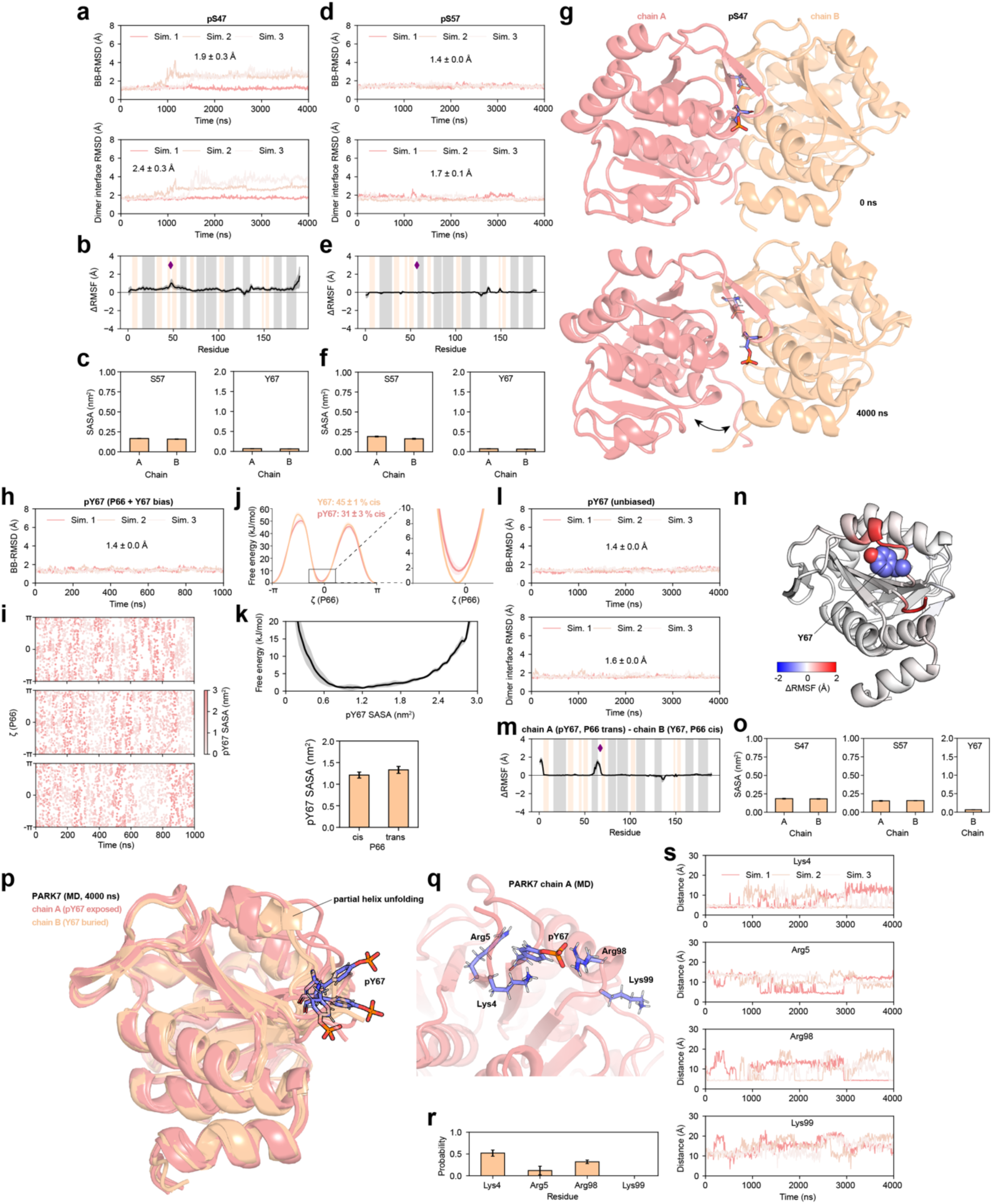
Molecular dynamics simulations of phosphorylated PARK7 variants pS47, pS57, and pY67. **a,d,** Backbone and all-atom dimer interface RMSD of PARK7 pS47 (both chains, panel a) and pS57 (both chains, panel b) dimer simulations calculated with respect to the input structures. **b,e,** Change in backbone dynamics quantified by the RMSF of the Cα atoms along the sequence of PARK7 from unbiased MD simulations of PARK7 pS47 (panel b) and pS57 (panel e) dimer simulations (relative to wild-type and averaged over both chains). Grey and orange-shaded regions correspond to α-helices and β-strands, respectively. **c,f,** Average sidechain SASA of unmodified, buried PTM sites S47, S57, and Y67 calculated from triplicate unbiased MD simulations of pS47 (panel c, both chains phosphorylated at S47) and pS57 (panel f, both chains phosphorylated at S57). These sidechain SASA values remained unchanged relative to the fully unmodified (wild-type) protein (**Extended Data Figure 9c**). **g,** Structure of the PARK7 pS47 dimer at the beginning and end of unbiased simulation 3, showing a loss of dimer contacts (increased RMSD, panel b) and conformational change. The pS47 residues are highlighted with the stick representation. **h,** Backbone RMSD of the PARK7 pY67 dimer (only chain A Y67 modified) during metadynamics simulations calculated with respect to the experimental structure (PDB 2OR3), biasing both proline isomerisation of P66 and Y67-protein contacts. **i,** Metadynamics simulations of PARK7 (P66 and P66+Y67 bias as in panel h) showing the improper dihedral angle of P66, ζ, where values near 0 and ± π correspond to the *cis* and *trans* conformations, respectively. The scatter points are coloured according to the sidechain SASA of pY67. **j,** Free energy landscape of PARK7 dimer wild-type Y67 and pY67 as a function of P66 isomerisation (ζ) and their equilibrium populations of *cis* P66. **k,** Free energy landscape of PARK7 as a function of pY67 sidechain SASA and the average sidechain SASA in the P66 *cis* and *trans* conformational ensembles (bottom), calculated from the metadynamics simulations. **l,** Backbone and all-atom dimer interface RMSD of PARK7 pY67 (only chian A modified) dimer unbiased simulations calculated with respect to the input structures. **m,** Change in backbone dynamics of the modified chain A quantified by the RMSF of the Cα atoms along the sequence of PARK7 from unbiased MD simulations of PARK7 pY67 (panel l) relative to chain B (unmodified, wild-type Y67). **n,** RMSF changes mapped on the chain A structure of PARK7, highlighting the position of Y67. **o,** Average sidechain SASA of unmodified, buried PTM sites S47, S57, and Y67 calculated from triplicate unbiased MD simulations of pY67 (panel l, only chain A phosphorylated at Y67) These sidechain SASA values remained unchanged relative to the fully unmodified (wild-type) protein (**Extended Data Figure 9c**). **p,** Superimposed structures of chain A (pY67) and B (Y67) at the end of the unbiased MD simulations, showing local unfolding near the phosphorylation site in all three simulations. The pY67 sidechain is highlighted in the sticks representation. **q,** A representative structure highlights the most probable interactions of pY67 with Lys4 and Arg98. **r,s,** Average contact probabilities (5 Å contact cut-off) and contact distances between the phosphorylated pY67 and positively charged residues (NZ atom for lysine, CZ atom for arginine).

## SUPPLEMENTARY FIGURE LEGENDS

**Supplementary Figure 1.**
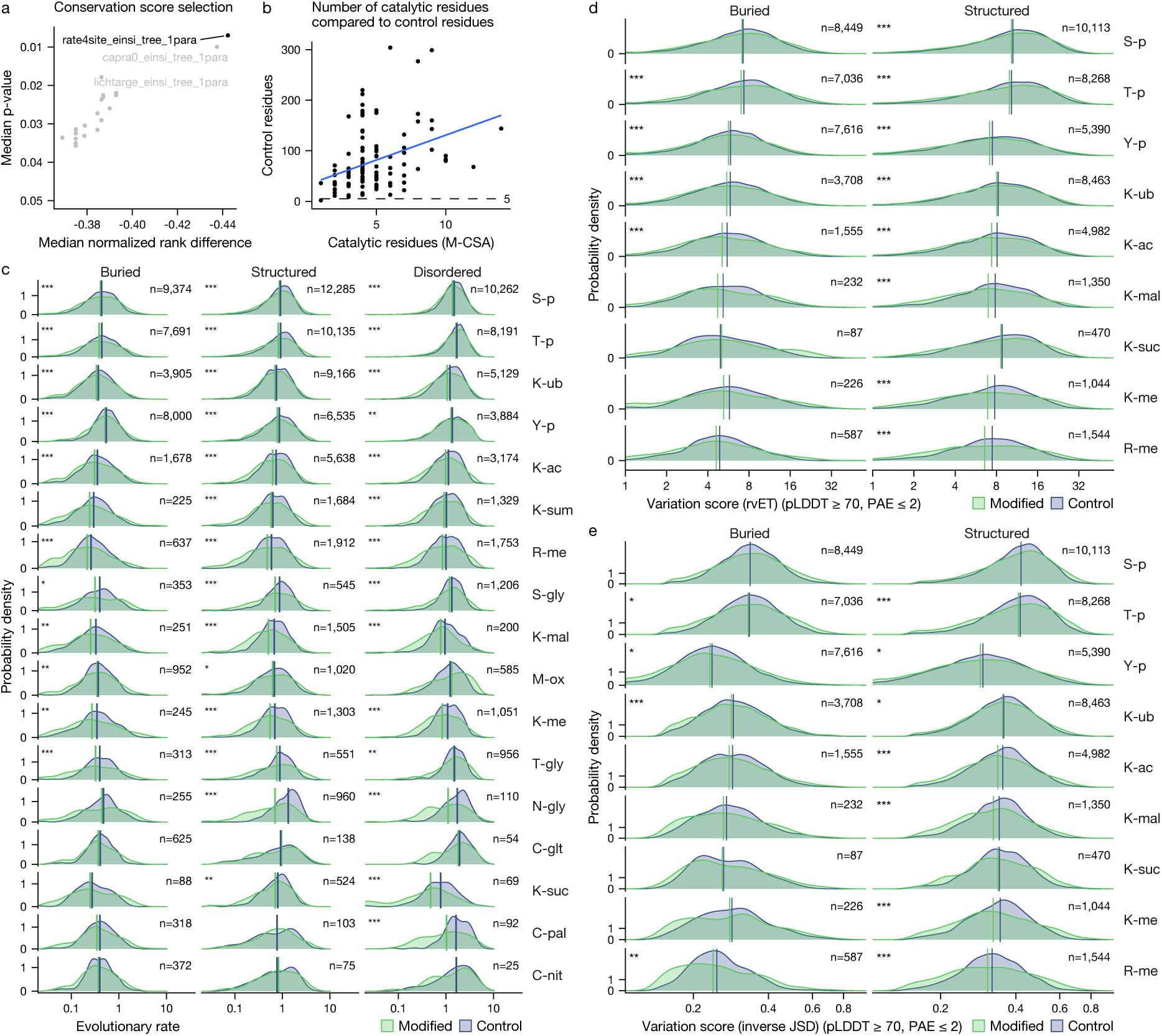
Buried residues that are modified in humans evolve significantly more slowly across species, including when assessed using different conservation scoring methods and alignment and phylogenetic methods. Please refer to **Fig. 2b** for comparison. **a**, Using enzyme catalytic residues from M-CSA to benchmark conservation scores, we observed that Rate4Site, MAFFT E-INS-i, using a species guide tree and including the best-matching orthologues in one-to-many orthology situations led to optimal ability to detect conservation of catalytic residues. This established Rate4Site as the strongest scoring method. Alternatively, using the Jensen-Shannon Divergence (Capra *et al.*) or the real-valued Evolutionary Trace score (Lichtarge *et al.*) led to good results as well. **b**, The number of available control residues is reasonably high for all of the M-CSA enzymes studied. **c**, Unfiltered version of **Fig. 2b** with no structural confidence filter, allowing assessment of disordered modification sites. Buried, structured and disordered sites all showed highly significant conservation compared to control residues of the same structural background and modification type and stratified by protein. **d**, The real-valued Evolutionary Trace score (rvET) likewise reported significant conservation of buried sites, though not for serine phosphorylation. **e**, Jensen-Shannon Divergence likewise showed significant conservation of buried sites, though to a lesser extent.

**Supplementary Figure 2.**
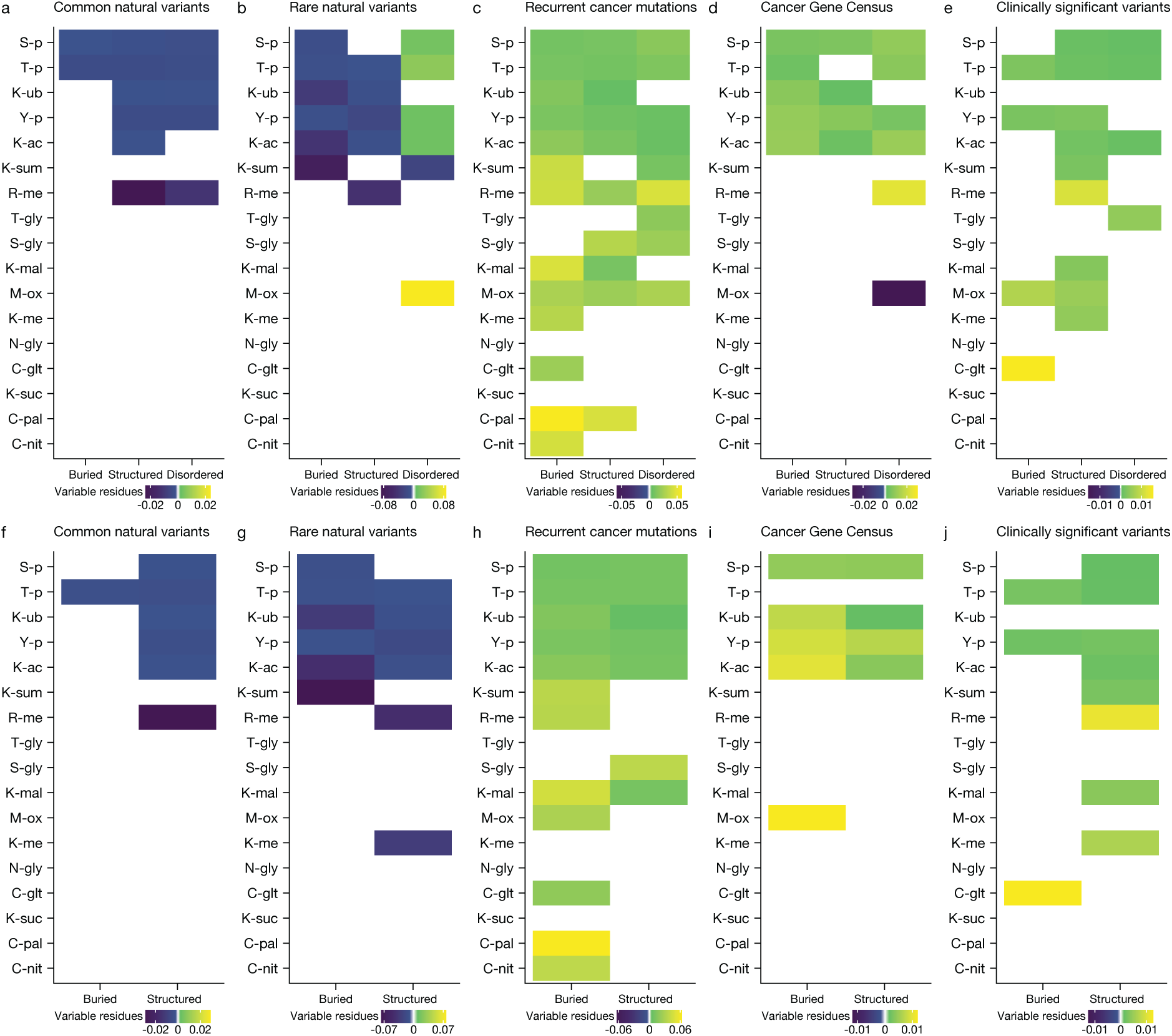
Buried residues that are modified are less often variable in the human population, and are more likely to have disease variants and cancer mutations, shown for additional modification types. Please refer to **Fig. 3a-e** for comparison. **a-e**, Extended versions of **Fig. 3a-e** showing all 17 modification types with more than 1,000 known human sites in our dataset. **f-j**, As a-e, but filtered for structural model confidence using AlphaFold pLDDT ≥ 70 (“confident”).

**Supplementary Figure 3.**
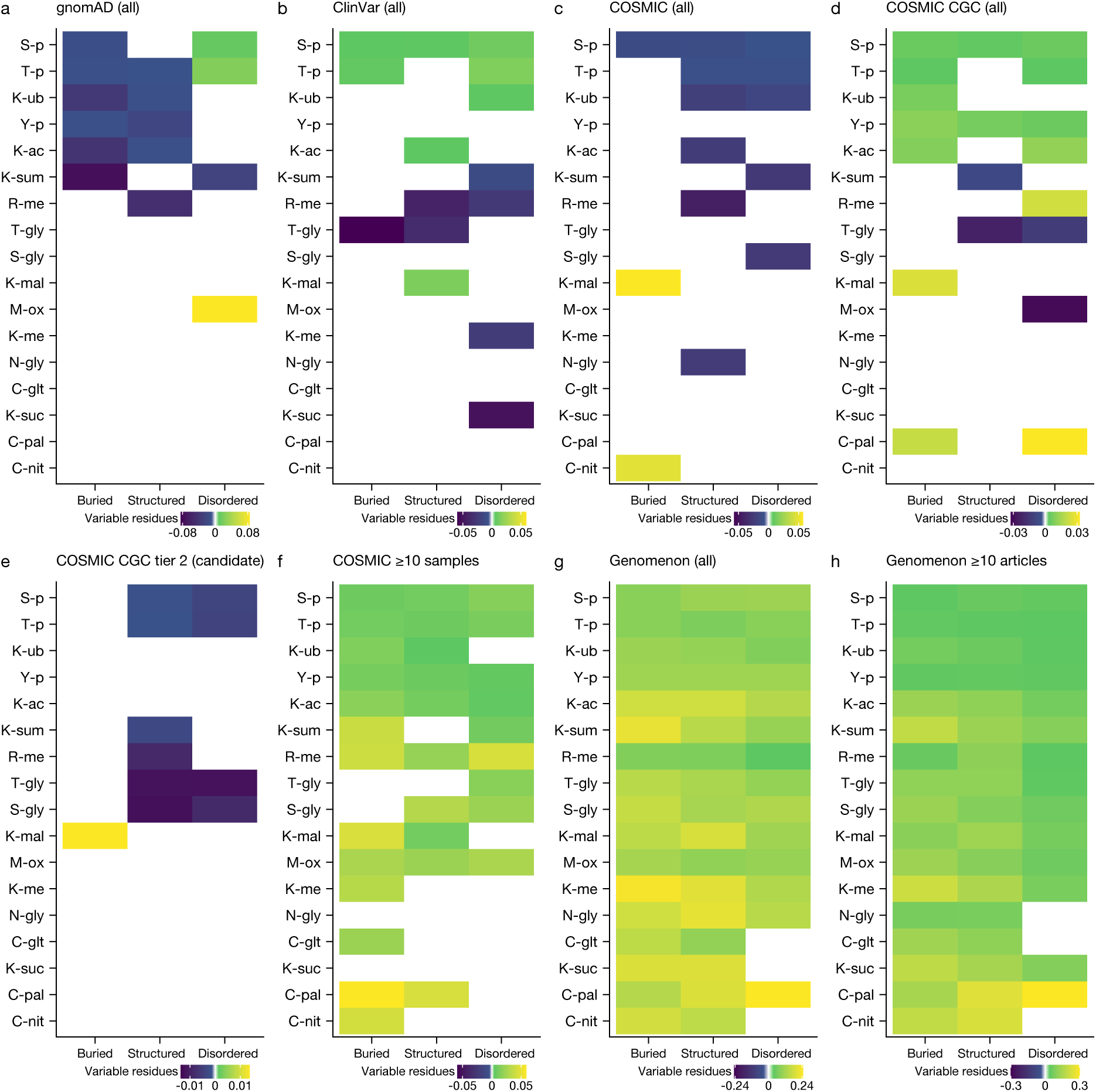
Buried residues that are modified are less often variable in the human population, and are more likely to have disease variants and cancer mutations. Please refer to **Fig. 3a-e** for comparison. Extended versions of **Fig. 3a-e** showing all 17 modification types with more than 1,000 known human sites in our dataset. **a**, Alternative gnomAD v4 plot showing all gnomAD variants (rare and common variants combined). **b**, All ClinVar variants combined, including those not marked as clinically significant. **c**, All COSMIC somatic cancer mutations combined, regardless of recurrence (the number of samples observed in). These appear to behave mainly like natural variants. **d**, COSMIC Cancer Gene Census (CGC) oncogenes and tumour suppressors, including the less confident “tier 2” genes. **e**, COSMIC CGC “tier 2” genes, i.e. candidate genes. **f**, Extended version of **Fig. 3c** showing all 17 modification types. **g**, All variants from the Genomenon Mastermind “Cited Variants Record” (CVR-2). These are any variants reported in the literature, and they appear strongly enriched in modification sites. **h**, Genomenon Mastermind CVR-2 variants that are cited in ≥ 10 articles, showing highly similar enrichment in modification sites, indicating the high interest shown to these in the literature.

**Supplementary Figure 4.**
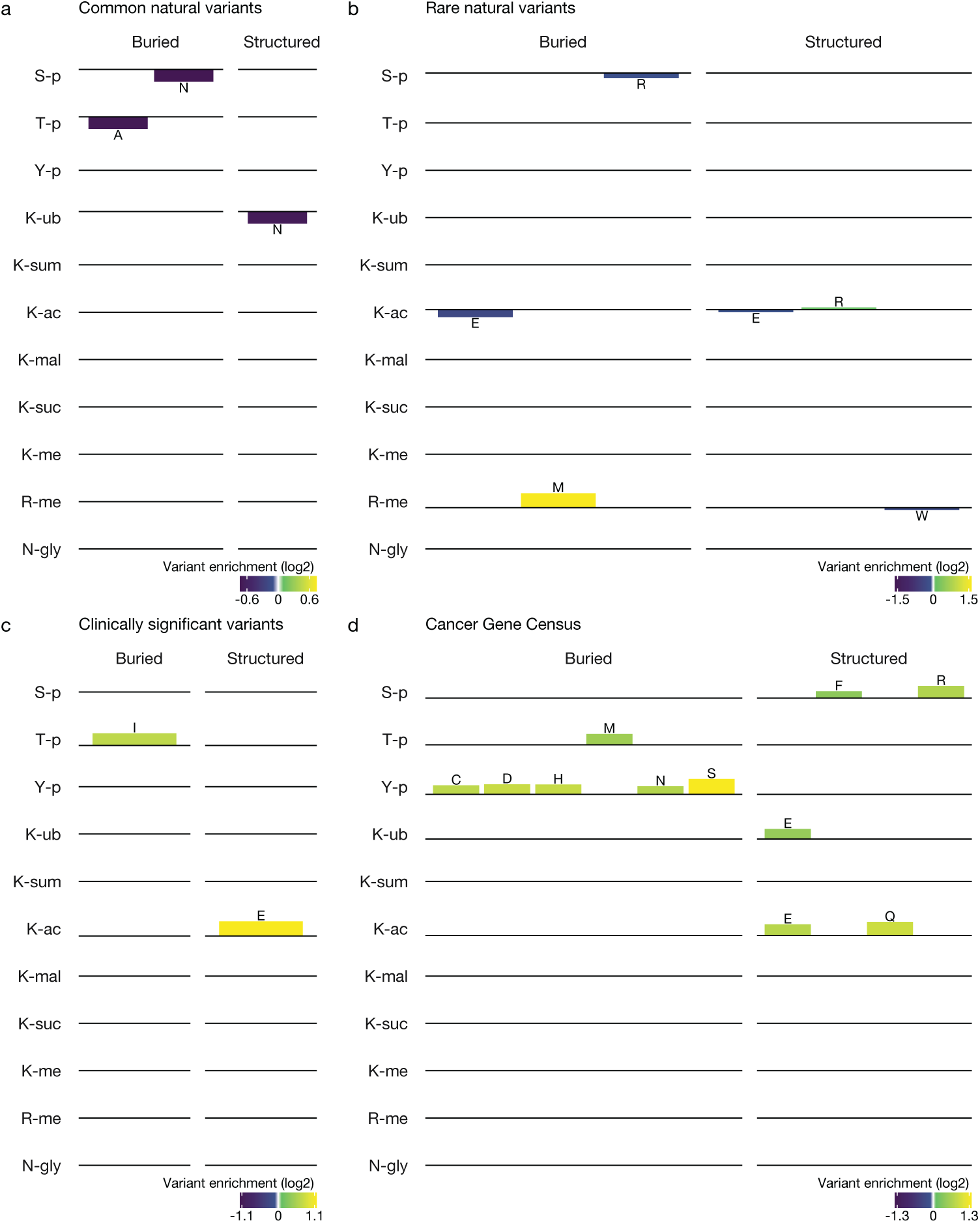
Buried residues that are modified show amino acid variation patterns supporting their functionality. Natural variants showed a depletion in disruptive variants compared to the modified state and an enrichment in variants resembling the modified or unmodified state. Disease variants conversely showed an enrichment in disruptive as well as potentially mimetic amino acids. Bar plots showing statistically significant amino acid enrichments and depletions at modified residues compared to unmodified controls in the same structural category, indicated by a colour scale for each panel, with purple signifying amino acid depletion and green signifying enrichment. Modified and control residues were stringently filtered for structural model confidence using pLDDT ≥ 70 and PAE ≤ 2, as in **Fig. 2b**. Blank panels indicate statistically insignificant differences. Statistical significance was assessed using Fisher’s exact test with FDR correction (P < 0.05). **a**, Common natural variants (more than 1 in 10,000 individuals) in the human population from gnomAD v4, showing e.g. a significant depletion in asparagine (N) for serine phosphorylation sites (S-p). **b**, Rare natural variants (1 in 10,000 individuals or fewer). **c**, Genetic disorder-associated variants marked “clinically significant” in ClinVar, showing an enrichment in variability for buried phosphorylation sites. **d**, Somatic mutations affecting Cancer Gene Census genes from COSMIC, a set of highly validated tumour drivers and suppressors.

**Supplementary Figure 5.**
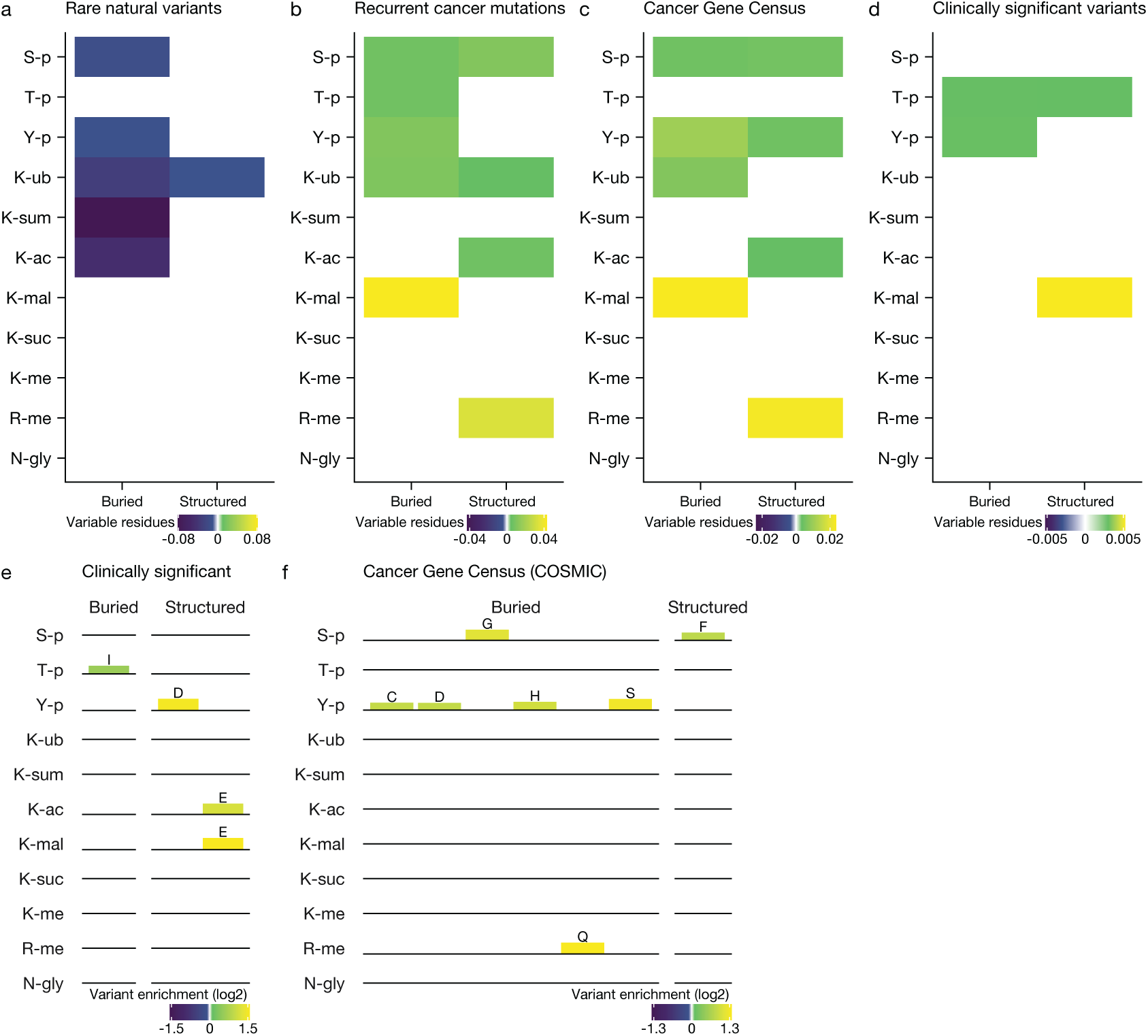
Including when highly stringently structurally filtered, buried residues that are modified are less often variable in the human population, and are more likely to have disease variants and cancer mutations, as well as amino acid variants supporting their functionality. Please refer to **Fig. 3a-h** for comparison. **a-d,** Highly stringently filtered versions of **Fig. 3b-e**, with AlphaFold pLDDT ≥ 90 (“highly confident”) and having at least one contacting residue with PAE ≤ 1 Å. Results for these panels remained similar. No significant depletions (or enrichments) were observed for the equivalent of **Fig. 3a** (common variants), which is hence not shown. **e**, Slightly less stringently filtered version of **Supplementary Fig. 4c** (pLDDT ≥ 70, “confident”, but no PAE filtering), showing significant enrichment of aspartic acid (D) substitutions at structured tyrosine phosphorylation (Y-p) sites. **f**, Highly stringently filtered version of **Fig. 3h**, showing many of the same enrichments, including aspartate (D) at buried tyrosine phosphorylation (Y-p) sites.

**Supplementary Figure 6.**
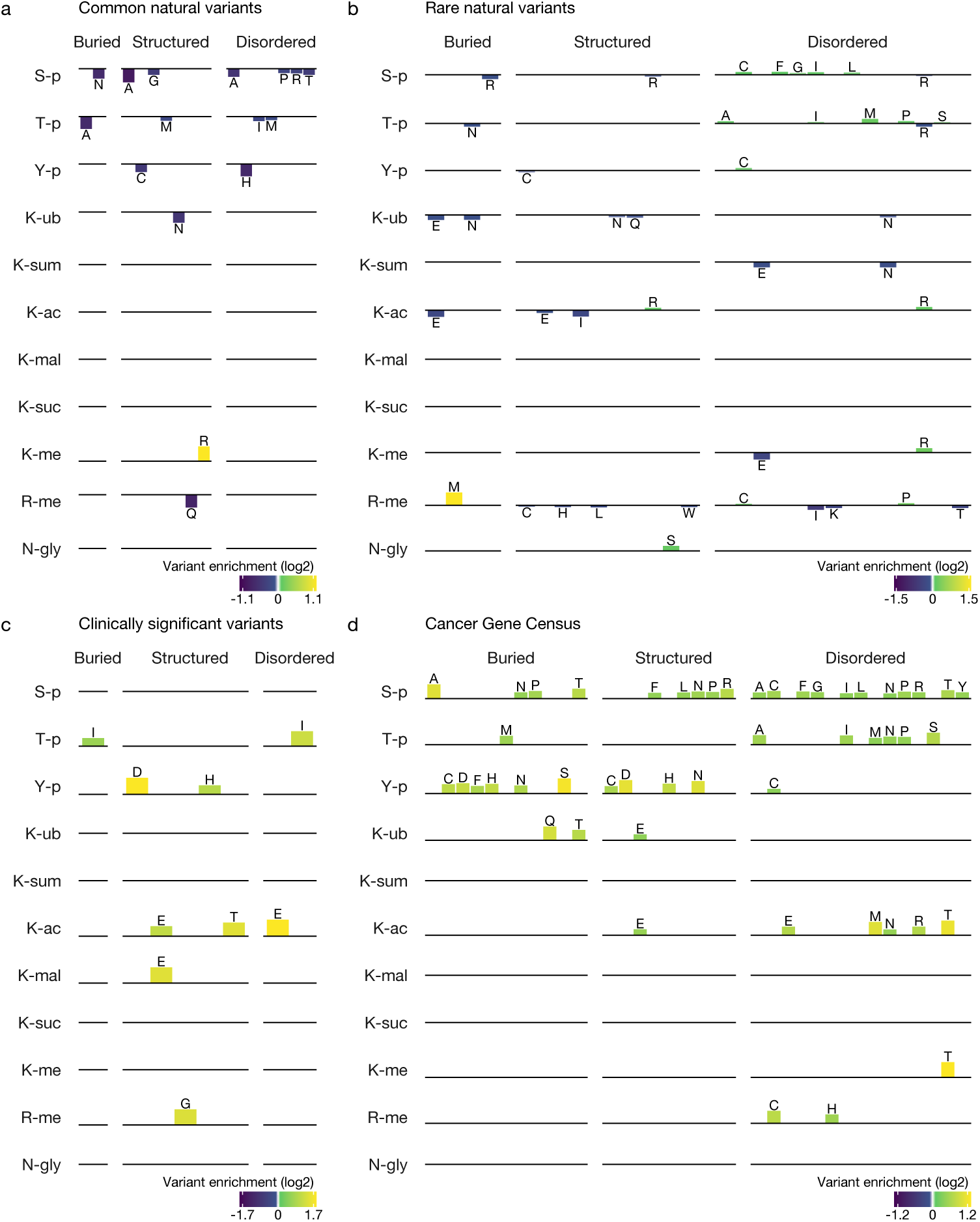
Including when not filtering for enhanced structural model confidence and including for disordered sites, buried residues that are modified are more likely to have amino acid patterns supporting their functionality. Please refer to **Supplementary Fig. 4** for comparison. **a-d,** Amino acid enrichments and depletions at modified residues compared to unmodified controls in the same structural category. No structural filtering by pLDDT or PAE was applied, allowing us to study disordered regions as well, essentially making these extended versions of **Supplementary Fig. 4**. The patterns observed remained similar, with substitutions of lysine (K) to unmodified-like arginine (R) more notable in disordered regions, particularly for gnomAD rare natural variants and Cancer Gene Census genes at lysine acetylation (K-ac) and methylation (K-me) sites. These variants essentially remove a modification site.

**Supplementary Figure 7.**
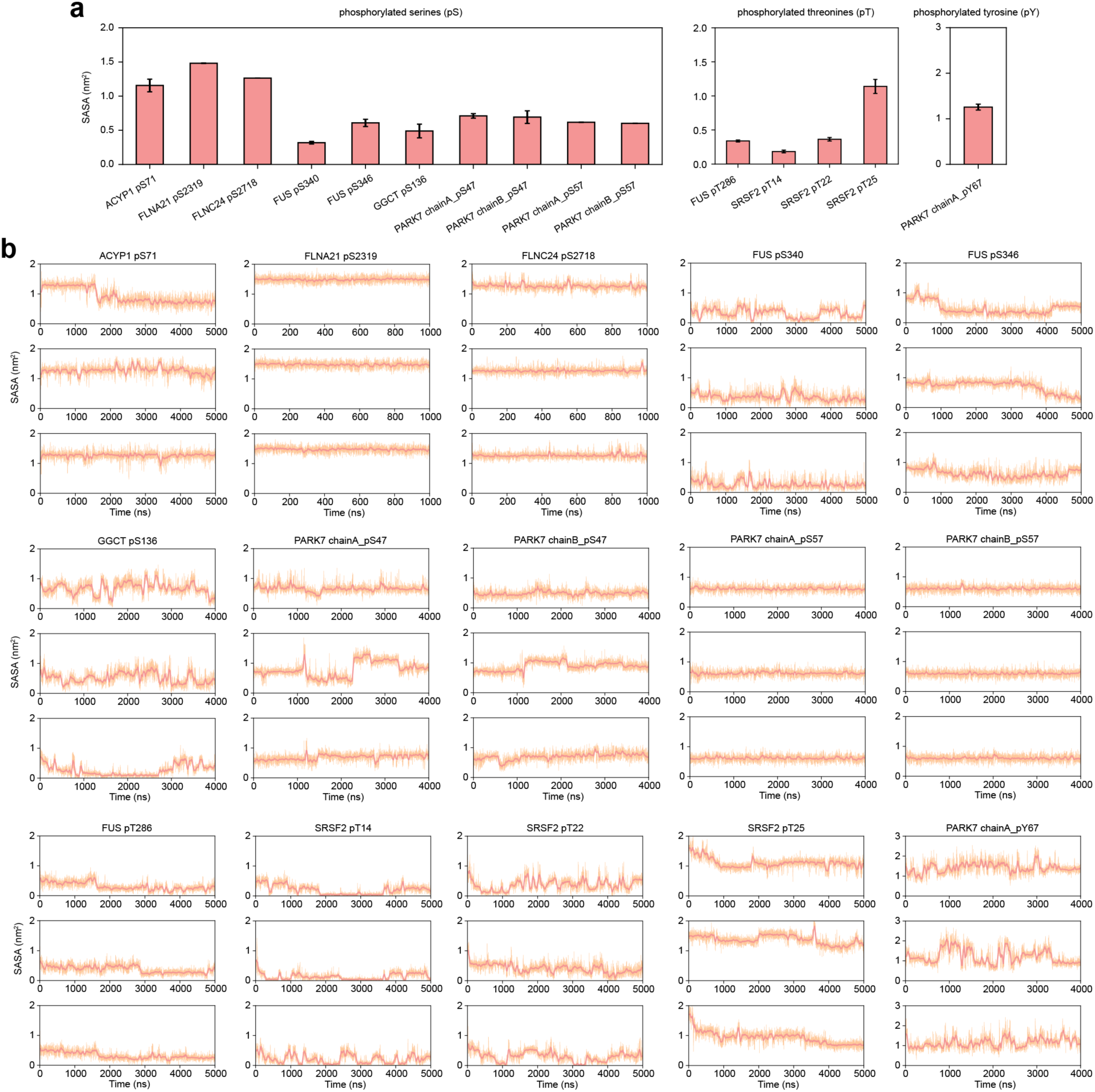
Solvent-accessible surface areas of phosphorylated sidechains calculated from all-atom MD simulations. **a,** Average sidechain solvent-accessible surface area (SASA) of all phosphorylated serine, threonine and tyrosine residues (mean ± SEM from three independent simulations shown in panel b). **b,** Sidechain SASA of the phosphorylated residues as a function of simulation time with a moving average over 1 ns.

**Supplementary Figure 8.**
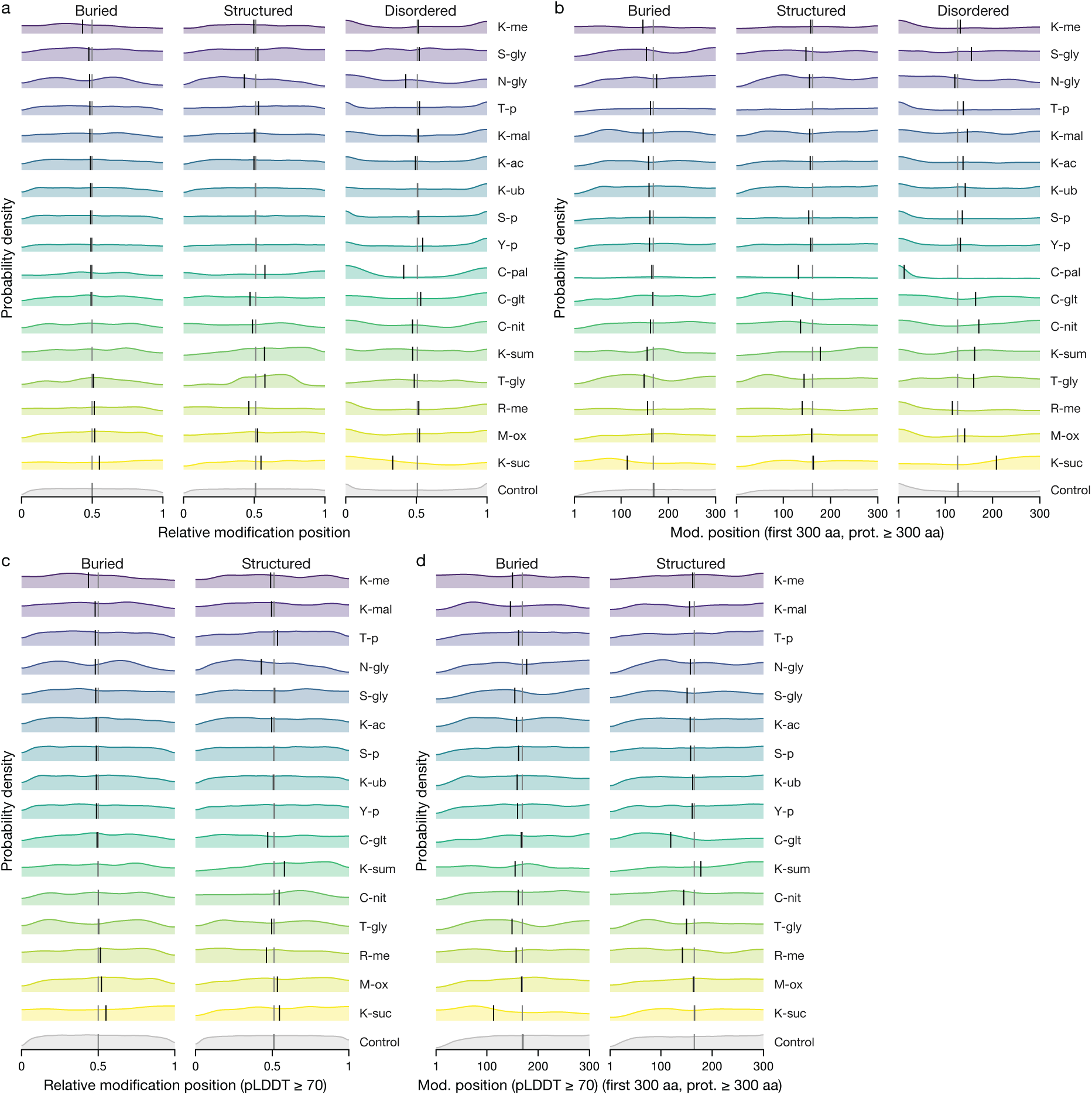
Buried modification sites are significantly shifted towards the N-terminus, suggesting early modification. We found highly significant shifts towards the N-terminus for buried sites, supporting the idea that some of these modifications may occur co-translationally targeting folding intermediates or unfolded nascent chains. **a**, Density plot showing relative positioning of modification sites across proteins compared to structurally equivalent controls, with control amino acid frequency weighted by modification site amino acid frequency for comparability. Buried sites showed a highly significant shift of -0.8% towards the N-terminus (Wilcoxon rank-sum test with continuity correction, two-tailed, n(modified) = 112,451, n(control) = 497,924, W = 2.8e10, P = 4.4e-17). This was robust both when stratifying by protein (P = 9.5e-16) or by modification type and protein (P = 5.7e-07) using asymptotic Wilcoxon-Mann-Whitney tests with a minimum block size of 2. Structured and disordered sites instead showed significant C-terminal shifts by +0.2% (P = 0.0026) and +0.6% (P = 2.2e-20), respectively. Modification types are ordered by median buried position for clarity. Vertical lines show the median in each facet, weighted in case of the control. **b**, Density plot showing the absolute positioning of modication sites for the first 300 residues in proteins of 300 amino acids or longer (70% of proteins, containing 35% of modification sites). We expected these 300 residues to include the first globular domain for most proteins, as supported by the controls, where disorder rises slightly towards residue 300. For disordered sites, the control also reflects the tendency for protein N- and C-termini to be intrinsically disordered^121^. Buried sites showed a highly significant shift of -7 amino acids towards the N-terminus (Wilcoxon rank-sum test with continuity correction, two-tailed, n(modified) = 43,926, n(control) = 180,651, W = 3.8e9, P = 8.4e-59). This was robust both when stratifying by protein (P < 2.2e-16) or by modification type and protein (P < 2.2e-16) using asymptotic Wilcoxon-Mann-Whitney tests with a minimum block size of 2. Structured and disordered sites instead showed significant shifts by -3 (P = 5.6e-15) and +5 (P = 1.5e-54) amino acids, respectively. Modification types are ordered as in panel a. **c**, Filtering by AlphaFold pLDDT score (≥ 70, confident) led to highly similar and slightly stronger results than in panel a. The shift observed for buried sites was -0.9% towards the N-terminus (P = 2.3e-21). **d**, Filtering by AlphaFold pLDDT score (≥ 70, confident) likewise led to highly similar and slightly stronger results than in panel b. The shift observed for buried sites was the same at -7 amino acids towards the N-terminus (P = 1.4e-58).

**Supplementary Table 1 | Amino acid enrichment and depletion for contact and variant analyses.** This table gives detailed statistics for the enrichments and depletions shown in **Fig. 3f** as well as in **Fig. 3g-h**.

(Table included as a separate Excel .xlsx file)

**Supplementary Table 2 | Candidate buried sites identified.** This table gives a detailed overview of their structural context and potential disease relevance. Using structural and evolutionary filtering criteria (see Methods), we identified 4,507 candidate sites in 1,346 proteins. 581 sites are flanked by *cis-* or *trans-*prolines in the -1 or +1 positions, and 2,413 of these proteins have at least one ClinVar variant marked “clinically significant”, indicating their potential disease relevance.

(Table included as a separate Excel .xlsx file)

**Supplementary Table 3.**
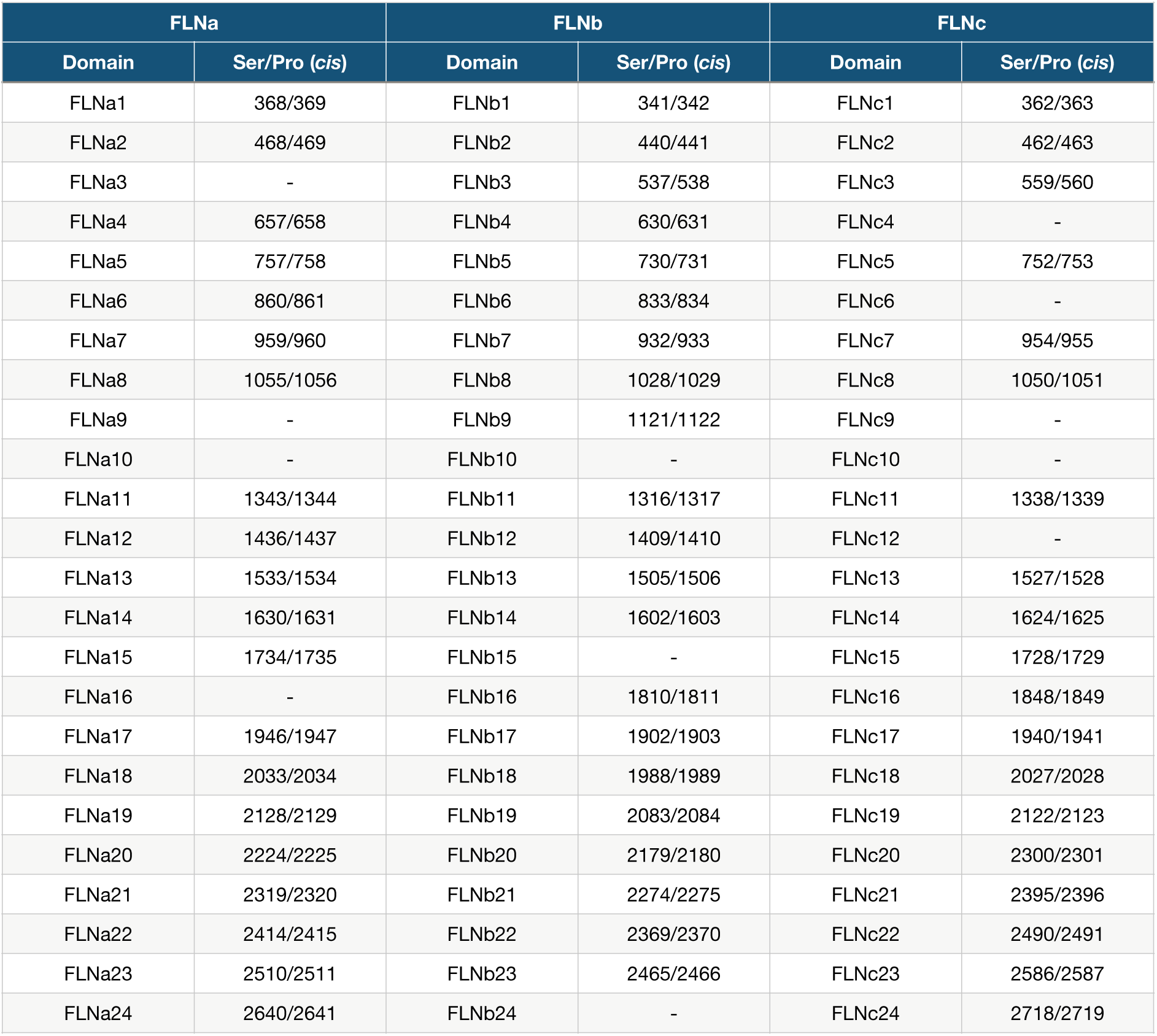
Conservation of buried phosphoserine-proline(*cis*) motif in human filamin Ig-like domains. 60/72 domains contain annotated, buried phosphoserine site upstream of the conserved C-terminal *cis*-proline. The AlphaFold2 models of FLNa/b from the AlphaFold database were used for this analysis, and an AlphaFold3 prediction was generated for FLNc.

| FLNa |  | FLNb |  | FLNc |  |
| --- | --- | --- | --- | --- | --- |
| Domain | Ser/Pro (*cis*) | Domain | Ser/Pro (*cis*) | Domain | Ser/Pro (*cis*) |
| FLNa1 | 368/369 | FLNb1 | 341/342 | FLNc1 | 362/363 |
| FLNa2 | 468/469 | FLNb2 | 440/441 | FLNc2 | 462/463 |
| FLNa3 | - | FLNb3 | 537/538 | FLNc3 | 559/560 |
| FLNa4 | 657/658 | FLNb4 | 630/631 | FLNc4 | - |
| FLNa5 | 757/758 | FLNb5 | 730/731 | FLNc5 | 752/753 |
| FLNa6 | 860/861 | FLNb6 | 833/834 | FLNc6 | - |
| FLNa7 | 959/960 | FLNb7 | 932/933 | FLNc7 | 954/955 |
| FLNa8 | 1055/1056 | FLNb8 | 1028/1029 | FLNc8 | 1050/1051 |
| FLNa9 | - | FLNb9 | 1121/1122 | FLNc9 | - |
| FLNa10 | - | FLNb10 | - | FLNc10 | - |
| FLNa11 | 1343/1344 | FLNb11 | 1316/1317 | FLNc11 | 1338/1339 |
| FLNa12 | 1436/1437 | FLNb12 | 1409/1410 | FLNc12 | - |
| FLNa13 | 1533/1534 | FLNb13 | 1505/1506 | FLNc13 | 1527/1528 |
| FLNa14 | 1630/1631 | FLNb14 | 1602/1603 | FLNc14 | 1624/1625 |
| FLNa15 | 1734/1735 | FLNb15 | - | FLNc15 | 1728/1729 |
| FLNa16 | - | FLNb16 | 1810/1811 | FLNc16 | 1848/1849 |
| FLNa17 | 1946/1947 | FLNb17 | 1902/1903 | FLNc17 | 1940/1941 |
| FLNa18 | 2033/2034 | FLNb18 | 1988/1989 | FLNc18 | 2027/2028 |
| FLNa19 | 2128/2129 | FLNb19 | 2083/2084 | FLNc19 | 2122/2123 |
| FLNa20 | 2224/2225 | FLNb20 | 2179/2180 | FLNc20 | 2300/2301 |
| FLNa21 | 2319/2320 | FLNb21 | 2274/2275 | FLNc21 | 2395/2396 |
| FLNa22 | 2414/2415 | FLNb22 | 2369/2370 | FLNc22 | 2490/2491 |
| FLNa23 | 2510/2511 | FLNb23 | 2465/2466 | FLNc23 | 2586/2587 |
| FLNa24 | 2640/2641 | FLNb24 | - | FLNc24 | 2718/2719 |

**Supplementary Table 4.**
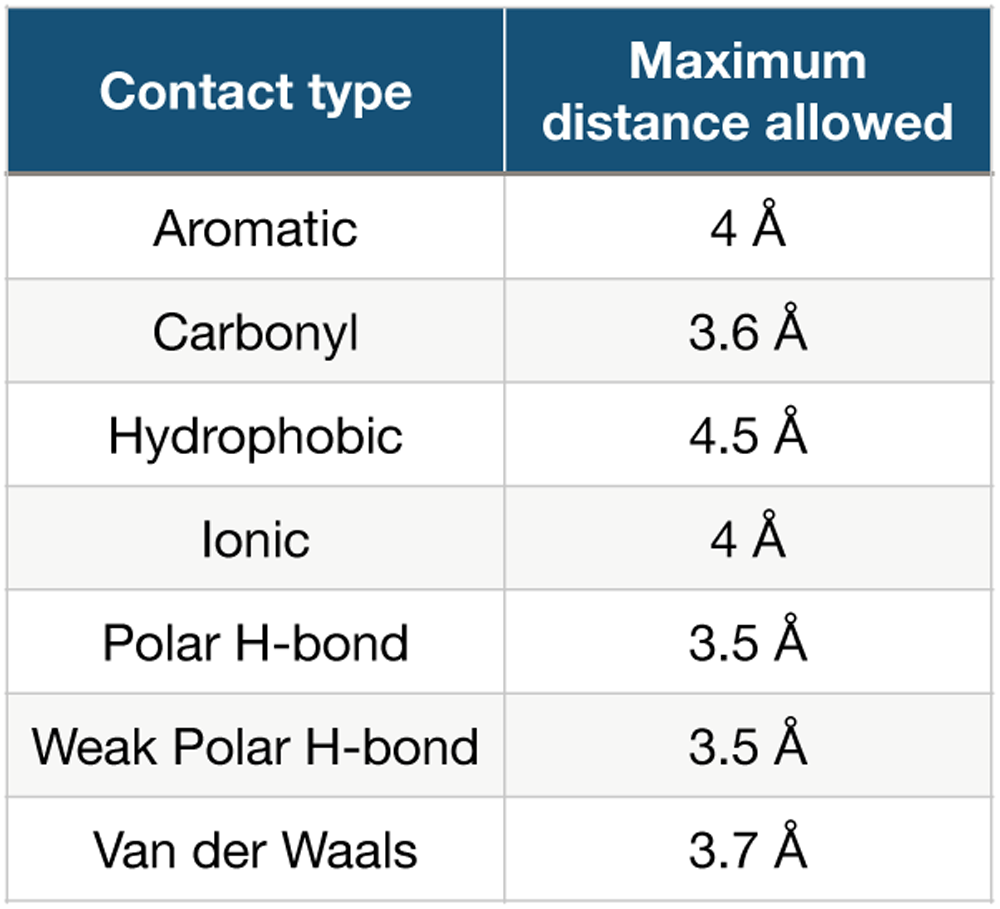
Distance cut-offs used for contact detection. This table shows the default distance cut-offs used by Lahuta for different types of contacts.

| Contact type | Maximum distance allowed |
| --- | --- |
| Aromatic | 4 Å |
| Carbonyl | 3.6 Å |
| Hydrophobic | 4.5 Å |
| Ionic | 4 Å |
| Polar H-bond | 3.5 Å |
| Weak Polar H-bond | 3.5 Å |
| Van der Waals | 3.7 Å |

**Supplementary Table 5.**
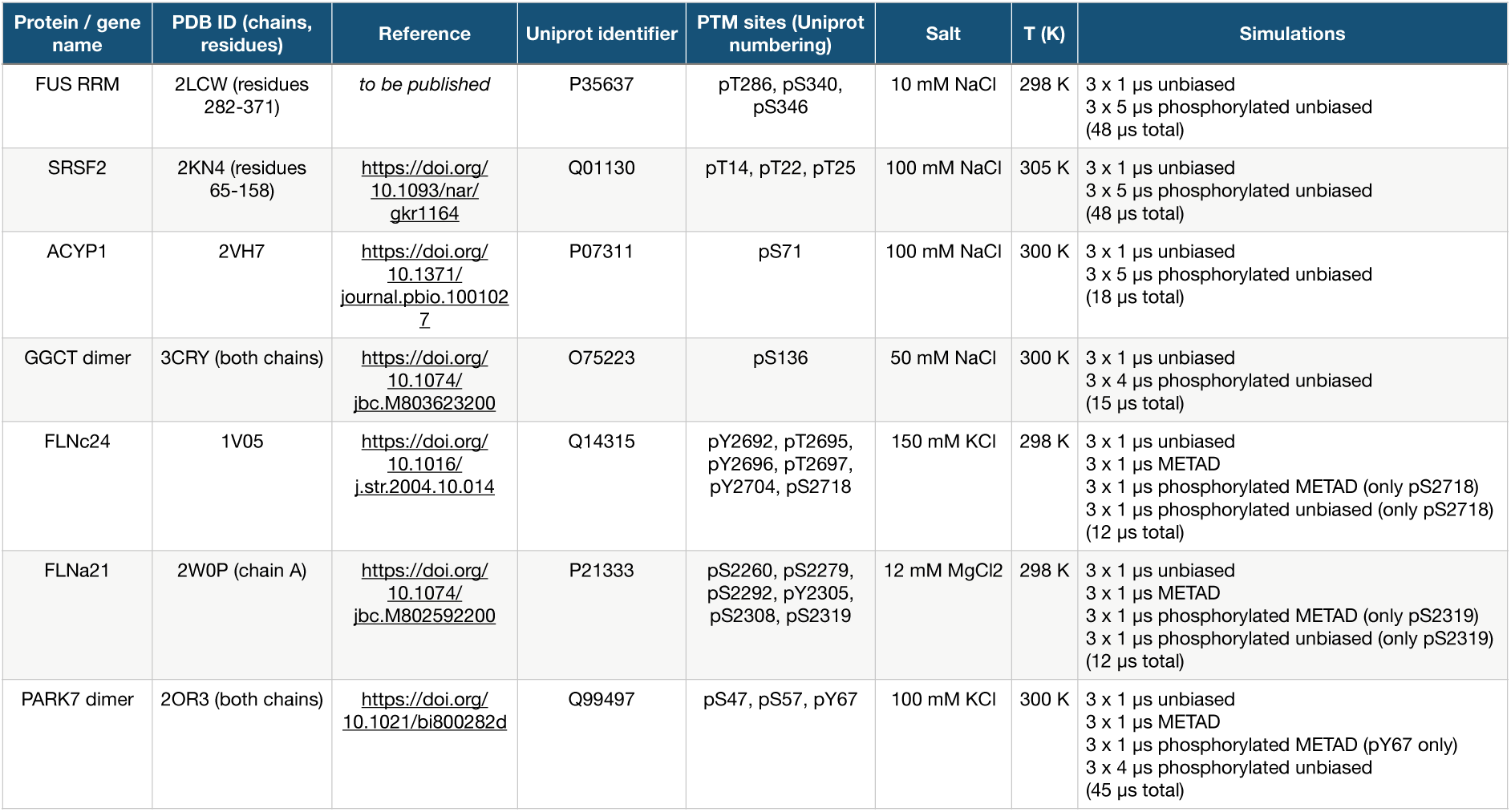
Summary of MD simulations performed in this study. A total dataset of 198 μs of MD simulations was accumulated in this work. “METAD” refers to metadynamics simulations.

## MAIN REFERENCES

1 Phillips, A. H. & Kriwacki, R. W. Intrinsic protein disorder and protein modifications in the processing of biological signals. Curr Opin Struct Biol 60, 1–6 (2020). 10.1016/j.sbi.2019.09.003

2 Vellosillo, P. & Minguez, P. A global map of associations between types of protein posttranslational modifications and human genetic diseases. iScience 24, 102917 (2021). 10.1016/j.isci.2021.102917

3 Keenan, E. K., Zachman, D. K. & Hirschey, M. D. Discovering the landscape of protein modifications. Mol Cell 81, 1868–1878 (2021). 10.1016/j.molcel.2021.03.015

4 Bradley, D. The evolution of post-translational modifications. Curr Opin Genet Dev 76, 101956 (2022). 10.1016/j.gde.2022.101956

5 Deribe, Y. L., Pawson, T. & Dikic, I. Post-translational modifications in signal integration. Nat Struct Mol Biol 17, 666–672 (2010). 10.1038/nsmb.1842

6 Jimenez, J. L., Hegemann, B., Hutchins, J. R., Peters, J. M. & Durbin, R. A systematic comparative and structural analysis of protein phosphorylation sites based on the mtcPTM database. Genome Biol 8, R90 (2007). 10.1186/gb-2007-8-5-r90

7 Chan, S. H. S. et al. Structures of protein folding intermediates on the ribosome. bioRxiv, 2025.2004.2007.647236 (2025). 10.1101/2025.04.07.647236

8 Mann, M. & Jensen, O. N. Proteomic analysis of post-translational modifications. Nat Biotechnol 21, 255–261 (2003). 10.1038/nbt0303-255

9 Walsh, C. T., Garneau-Tsodikova, S. & Gatto, G. J., Jr. Protein posttranslational modifications: the chemistry of proteome diversifications. Angew Chem Int Ed Engl 44, 7342–7372 (2005). 10.1002/anie.200501023

10 Tompa, P., Davey, N. E., Gibson, T. J. & Babu, M. M. A million peptide motifs for the molecular biologist. Mol Cell 55, 161–169 (2014). 10.1016/j.molcel.2014.05.032

11 Bah, A. & Forman-Kay, J. D. Modulation of Intrinsically Disordered Protein Function by Post-translational Modifications. J Biol Chem 291, 6696–6705 (2016). 10.1074/jbc.R115.695056

12 Csizmok, V., Follis, A. V., Kriwacki, R. W. & Forman-Kay, J. D. Dynamic Protein Interaction Networks and New Structural Paradigms in Signaling. Chem Rev 116, 6424–6462 (2016). 10.1021/acs.chemrev.5b00548

13 Jakob, U., Kriwacki, R. & Uversky, V. N. Conditionally and transiently disordered proteins: awakening cryptic disorder to regulate protein function. Chem Rev 114, 6779–6805 (2014). 10.1021/cr400459c

14 Mitrea, D. M. & Kriwacki, R. W. Regulated unfolding of proteins in signaling. FEBS Lett 587, 1081–1088 (2013). 10.1016/j.febslet.2013.02.024

15 Henriques, J. & Lindorff-Larsen, K. Protein Dynamics Enables Phosphorylation of Buried Residues in Cdk2/Cyclin-A-Bound p27. Biophys J 119, 2010–2018 (2020). 10.1016/j.bpj.2020.06.040

16 Orioli, S., Henning Hansen, C. G. & Lindorff-Larsen, K. Transient exposure of a buried phosphorylation site in an autoinhibited protein. Biophys J 121, 91–101 (2022). 10.1016/j.bpj.2021.11.2890

17 Keshwani, M. M. et al. Cotranslational cis-phosphorylation of the COOH-terminal tail is a key priming step in the maturation of cAMP-dependent protein kinase. Proc Natl Acad Sci U S A 109, E1221–1229 (2012). 10.1073/pnas.1202741109

18 Oh, W. J. et al. mTORC2 can associate with ribosomes to promote cotranslational phosphorylation and stability of nascent Akt polypeptide. EMBO J 29, 3939–3951 (2010). 10.1038/emboj.2010.271

19 Mocan, I. et al. Protein phosphorylation corrects the folding defect of the neuroblastoma (S120G) mutant of human nucleoside diphosphate kinase A/Nm23-H1. Biochem J 403, 149–156 (2007). 10.1042/BJ20061141

20 Hovland, R., Doskeland, A. P., Eikhom, T. S., Robaye, B. & Doskeland, S. O. cAMP induces co-translational modification of proteins in IPC-81 cells. Biochem J 342 (**Pt 2**), 369–377 (1999).

21 Kamacioglu, A., Tuncbag, N. & Ozlu, N. Structural analysis of mammalian protein phosphorylation at a proteome level. Structure 29, 1219–1229 e1213 (2021). 10.1016/j.str.2021.06.008

22 Bludau, I. et al. The structural context of posttranslational modifications at a proteome-wide scale. PLoS Biol 20, e3001636 (2022). 10.1371/journal.pbio.3001636

23 Nissley, D. A. et al. Universal protein misfolding intermediates can bypass the proteostasis network and remain soluble and less functional. Nat Commun 13, 3081 (2022). 10.1038/s41467-022-30548-5

24 Ochoa, D. et al. The functional landscape of the human phosphoproteome. Nat Biotechnol 38, 365–373 (2020). 10.1038/s41587-019-0344-3

25 Varadi, M. et al. AlphaFold Protein Structure Database in 2024: providing structure coverage for over 214 million protein sequences. Nucleic Acids Res 52, D368–D375 (2024). 10.1093/nar/gkad1011

26 Lang, B. & Babu, M. M. AlphaSync is an enhanced AlphaFold structure database synchronized with UniProt. bioRxiv (2025). 10.1101/2025.03.12.642845

27 Kumar, A. et al. Phosphorylation-induced unfolding regulates p19(INK4d) during the human cell cycle. Proc Natl Acad Sci U S A 115, 3344–3349 (2018). 10.1073/pnas.1719774115

28 Xiao, Y. et al. Phosphorylation releases constraints to domain motion in ERK2. Proc Natl Acad Sci U S A 111, 2506–2511 (2014). 10.1073/pnas.1318899111

29 Tien, M. Z., Meyer, A. G., Sydykova, D. K., Spielman, S. J. & Wilke, C. O. Maximum allowed solvent accessibilites of residues in proteins. PLoS One 8, e80635 (2013). 10.1371/journal.pone.0080635

30 Levy, E. D. A simple definition of structural regions in proteins and its use in analyzing interface evolution. J Mol Biol 403, 660–670 (2010). 10.1016/j.jmb.2010.09.028

31 Akdel, M. et al. A structural biology community assessment of AlphaFold2 applications. Nat Struct Mol Biol 29, 1056–1067 (2022). 10.1038/s41594-022-00849-w

32 UniProt, C. UniProt: the Universal Protein Knowledgebase in 2025. Nucleic Acids Res 53, D609–D617 (2025). 10.1093/nar/gkae1010

33 Hornbeck, P. V. et al. 15 years of PhosphoSitePlus(R): integrating post-translationally modified sites, disease variants and isoforms. Nucleic Acids Res 47, D433–D441 (2019). 10.1093/nar/gky1159

34 Li, Z. et al. dbPTM in 2022: an updated database for exploring regulatory networks and functional associations of protein post-translational modifications. Nucleic Acids Res 50, D471–D479 (2022). 10.1093/nar/gkab1017

35 Johansson, F. & Toh, H. A comparative study of conservation and variation scores. BMC Bioinformatics 11, 388 (2010). 10.1186/1471-2105-11-388

36 Ribeiro, A. J. M. et al. Mechanism and Catalytic Site Atlas (M-CSA): a database of enzyme reaction mechanisms and active sites. Nucleic Acids Res 46, D618–D623 (2018). 10.1093/nar/gkx1012

37 Amoutzias, G. D. et al. Posttranslational regulation impacts the fate of duplicated genes. Proc Natl Acad Sci U S A 107, 2967–2971 (2010). 10.1073/pnas.0911603107

38 Chen, S. et al. A genomic mutational constraint map using variation in 76,156 human genomes. Nature 625, 92–100 (2024). 10.1038/s41586-023-06045-0

39 Sondka, Z. et al. COSMIC: a curated database of somatic variants and clinical data for cancer. Nucleic Acids Res 52, D1210–D1217 (2024). 10.1093/nar/gkad986

40 Sondka, Z. et al. The COSMIC Cancer Gene Census: describing genetic dysfunction across all human cancers. Nat Rev Cancer 18, 696–705 (2018). 10.1038/s41568-018-0060-1

41 Landrum, M. J. et al. ClinVar: updates to support classifications of both germline and somatic variants. Nucleic Acids Res 53, D1313–D1321 (2025). 10.1093/nar/gkae1090

42 Weirich, S. & Jeltsch, A. Limited choice of natural amino acids as mimetics restricts design of protein lysine methylation studies. Nat Commun 14, 4097 (2023). 10.1038/s41467-023-39777-8

43 Dall’Agnese, A. et al. Proteolethargy is a pathogenic mechanism in chronic disease. Cell 188, 207–221 e230 (2025). 10.1016/j.cell.2024.10.051

44 Wang, X. & Hayes, J. J. Acetylation mimics within individual core histone tail domains indicate distinct roles in regulating the stability of higher-order chromatin structure. Mol Cell Biol 28, 227–236 (2008). 10.1128/MCB.01245-07

45 Pearlman, S. M., Serber, Z. & Ferrell, J. E., Jr. A mechanism for the evolution of phosphorylation sites. Cell 147, 934–946 (2011). 10.1016/j.cell.2011.08.052

46 Li, P., Martins, I. R., Amarasinghe, G. K. & Rosen, M. K. Internal dynamics control activation and activity of the autoinhibited Vav DH domain. Nat Struct Mol Biol 15, 613–618 (2008). 10.1038/nsmb.1428

47 Yu, B. et al. Structural and energetic mechanisms of cooperative autoinhibition and activation of Vav1. Cell 140, 246–256 (2010). 10.1016/j.cell.2009.12.033

48 Tsytlonok, M. et al. Dynamic anticipation by Cdk2/Cyclin A-bound p27 mediates signal integration in cell cycle regulation. Nat Commun 10, 1676 (2019). 10.1038/s41467-019-09446-w

49 Reimer, U. et al. Side-chain effects on peptidyl-prolyl cis/trans isomerisation. J Mol Biol 279, 449–460 (1998). 10.1006/jmbi.1998.1770

50 Waudby, C. A. et al. Systematic mapping of free energy landscapes of a growing filamin domain during biosynthesis. Proc Natl Acad Sci U S A 115, 9744–9749 (2018). 10.1073/pnas.1716252115

51 Arenas, A. et al. Lysine acetylation regulates the RNA binding, subcellular localization and inclusion formation of FUS. Hum Mol Genet 29, 2684–2697 (2020). 10.1093/hmg/ddaa159

52 Phelan, M. M. et al. The structure and selectivity of the SR protein SRSF2 RRM domain with RNA. Nucleic Acids Res 40, 3232–3244 (2012). 10.1093/nar/gkr1164

53 Witt, A. C. et al. Cysteine pKa depression by a protonated glutamic acid in human DJ-1. Biochemistry 47, 7430–7440 (2008). 10.1021/bi800282d

54 Loughlin, F. E. et al. The Solution Structure of FUS Bound to RNA Reveals a Bipartite Mode of RNA Recognition with Both Sequence and Shape Specificity. Mol Cell 73, 490–504 e496 (2019). 10.1016/j.molcel.2018.11.012

55 Kumar, M. et al. ELM-the Eukaryotic Linear Motif resource-2024 update. Nucleic Acids Res 52, D442–D455 (2024). 10.1093/nar/gkad1058

56 Daubner, G. M., Clery, A., Jayne, S., Stevenin, J. & Allain, F. H. A syn-anti conformational difference allows SRSF2 to recognize guanines and cytosines equally well. EMBO J 31, 162–174 (2012). 10.1038/emboj.2011.367

57 Oakley, A. J. et al. The identification and structural characterization of C7orf24 as gamma-glutamyl cyclotransferase. An essential enzyme in the gamma-glutamyl cycle. J Biol Chem 283, 22031–22042 (2008). 10.1074/jbc.M803623200

58 Chan, S. H. S. et al. The ribosome stabilizes partially folded intermediates of a nascent multi-domain protein. Nat Chem 14, 1165–1173 (2022). 10.1038/s41557-022-01004-0

59 Streit, J. O. et al. The ribosome lowers the entropic penalty of protein folding. Nature 633, 232–239 (2024). 10.1038/s41586-024-07784-4

60 Lad, Y. et al. Structural basis of the migfilin-filamin interaction and competition with integrin beta tails. J Biol Chem 283, 35154–35163 (2008). 10.1074/jbc.M802592200

61 Pudas, R., Kiema, T. R., Butler, P. J., Stewart, M. & Ylanne, J. Structural basis for vertebrate filamin dimerization. Structure 13, 111–119 (2005). 10.1016/j.str.2004.10.014

62 McCoy, A. J., Fucini, P., Noegel, A. A. & Stewart, M. Structural basis for dimerization of the Dictyostelium gelation factor (ABP120) rod. Nat Struct Biol 6, 836–841 (1999). 10.1038/12296

63 Szeto, S. G. Y., Williams, E. C., Rudner, A. D. & Lee, J. M. Phosphorylation of filamin A by Cdk1 regulates filamin A localization and daughter cell separation. Exp Cell Res 330, 248–266 (2015). 10.1016/j.yexcr.2014.10.024

64 Han, J. H., Batey, S., Nickson, A. A., Teichmann, S. A. & Clarke, J. The folding and evolution of multidomain proteins. Nat Rev Mol Cell Biol 8, 319–330 (2007). 10.1038/nrm2144

65 Streit, J. O. et al. Long-range electrostatic forces govern how proteins fold on the ribosome. bioRxiv (2025). 10.1101/2025.02.10.637539

66 Holtkamp, W. et al. Cotranslational protein folding on the ribosome monitored in real time. Science 350, 1104–1107 (2015). 10.1126/science.aad0344

67 Wales, T. E. et al. Resolving chaperone-assisted protein folding on the ribosome at the peptide level. Nat Struct Mol Biol (2024). 10.1038/s41594-024-01355-x

68 Wang, F., Durfee, L. A. & Huibregtse, J. M. A cotranslational ubiquitination pathway for quality control of misfolded proteins. Mol Cell 50, 368–378 (2013). 10.1016/j.molcel.2013.03.009

69 He, Y., Chen, Y., Alexander, P. A., Bryan, P. N. & Orban, J. Mutational tipping points for switching protein folds and functions. Structure 20, 283–291 (2012). 10.1016/j.str.2011.11.018

70 Xia, F. et al. Raf activation is regulated by tyrosine 510 phosphorylation in Drosophila. PLoS Biol 6, e128 (2008). 10.1371/journal.pbio.0060128

71 Chen, Z. & Cole, P. A. Synthetic approaches to protein phosphorylation. Curr Opin Chem Biol 28, 115–122 (2015). 10.1016/j.cbpa.2015.07.001

72 Stateva, S. R. et al. Characterization of phospho-(tyrosine)-mimetic calmodulin mutants. PLoS One 10, e0120798 (2015). 10.1371/journal.pone.0120798

73 Duarte, M. L. et al. Protein folding creates structure-based, noncontiguous consensus phosphorylation motifs recognized by kinases. Sci Signal 7, ra105 (2014). 10.1126/scisignal.2005412

74 Lu, K. P., Finn, G., Lee, T. H. & Nicholson, L. K. Prolyl cis-trans isomerization as a molecular timer. Nat Chem Biol 3, 619–629 (2007). 10.1038/nchembio.2007.35

75 Schiene-Fischer, C. Multidomain Peptidyl Prolyl cis/trans Isomerases. Biochim Biophys Acta 1850, 2005–2016 (2015). 10.1016/j.bbagen.2014.11.012

76 Yaffe, M. B. et al. Sequence-specific and phosphorylation-dependent proline isomerization: a potential mitotic regulatory mechanism. Science 278, 1957–1960 (1997). 10.1126/science.278.5345.1957

77 Follis, A. V. et al. Pin1-Induced Proline Isomerization in Cytosolic p53 Mediates BAX Activation and Apoptosis. Mol Cell 59, 677–684 (2015). 10.1016/j.molcel.2015.06.029

78 Brichkina, A. et al. Proline isomerisation as a novel regulatory mechanism for p38MAPK activation and functions. Cell Death Differ 23, 1592–1601 (2016). 10.1038/cdd.2016.45

79 Cassaignau, A. M. E., Cabrita, L. D. & Christodoulou, J. How Does the Ribosome Fold the Proteome? Annu Rev Biochem 89, 389–415 (2020). 10.1146/annurev-biochem-062917-012226

80 Kaiser, C. M., Goldman, D. H., Chodera, J. D., Tinoco, I., Jr. & Bustamante, C. The ribosome modulates nascent protein folding. Science 334, 1723–1727 (2011). 10.1126/science.1209740

81 Kramer, G., Boehringer, D., Ban, N. & Bukau, B. The ribosome as a platform for co-translational processing, folding and targeting of newly synthesized proteins. Nat Struct Mol Biol 16, 589–597 (2009). 10.1038/nsmb.1614

82 Hyvonen, M. & Saraste, M. Structure of the PH domain and Btk motif from Bruton’s tyrosine kinase: molecular explanations for X-linked agammaglobulinaemia. EMBO J 16, 3396–3404 (1997). 10.1093/emboj/16.12.3396

83 Lomakin, I. B., Dmitriev, S. E. & Steitz, T. A. Crystal structure of the DENR-MCT-1 complex revealed zinc-binding site essential for heterodimer formation. Proc Natl Acad Sci U S A 116, 528–533 (2019). 10.1073/pnas.1809688116

84 Li, Z. et al. dbPTM in 2022: an updated database for exploring regulatory networks and functional associations of protein post-translational modifications. Nucleic Acids Res 50, D471–D479 (2022). 10.1093/nar/gkab1017

85 Lee, J. M., Hammaren, H. M., Savitski, M. M. & Baek, S. H. Control of protein stability by post-translational modifications. Nat Commun 14, 201 (2023). 10.1038/s41467-023-35795-8

86 Doulias, P. T. et al. Structural profiling of endogenous S-nitrosocysteine residues reveals unique features that accommodate diverse mechanisms for protein S-nitrosylation. Proc Natl Acad Sci U S A 107, 16958–16963 (2010). 10.1073/pnas.1008036107

87 Grek, C. L., Zhang, J., Manevich, Y., Townsend, D. M. & Tew, K. D. Causes and consequences of cysteine S-glutathionylation. J Biol Chem 288, 26497–26504 (2013). 10.1074/jbc.R113.461368

88 Kim, H. Y. & Gladyshev, V. N. Methionine sulfoxide reductases: selenoprotein forms and roles in antioxidant protein repair in mammals. Biochem J 407, 321–329 (2007). 10.1042/BJ20070929

89 Lee, B. C. et al. MsrB1 and MICALs regulate actin assembly and macrophage function via reversible stereoselective methionine oxidation. Mol Cell 51, 397–404 (2013). 10.1016/j.molcel.2013.06.019

90 Ko, P. J. & Dixon, S. J. Protein palmitoylation and cancer. EMBO Rep 19 (2018). 10.15252/embr.201846666

91 Lipman, D. J. & Pearson, W. R. Rapid and sensitive protein similarity searches. Science 227, 1435–1441 (1985). 10.1126/science.2983426

92 Joosten, R. P. et al. A series of PDB related databases for everyday needs. Nucleic Acids Res 39, D411–419 (2011). 10.1093/nar/gkq1105

93 Kabsch, W. & Sander, C. Dictionary of protein secondary structure: pattern recognition of hydrogen-bonded and geometrical features. Biopolymers 22, 2577– 2637 (1983). 10.1002/bip.360221211

94 Hamelryck, T. & Manderick, B. PDB file parser and structure class implemented in Python. Bioinformatics 19, 2308–2310 (2003). 10.1093/bioinformatics/btg299

95 Harrison, P. W. et al. Ensembl 2024. Nucleic Acids Res 52, D891–D899 (2024). 10.1093/nar/gkad1049

96 Diss, G., Freschi, L. & Landry, C. R. Where do phosphosites come from and where do they go after gene duplication? Int J Evol Biol 2012, 843167 (2012). 10.1155/2012/843167

97 Katoh, K. & Standley, D. M. MAFFT multiple sequence alignment software version 7: improvements in performance and usability. Mol Biol Evol 30, 772–780 (2013). 10.1093/molbev/mst010

98 Vos, R. A., Caravas, J., Hartmann, K., Jensen, M. A. & Miller, C. BIO::Phylo-phyloinformatic analysis using perl. BMC Bioinformatics 12, 63 (2011). 10.1186/1471-2105-12-63

99 Suyama, M., Torrents, D. & Bork, P. PAL2NAL: robust conversion of protein sequence alignments into the corresponding codon alignments. Nucleic Acids Res 34, W609–612 (2006). 10.1093/nar/gkl315

100 Guindon, S. et al. New algorithms and methods to estimate maximum-likelihood phylogenies: assessing the performance of PhyML 3.0. Syst Biol 59, 307–321 (2010). 10.1093/sysbio/syq010

101 Ruan, J. et al. TreeFam: 2008 Update. Nucleic Acids Res 36, D735–740 (2008). 10.1093/nar/gkm1005

102 Capra, J. A. & Singh, M. Predicting functionally important residues from sequence conservation. Bioinformatics 23, 1875–1882 (2007). 10.1093/bioinformatics/btm270

103 Pupko, T., Bell, R. E., Mayrose, I., Glaser, F. & Ben-Tal, N. Rate4Site: an algorithmic tool for the identification of functional regions in proteins by surface mapping of evolutionary determinants within their homologues. Bioinformatics 18 **Suppl 1**, S71– 77 (2002). 10.1093/bioinformatics/18.suppl_1.s71

104 Mihalek, I., Res, I. & Lichtarge, O. A family of evolution-entropy hybrid methods for ranking protein residues by importance. J Mol Biol 336, 1265–1282 (2004). 10.1016/j.jmb.2003.12.078

105 McLaren, W. et al. The Ensembl Variant Effect Predictor. Genome Biol 17, 122 (2016). 10.1186/s13059-016-0974-4

106 Chunn, L. M. et al. Mastermind: A Comprehensive Genomic Association Search Engine for Empirical Evidence Curation and Genetic Variant Interpretation. Front Genet 11, 577152 (2020). 10.3389/fgene.2020.577152

107 ww, P. D. B. c. Protein Data Bank: the single global archive for 3D macromolecular structure data. Nucleic Acids Res 47, D520–D528 (2019). 10.1093/nar/gky949

108 Abraham, M. J. et al. GROMACS: High performance molecular simulations through multi-level parallelism from laptops to supercomputers. SoftwareX 1-2, 19–25 (2015). 10.1016/j.softx.2015.06.001

109 Huang, J. et al. CHARMM36m: an improved force field for folded and intrinsically disordered proteins. Nat Methods 14, 71–73 (2017). 10.1038/nmeth.4067

110 Hess, B., Bekker, H., Berendsen, H. J. C. & Fraaije, J. G. E. M. LINCS: A linear constraint solver for molecular simulations. J Comput Chem 18, 1463–1472 (1997). 10.1002/(SICI)1096-987X(199709)18:12%3C1463::AID-JCC4%3E3.0.CO;2-H

111 Darden, T., York, D. & Pedersen, L. Particle Mesh Ewald - an N.Log(N) Method for Ewald Sums in Large Systems. J Chem Phys 98, 10089–10092 (1993). 10.1063/1.464397

112 Bussi, G., Donadio, D. & Parrinello, M. Canonical sampling through velocity rescaling. J Chem Phys 126, 014101 (2007). 10.1063/1.2408420

113 Berendsen, H. J. C., Postma, J. P. M., Vangunsteren, W. F., Dinola, A. & Haak, J. R. Molecular-Dynamics with Coupling to an External Bath. J Chem Phys 81, 3684–3690 (1984). 10.1063/1.448118

114 Parrinello, M. & Rahman, A. Polymorphic Transitions in Single-Crystals - a New Molecular-Dynamics Method. J Appl Phys 52, 7182–7190 (1981). 10.1063/1.328693

115 Michaud-Agrawal, N., Denning, E. J., Woolf, T. B. & Beckstein, O. MDAnalysis: A Toolkit for the Analysis of Molecular Dynamics Simulations. J Comput Chem 32, 2319–2327 (2011). 10.1002/jcc.21787

116 Barducci, A., Bussi, G. & Parrinello, M. Well-tempered metadynamics: a smoothly converging and tunable free-energy method. Phys Rev Lett 100, 020603 (2008). 10.1103/PhysRevLett.100.020603

117 Consortium, P. Promoting transparency and reproducibility in enhanced molecular simulations. Nat Methods 16, 670–673 (2019). 10.1038/s41592-019-0506-8

118 Tribello, G. A., Bonomi, M., Branduardi, D., Camilloni, C. & Bussi, G. PLUMED 2: New feathers for an old bird. Comput Phys Commun 185, 604–613 (2014). 10.1016/j.cpc.2013.09.018

119 Melis, C., Bussi, G., Lummis, S. C. & Molteni, C. Trans-cis switching mechanisms in proline analogues and their relevance for the gating of the 5-HT3 receptor. J Phys Chem B 113, 12148–12153 (2009). 10.1021/jp9046962

120 Tiwary, P. & Parrinello, M. A time-independent free energy estimator for metadynamics. J Phys Chem B 119, 736–742 (2015). 10.1021/jp504920s

121 Romero, P. et al. Sequence complexity of disordered protein. Proteins 42, 38–48 (2001). 10.1002/1097-0134(20010101)42:1<38::aid-prot50>3.0.co;2-3

